# The peptide LyeTx I mnΔK induces transcriptomic reprogramming in a novel Multidrug-resistant *Acinetobacter baumannii*

**DOI:** 10.1101/2025.09.29.679197

**Authors:** Frederico Gabriel de Carvalho Oliveira, Katharina de Oliveira Barros, Dáfne de Oliveira Vianei, Julia Raspante Martins, Jessica Caroline Duarte, Daniela de Laet Souza, Izabela Mamede, Rennan Garcias Moreira, Ranato Santana de Aguiar, Filipe Alex da Silva, Vera Lúcia dos Santos, Alessandro de Mello Varani, Thiago Mafra Batista, Carlos Renato Machado, Carlos Augusto Rosa, Maria Elena de Lima, Glória Regina Franco

**Affiliations:** Departamento de Bioquímica e Imunologia, ICB, Universidade Federal de Minas Gerais, Belo Horizonte, Minas Gerais, Brazil; Departamento de Microbiologia, ICB, Universidade Federal de Minas Gerais, Belo Horizonte, Minas Gerais, Brazil; Programa Interunidades de Pós-Graduação em Bioinformática da UFMG, ICB, Universidade Federal de Minas Gerais, Belo Horizonte, Minas Gerais, Brazil; Departamento de Genética, Ecologia e Evolução, ICB, Universidade Federal de Minas Gerais, Belo Horizonte, Minas Gerais, Brazil; Departamento de Tecnologia, Faculdade de Ciências Agrárias e Veterinárias de Jaboticabal, Universidade Estadual Paulista Júlio de Mesquita Filho, Jaboticabal, São Paulo, Brazil; Instituto Nacional da Mata Atlântica, Santa Teresa, Espírito Santo, Brazil; Faculdade de Saúde Santa Casa de Belo Horizonte, Belo Horizonte, Minas Gerais, Brazil

**Keywords:** Antimicrobial peptides (AMP), Gene Expression, Transcriptomics, Comparative Genomics, Bacteria, Antibiotic Resistance, Meropenem, LyeTx I mnΔK, *Acinetobacter baumannii*

## Abstract

*Acinetobacter baumannii* is a critical pathogen in healthcare-associated infections, and treatment is challenging due to the emergence of multidrug-resistant strains. Antimicrobial peptides, such as LyeTx I mnΔK, a synthetic peptide derived of a toxin from the spider *Lycosa erythrognatha*, represent a promising alternative due to their broad-spectrum activity and synergistic potential with antibiotics like meropenem. This study aimed to compare the genomes of several *A. baumannii* strains, including a novel multidrug-resistant *A. baumannii* isolate (AC37), and to evaluate the antimicrobial effects of LyeTx I mnΔK-alone and in combination with meropenem-through transcriptomic analysis. Genome assembly and annotation of AC37 revealed 31 antibiotic resistance genes, and phylogenetic analysis comprising 123 *A. baumannii* genomes, including the reference strain, identified three unique resistant genes in the AC37 strain. Mobilome analysis showed 13 genes associated with mobile genetic elements, including two of the unique genes, highlighting horizontal gene transfer events. Transcriptomic profiling revealed that treatment with LyeTx I mnΔK peptide alone induced several differentially expressed genes, including two efflux pump operons. Additionally, pathways related to protein synthesis, export, and secretion were activated, indicating a broader cellular response to the peptide. The treatment with LyeTx I mnΔK in combination with meropenem disrupted oxidative phosphorylation, further revealing the metabolic plasticity of the bacterial response to external stresses. This study characterizes a new *A. baumannii* isolate and provides new insights into the bacterial response to a potential novel therapeutic molecule.

## INTRODUCTION

*Acinetobacter baumannii* is an opportunistic pathogen primarily associated with nosocomial infections (1–3). Susceptibility to infection by this bacterium is significantly increased in patients who have undergone recent surgical procedures or have concomitant infections. This microorganism exhibits tropism for multiple tissues, including the skin, urinary tract, wounds, and the central nervous system. Pulmonary colonization, in particular, represents the most severe infectious manifestation, often leading to severe pneumonia and a high mortality rate (4, 5).

The treatment of bacterial infections presents a significant challenge, largely due to the ability of bacteria to evolve or acquire diverse resistance mechanisms, including drug-degrading or modifying enzymes, efflux pumps, and alterations in cellular permeability (6, 7). Approximately 45% of *A. baumannii* isolates are classified as multidrug-resistant (MDR), representing one of the highest rates among the ESKAPE group (*Enterococcus faecium, Staphylococcus aureus, Klebsiella pneumoniae, A. baumannii, Pseudomonas aeruginosa*, and *Enterobacter* species) (8). Inappropriate and excessive use of carbapenems, combined with delayed diagnosis, has driven the emergence of carbapenem-resistant *A. baumannii* (CRAB) (9). Currently, a class of polymyxin polypeptides, such as colistin, remains the primary therapeutic option for treating *A. baumannii* infections. Despite their efficacy, these drugs are associated with adverse effects - particularly nephrotoxicity and neurotoxicity - which limit their clinical use and highlight the need for therapeutic alternatives (10, 11).

The Centers for Disease Control and Prevention (CDC) classifies the search for new therapeutic alternatives to treat CRABs as a top public health priority. Among the most promising alternatives are Antimicrobial Peptides (AMPs). AMPs are composed of a net positive charge and amphiphilic structure, which enables electrostatic interactions with negatively charged components of microbial cell walls and membranes (12–14). The LyeTx I mnΔK, a synthetic and modified peptide of 16 aminoacid residues, derived of a toxin from the spider *Lycosa erythrognatha*, exhibited broad-spectrum antimicrobial activity against gram-positive and gram-negative bacteria as well as yeasts (15). Importantly, this peptide also inhibited biofilm formation, presented low cytotoxicity and hemolytic activity, and acted synergistically with meropenem, a clinically relevant antibiotic (16).

In this study, we performed a comprehensive genomic analysis, identifying three unique genes associated with resistance mechanisms in a novel multidrug-resistant *A. baumannii* strain - AC37 - isolated from a central venous catheter of a patient at Santa Casa in Belo Horizonte, MG, Brazil (16). Furthermore, we evaluated the antimicrobial efficacy of the LyeTx I mnΔK peptide, both alone and in combination with meropenem, by analyzing the transcriptome of treated bacteria. This analysis revealed significant alterations in key metabolic pathways, including protein synthesis, fatty acid catabolism and the electron transport chain, suggesting potential mechanisms of action for the peptide. To contextualize our results, we compared them with BioProjects of *A. baumannii* treated exclusively with meropenem and found that the antibiotics treatment suppresses fatty acid biosynthesis. These findings enhance our understanding of CRAB resistance and provide promising insights to the development of a novel therapeutic strategy.

## RESULTS

### Genomic clustering places AC37 among both resistant and susceptible isolates, yet divergent from the *A. baumannii* reference strain

Sequencing of the novel *A. baumannii* AC37 strain generated 5,117,651 paired-end reads with an average length of 301 base pairs, resulting in approximately 750-fold genome coverage. The assembled genome size was 3,843,875 bp, with an N50 of 206,736 bp and an L50 of 8 contigs **(Table S1)**.

Genome visualization and plasmid profiling detected one single plasmid of 8,731 bp, identified as NZ_CP131945.1, as deposited at National Center for Biotechnology Information (NCBI) **(Fig. S1)**. Although plasmids often carry antibiotic resistance genes in bacterial species, this plasmid did not contain any such genes. Instead, it harbored the BrnT/BrnA toxin-antitoxin system, a genetic element known to modulate bacterial physiology under stress conditions. These genes are organized within an operon that encodes a cytoplasmic toxic protein (BrnTp) and its cognate antitoxin (BrnAp). BrnTp functions as a ribonuclease and forms a 2:2 heterotetrameric complex with BrnAp. This tetrameric complex can autoregulate its own expression by binding to its promoter region via BrnAp, leading to transcriptional repression (17, 18).

To elucidate the placement of the novel AC37 strain, a maximum likelihood phylogenetic analysis was conducted with 123 *A. baumannii* genomes obtained from the NCBI Genomes database **(Fig. 1, Table S2)**. The AC37 strain clustered within a clade comprising diverse isolates, including both multidrug-resistant and non-multidrug-resistant strains. Interestingly, the AC37 strain exhibited significant phylogenetic divergence from the reference strain ATCC 19606, underscoring the critical importance of comprehensive genomic characterization for discerning the evolutionary dynamics and phenotypic variability of emerging *A. baumannii* strains.

**Figure 1.**
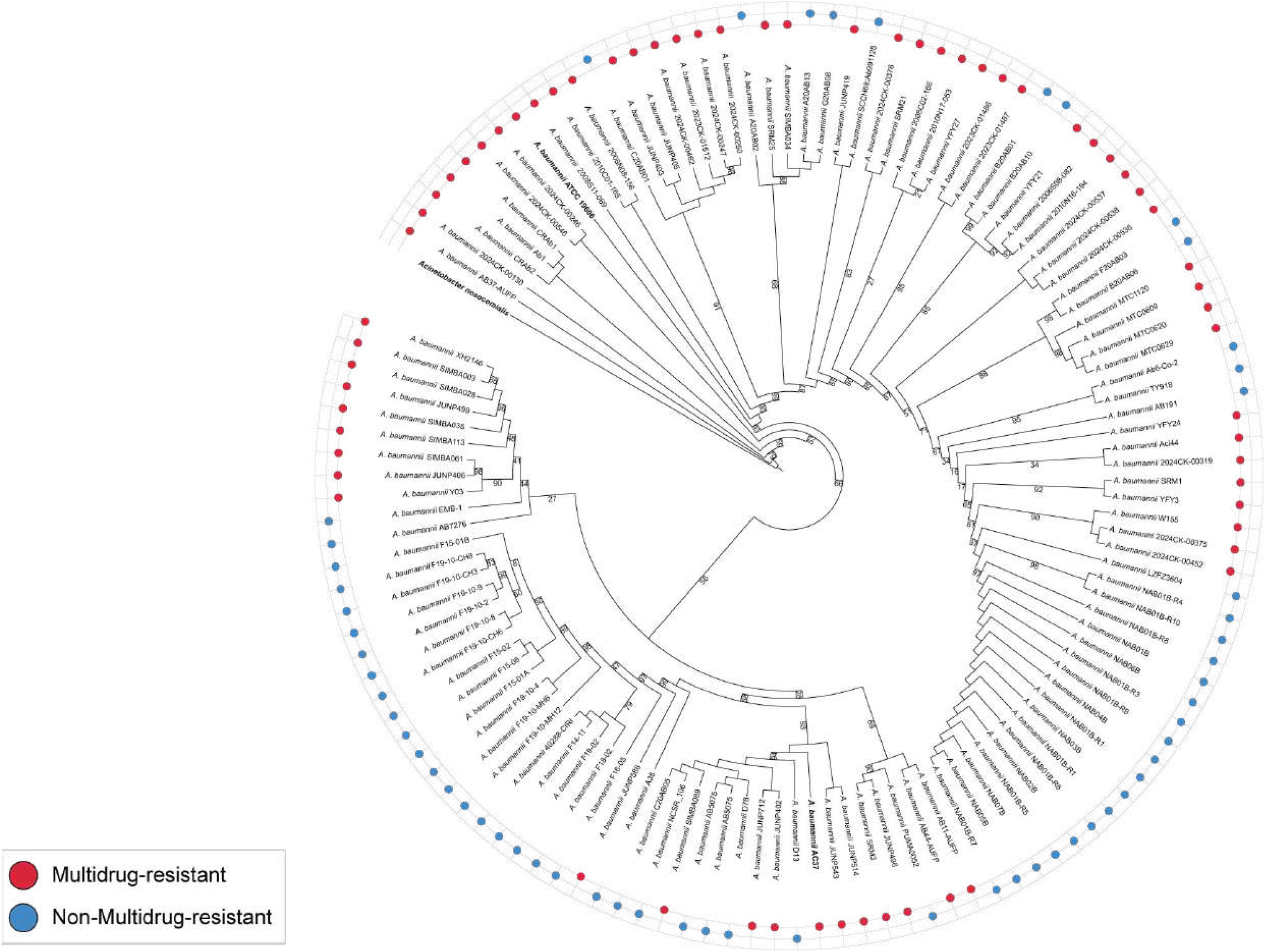
The AC37 strain is positioned within a diverse cluster, distantly related to the reference genome ATCC 19606. Phylogenetic relationships of 123 *Acinetobacter baumannii* strains. The cladogram was generated from a concatenated alignment of 366 single-copy orthologous proteins identified by BUSCO in the 124 genomes included in this study. The maximum likelihood phylogeny was inferred using IQ-TREE2 with automatic selection of the best-fit substitution model with 1000 pseudoreplicates. Bootstrap values (< 100) are displayed. Blue circles indicate non-resistant strains, and red circles denote multidrug-resistant strains.

### AC37 resistome comprises 31 genes, seven of which are carried by mobile genetic elements and 3 are unique to the AC37 genome

We systematically identified antibiotic resistance genes using the Resistance Gene Identifier (RGI) in combination with the Comprehensive Antibiotic Resistance Database (CARD). This analysis revealed 31 resistance genes, comprising 11 genes with complete identity to CARD entries and 20 genes exhibiting allelic variations **(Fig. 2-A, Table S3)**. These genes were categorized according to their resistance mechanisms, with antibiotic efflux being the most prevalent mechanism, followed by inactivation, target alteration, target replacement, and reduced permeability. Considering the tendency of resistance genes to be located within mobile genetic elements (MGEs) (19), we also characterized the mobilome of this strain **(Fig. 2-B)**. Notably, 13 genes could be within MGE regions, including 7 antibiotic resistance genes and 2 genes conferring resistance to heavy metals.

**Figure 2.**
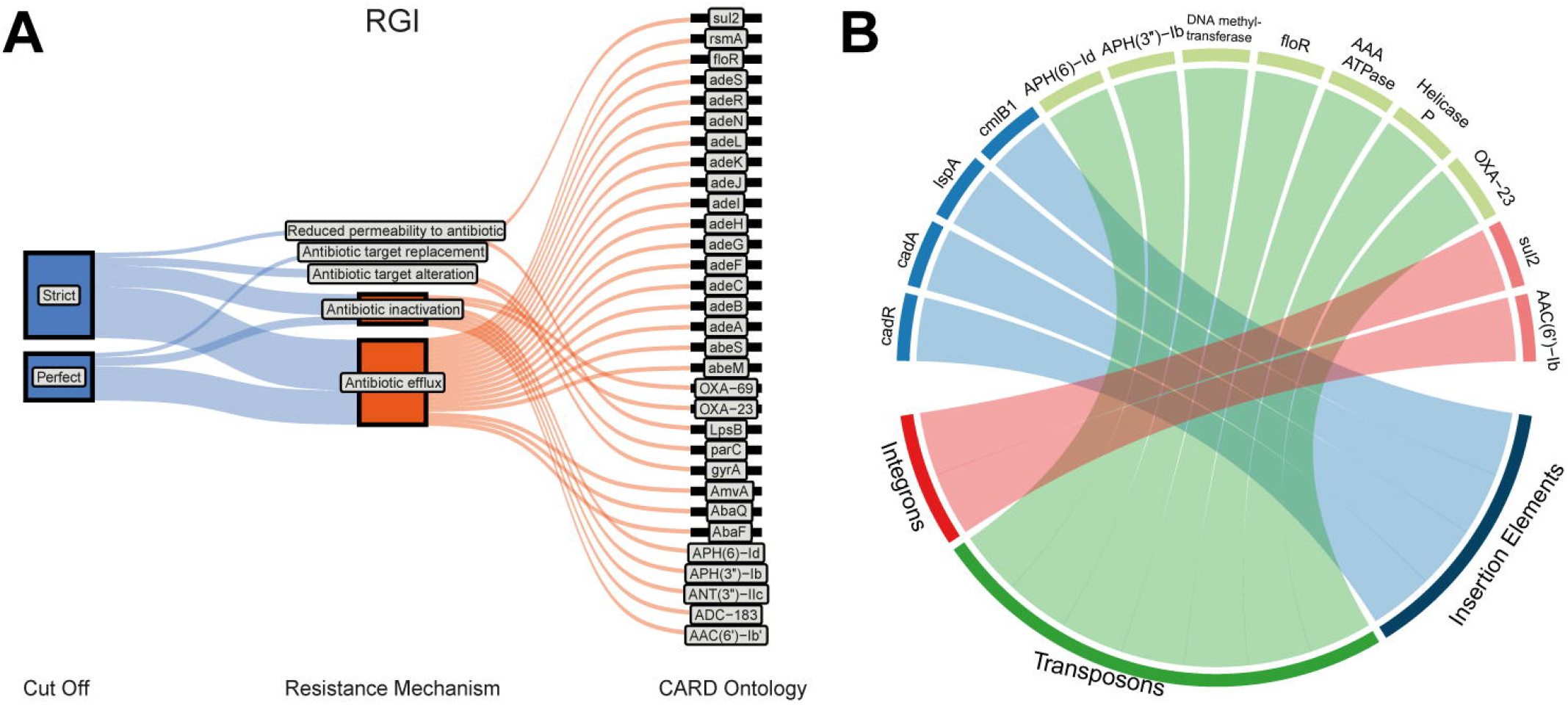
AC37 harbors 31 resistance genes, with seven located within MGEs. (A)Identification of 31 resistance genes using RGI with the CARD database in strict mode. Perfect comprises genes with complete identity to CARS entries; Strict includes genes exhibiting allelic variations. The Resistance Mechanism parameter displays the association of genes (CARD Ontology) with their mechanisms of action against antibiotics. B) Chord diagram of genes associated with transposable element regions are shown in green, genes located in insertion sequence regions in blue, and genes related to integron regions in red according to TNCentral database.

As part of the genomic analysis, we conducted a broad identification of resistance genes across all strains used to build the phylogenetic tree **(Fig. 3)**, aiming to compare resistome profiles and elucidate the distribution patterns of these genes. Our analysis revealed that a substantial proportion of resistance genes shared among all the strains were associated with antibiotic efflux mechanisms.

**Figure 3.**
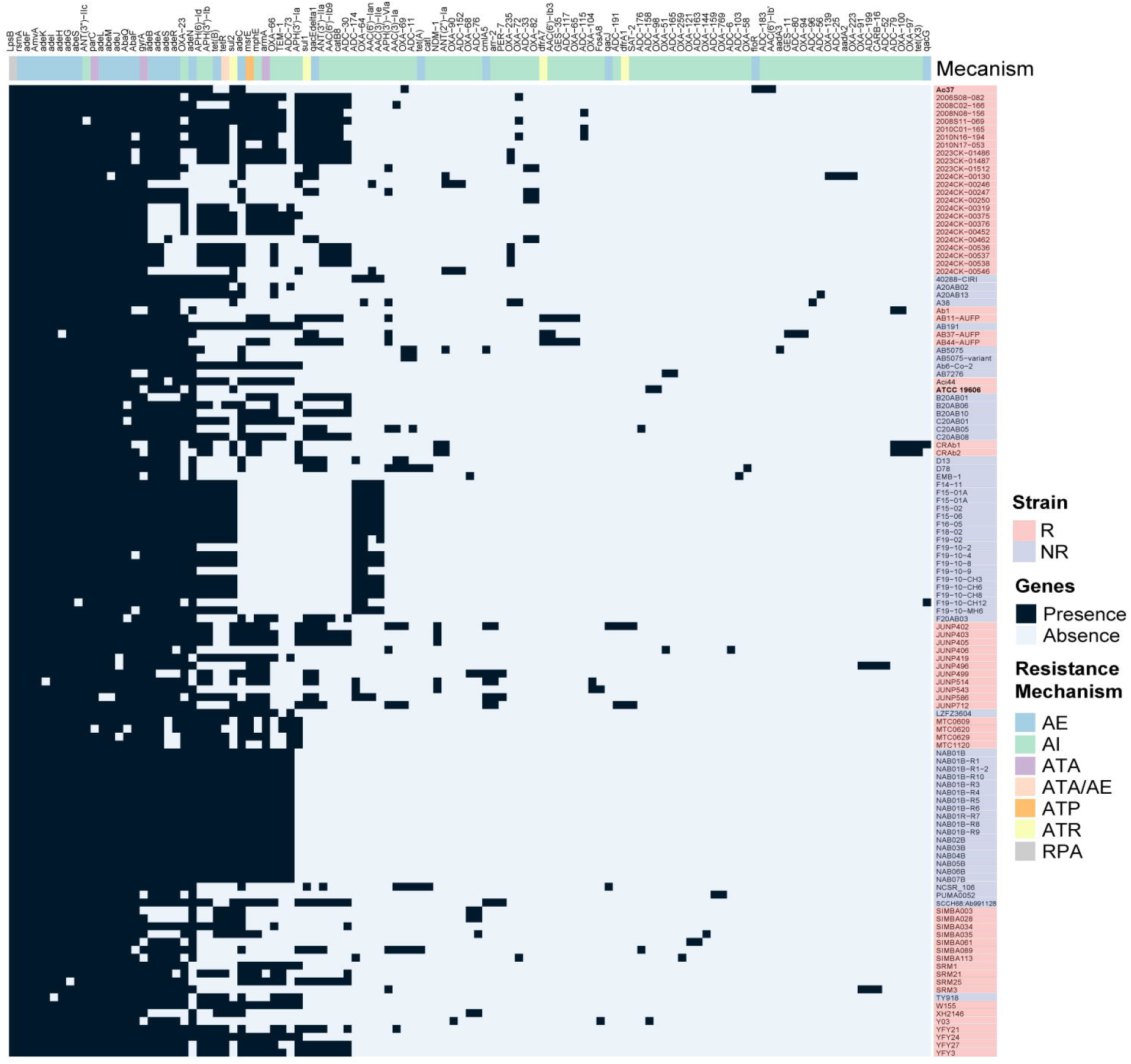
Comparative resistome analysis highlighting 3 unique genes in AC37. Presence/absence heatmap illustrating the distribution of resistance genes across the AC37 isolate and 122 other *Acinetobacter baumannii* genomes. Black squares indicate the presence of a specific resistance gene, while gray squares indicate its absence. Columns represent individual resistance genes, grouped by their resistance mechanism. Rows represent *A. baumannii* strains, with multidrug-resistant strains colored in red and non-resistant strains in blue. R: Resistant; NR: Non-resistant; AE: Antibiotic Efflux; AI: Antibiotic Inactivation; ATA: Antibiotic Target Alteration; ATA/AE: Antibiotic Target Alteration/Antibiotic Efflux; ATP: Antibiotic Target Protection; ATR: Antibiotic Target Replacement; RPA: Reduced Permeability to Antibiotic.

AC37 harbors three unique resistance genes, *AAC(6’)-Ib’, ADC-183*, and *floR*, which were not detected in the other genomes. According to CARD, these genes are relatively rare, with prevalence rates of 0.71%, 0.35%, and 2.83%, respectively, among *A. baumannii* strains. In addition, *AAC(6’)-Ib’* and *floR* genes could be located within MGEs **(Fig. 2-B)**. We next investigated the potential origin of these genes using the TNCentral database, which showed that the *AAC(6’)-Ib’* gene was initially identified in *P. aeruginosa*, a gram-negative bacterium commonly found in hospital settings (20, 21). Our data emphasizes the important role of horizontal gene transfer in the acquisition and dissemination of antibiotic resistance genes, particularly in clinical settings.

### LyeTx I mnΔK triggers broad transcriptomic changes in *A. baumannii* AC37

To elucidate the modulatory effects of the LyeTx I mnΔK peptide on gene expression in *A. baumannii*, the AC37 strain was cultured under subinhibitory concentrations of peptide (3 µM, MIC = 4µM), both alone and in combination with meropenem (1µM of LyeTx I mnΔK and 6.5 µg/mL meropenem, MIC of meropenem ≥ 16 µg/mL) (22, 23) (**Fig. S2-S3**).

Following the construction and sequencing of libraries, we obtained between 5 and 9 million paired-end reads per library. Principal component analysis (PCA) revealed distinct clustering of samples according to treatment conditions, with clear visual separation observed between different treatments (**Fig S4**).

Transcriptome analysis showed that LyeTx I mnΔK significantly modulated the expression of approximately 600 genes **(Fig. 4-A)**. Combined treatment with the peptide and meropenem also altered bacterial gene regulation, resulting in nearly 300 differentially expressed genes (DEGs) **(Fig. 4-B)**. Next, we investigated the specific effects of these treatments on the expression of previously identified resistance genes. Treatment with LyeTx I mnΔK led to the upregulation of five genes - *adeA, adeB, adeI, adeJ*, and *adeK* - with log_2_fold changes of 1,673; 1,226; 1,667; 1,463 and 1,238, respectively **(Table S4)**.

**Figure 4.**
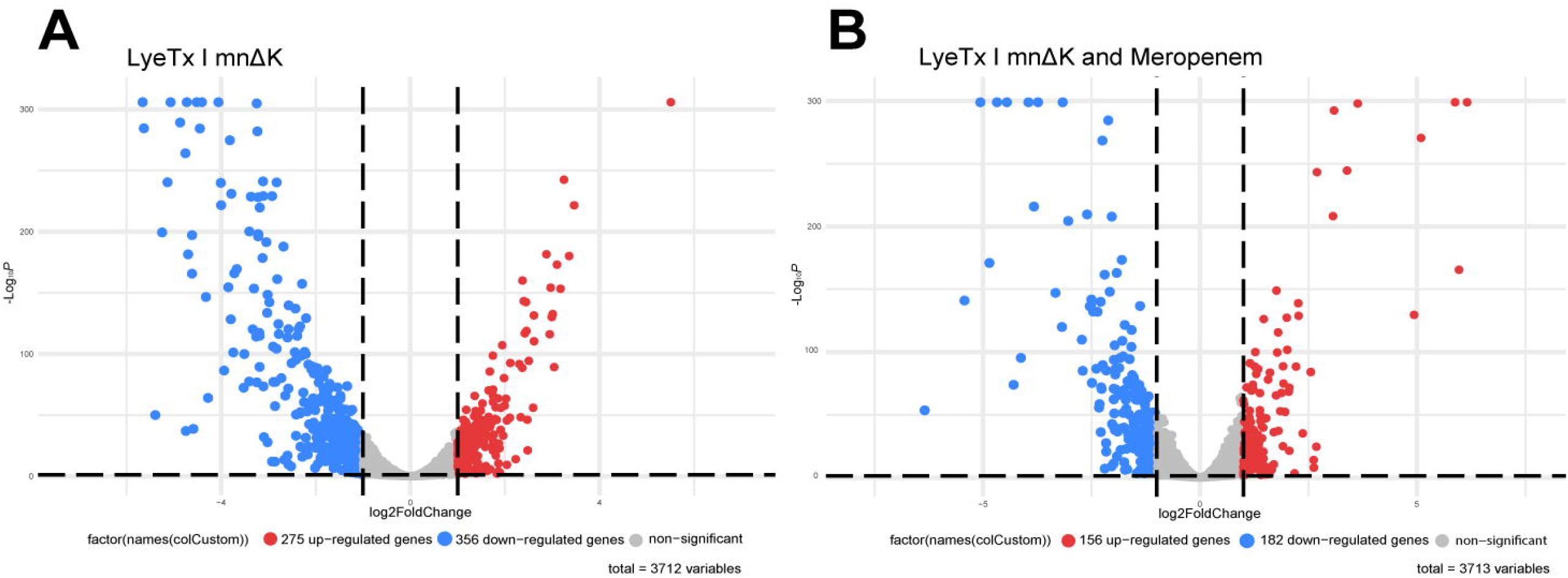
Transcriptomic response of AC37 to LyeTx I peptide and its synergistic combination with meropenem. Volcano plots illustrating differentially expressed genes (DEGs) in AC37 treated with LyeTx I alone (A) and the synergistic combination of LyeTx I and meropenem (B). The x-axis represents log2FoldChange, and the y-axis represents −log10(adjusted p-value). Blue circles indicate significantly underexpressed genes, red circles indicate significantly overexpressed genes, and gray circles represent non-significantly changed genes (absolute Log2 Fold Change < 1 and adjusted p-value ≥ 0.05).

### LyeTx I mnΔK modulates cellular metabolism by enhancing protein biosynthesis and secretion while suppressing lipid catabolism

Specific metabolic pathways modulated by the LyeTx I mnΔK peptide and its combination with meropenem were characterized through functional enrichment analyses. The results showed that treatment with the peptide as a single agent led to both activation and inhibition of several metabolic pathways. Pathways related to ribosome biogenesis, protein export, and the bacterial secretion system were significantly upregulated **(Fig. 5-A)**. In contrast, pathways involved in fatty acid and amino acid degradation were notably suppressed. Interestingly, the combined treatment with LyeTx I mnΔK and meropenem exclusively suppressed metabolic pathways, with a pronounced effect on oxidative phosphorylation **(Fig. 5-B)**. Genes within these pathways were also differentially regulated in this treatment **(Table S5)**.

**Figure 5.**
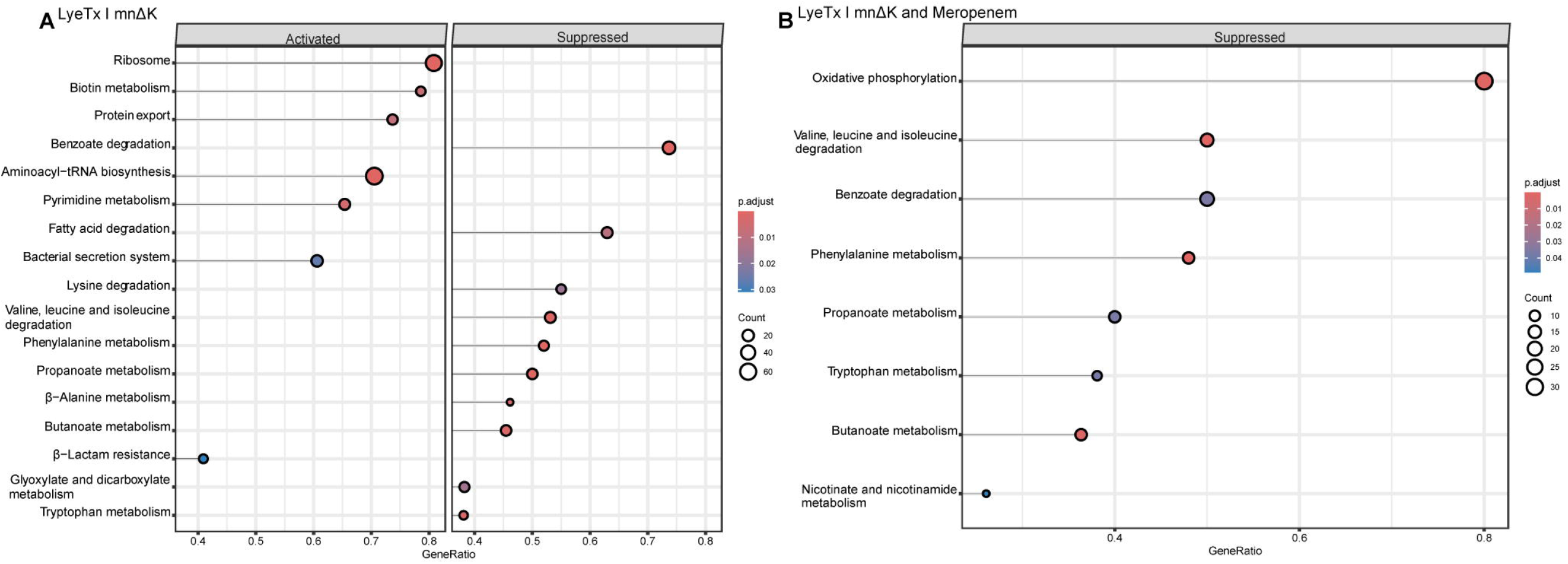
Functional enrichment of biological pathways in *Acinetobacter baumannii* AC37 treated with LyeTx I mnΔK and its synergistic combination with meropenem. Lollipop plots displaying KEGG pathways identified as significantly enriched (activated or inhibited) using gseKEGG analysis. The color gradient of the circles represents the adjusted p-value, with darker colors indicating higher significance. The diameter of each circle is proportional to the number of genes associated with the respective pathway. A) Pathways significantly enriched in response to LyeTx I mnΔK treatment. B) Pathways significantly enriched in response to the synergistic treatment of LyeTx I mnΔK and meropenem.

### Analysis of publicly available RNA-Seq data reveals time-dependent transcriptomic response to meropenem characterized by downregulation of fatty acid biosynthesis

To further explore the impact of meropenem treatment on gene expression regulation, we integrated publicly available RNA-Seq datasets of *A. baumannii* treated with this antibiotic **(Table S6)**. We selected studies using resistant strains, paired-end sequencing, and at least three biological replicates per condition, to ensure comparability. Our search identified a single study (SRA project PRJNA787205) meeting these criteria, which examined the transcriptional response of *A. baumannii* strain ATCC 17978 at 30 minutes, 3 hours, and 9 hours post-meropenem treatment. Analysis of the differential gene expression data revealed a time-dependent response, with a progressive increase in differentially expressed genes over time (**Fig. S5, Table S7**) upon meropenem. Functional enrichment analysis of these regulated genes showed a pronounced downregulation of several metabolic pathways, including biotin metabolism, fatty acid and specific amino acids biosynthesis (**Fig. S6**).

Lastly, we analyzed the expression of genes associated with glycolysis/gluconeogenesis pathways, fatty acid metabolism and oxidative phosphorylation - as annotated in the KEGG database. An antagonistic pattern was observed when comparing gene expression in response to the peptide and meropenem, most notably within the fatty acid metabolism genes (**Fig. 6**). We also notice the oxidative phosphorylation genes exhibited a widespread downregulation of the majority of detected genes, indicating that the synergistic effect of these two antibiotics has a significant impact in this pathway as part of the bacterial adaptive response.

**Figure 6.**
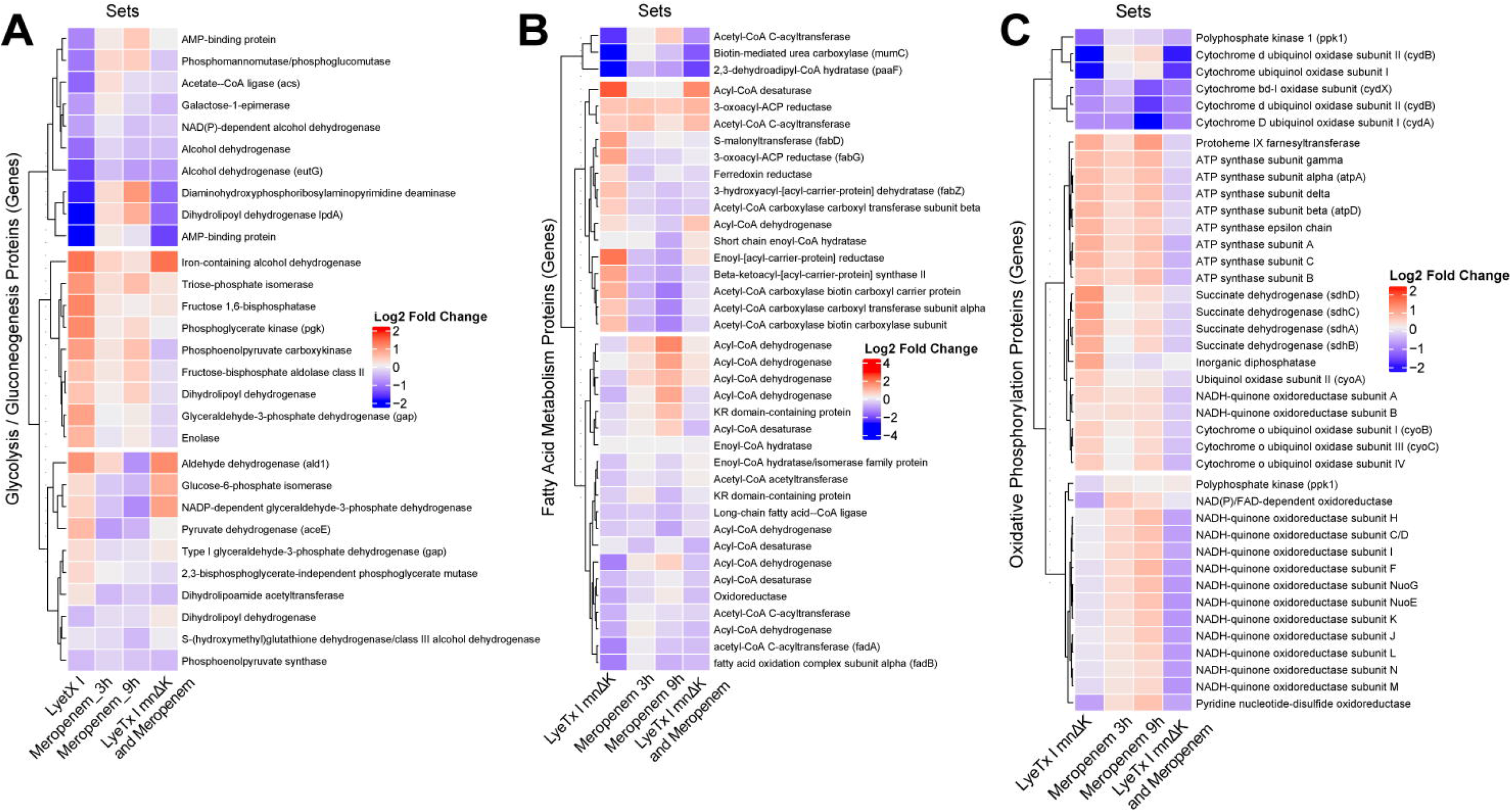
Differential expression patterns of key metabolic pathways across various antimicrobial treatments. Heatmaps displaying the log2 Fold Change in gene expression for (A) glycolysis and gluconeogenesis pathway genes, (B) fatty acid metabolism genes, and (C) oxidative phosphorylation pathway genes. Columns represent the following treatment conditions: LyeTx I mnΔK, Meropenem (3 hours), Meropenem (9 hours), and the synergistic combination of LyeTx I mnΔK and Meropenem. The color scale, shown at the bottom, indicates the magnitude and direction of gene expression change relative to the untreated control.

## DISCUSSION

This study was initiated with the genome sequencing, assembly, and annotation of the novel *A. baumannii* isolate - AC37 - followed by comparative genomic analyses with other *A. baumannii* strains to position AC37 within the specieś genomic landscape and explore its antimicrobial resistance. We also explored the transcriptional response of AC37 to the antimicrobial peptide LyeTx I mnΔK, both alone and in combination with meropenem, in an attempt to elucidate the molecular mechanisms underlying its antimicrobial effects and potential synergistic interactions.

In the AC37 genome, we identified a single plasmid (NZ_CP131945.1) that, despite lacking antibiotic resistance genes commonly associated with plasmids, harbored the BrnT/BrnA toxin-antitoxin (TA) system. This operon encodes the ribonuclease BrnTp and its cognate antitoxin BrnAp, which assemble into a 2:2 complex that autoregulates its transcription (17, 18). TA systems are frequently linked to plasmid stabilization in association with resistance determinants (24). However, the presence of BrnT/BrnA in a plasmid devoid of such genes highlights a potentially distinct role, more closely tied to stress physiology and plasmid maintenance. These systems are activated in response to multiple environmental stressors-including nutrient starvation, oxidative stress, pH imbalance, temperature changes and antimicrobial exposure- and can trigger persister cell formation (25, 26). Such mechanism is specially relevant for *A. baumannii*, a species characterized by robust biofilm formation and environmental resilience (27, 28). Thus, the BrnT/BrnA system on the NZ_CP131945.1 plasmid may confer an adaptive advantage by enhancing plasmid stability and promoting bacterial persistence under hostile conditions, possibly contributing to the pathogenicity of AC37 strain.

The analysis of 123 *A. baumannii* genomes revealed an average of 31.8 antibiotic resistance genes, a finding consistent with the observations of Hernández-González et al., (29), who reported an average of 29.4 resistance genes in their extensive study of 1472 *A. baumannii* genomes. The majority of shared resistance genes among the analyzed genomes were associated with antibiotic efflux mechanisms, predominantly involving the Resistance-Nodulation-Division (RND) efflux pump family.

Among the resistance genes identified in strain AC37, three were found to be unique: *AAC(6’)-Ib’, floR*, and *ADC-183*, whose low frequency was confirmed in a previous study (29). *AAC(6’)-Ib’* gene, which encodes an aminoglycoside acetyltransferase commonly found in various bacterial families (30–33), was included in a class I integron in AC37, with a potential origin in *P. aeruginosa*. Its ability to disseminate via MGEs (34) reinforces its role in the horizontal transfer of aminoglycoside resistance, potentially between species such as *P. aeruginosa* and *A. baumannii* in hospital environments. The *ADC-183* gene, exclusive of the *Acinetobacter* genus, encodes an AmpC β-lactamase that confers resistance to β-lactam antibiotics through hydrolysis (35–37). Finally, the *floR* gene originally identified in 1999 (38) encodes a phenicol exporter and was located within a mobile element in AC37, raising concerns due to its potential dissemination in hospital settings (39, 40).

Prior genomic characterization of the AC37 isolate was essential for the transcriptome analysis, given the observed divergences from the reference genome ATCC 19606, which would lead to suboptimal read mapping and loss of important information regarding treatment responses. Following accurate genome annotation of the AC37 isolate, we analyzed its transcriptional response to treatment with the antimicrobial peptide LyeTx I mnΔK and its combination with meropenem. LyeTx I mnΔK has shown promising bactericidal activity coupled with minimal toxicity (15). Previous studies have reported a synergistic effect with meropenem, enhancing antimicrobial efficacy against multidrug-resistant *A. baumannii* isolates (16). Given the growth of antibiotic resistance, exploring such novel therapeutic strategies is crucial.

We observed upregulation of *adeA, adeB, adeI, adeJ*, and *adeK* genes in LyeTx I mnΔK treatment, which encode RND efflux pumps, a key resistance mechanism that expels antimicrobials using the proton motive force (41–43). Overexpression of the adeABC operon, especially the *adeB* gene (a multidrug transporter), is characteristic of a MDR *A. baumannii*, conferring resistance to β-lactams, aminoglycosides, and chloramphenicol through active extrusion (44–47). The unchanged expression of *adeC* suggests possible use of alternative outer membrane proteins, reflecting the functional plasticity of the adeAB system (48). While the adeIJK operon is present in susceptible isolates, its overexpression contributes to resistance (49), exhibiting different substrate specificity from adeABC, and influencing the different resistance profiles among *A. baumannii* strains.

Treatment with the LyeTx I mnΔK peptide led to the upregulation of processes related to protein synthesis, export, and secretion, as well as pyrimidine metabolism and β-lactam resistance. Concerning protein export process, the Sec-dependent pathway showed a tendency of upregulation of several genes such as secY, secF, secE and secG (log_2_ fold changes of: 1,053; 1,016; 0,791 and 0,719, respectively). This pathway plays a pivotal role in bacterial physiology, facilitating the translocation of approximately one-third of the bacterial proteome to extracytoplasmic compartments, including the plasma membrane and periplasm (50, 51). The observed upregulation of genes related to the Sec-dependent pathway in response to LyeTx I mnΔK treatment suggests a potential adaptive response aimed at maintaining or remodeling membrane integrity under peptide-induced stress. This may reflect a cellular strategy to counteract the disruptive effects of the peptide on membrane structure and function.

Publicly available transcriptomic data from *A. baumannii* treated with meropenem revealed decreased expression of the fatty acid biosynthesis pathway.

We hypothesize that this downregulation reallocates the cellular pool of acetyl-CoA towards the tricarboxylic acid (TCA) cycle, thereby enhancing the generation of reducing equivalents (NADH and FADH_2_) required for ATP production via oxidative phosphorylation. This hypothesis is supported by the observed overexpression of several genes involved in oxidative phosphorylation in the same dataset, such as ATP synthase and NADH-quinone oxidoreductase subunits. Our results align with two previous studies. The first demonstrated increased expression of genes related to the TCA cycle and ATP synthase in *A. baumannii* treated with amikacin, imipenem, and meropenem (52). The second, focusing on a colistin-resistant strain, identified a marked suppression of the fatty acid biosynthesis pathway (53), collectively highlighting that this metabolic shift may represent a strategy to meet the increased energy demands associated with the stress response to meropenem-induced cell wall damage.

Synergistic treatment with LyeTx I mnΔK and meropenem consistently led to the pattern of metabolic pathway suppression. The oxidative phosphorylation pathway was repressed, with genes encoding subunits of the NADH-quinone oxidoreductase - NADH dehydrogenase complex I - exhibiting a tendency of downregulation (such as NuoD, NuoE and NuoG). This suggests a potential disruption of the proton motive force (PMF), as this complex is the first in the electron transporter chain and promotes proton translocation across the bacterial inner membrane. Similarly, genes encoding the cytochrome bd oxidase complex (*CydA, CydB*, and *CydX*), an important compound in bacterial respiratory electron transfer that reduces molecular oxygen to water (54) were also considerably downregulated. The combined suppression of these complexes directly impairs the establishment of the PMF, which is essential for ATP production via oxidative phosphorylation. These findings align with previous studies reporting that inhibition of the electron transport chain and subsequent disruption of the electrochemical gradient across the membrane lead to reduced ATP synthesis, decreased virulence, and impaired cell viability (55). The observed suppression of this pathway under the combined treatment highlights the potential of LyeTx I mnΔK and meropenem to compromise bacterial energy metabolism and viability.

The transcriptomic data highlighted complementary biological responses to individual treatments and these results emphasize the metabolic impact and potential therapeutic advantage of the combined antimicrobial strategy. The differentially expressed genes identified in the study offered insights into the putative molecular mechanism underlying LyeTx I mnΔK action and uncovered novel targets for further drug development investigation.

## EXPERIMENTAL PROCEDURES

### gDNA extraction, genome sequencing and phylogeny

*A. baumannii* strain AC37, isolated from a central venous catheter obtained from a patient at Hospital Santa Casa de Belo Horizonte, was subjected to whole-genome sequencing. Bacteria were grown in Mueller-Hinton broth at 37 ºC until the culture reached an optical density at 600 nm (OD_600_) of 0.8. Genomic DNA was extracted using the QIAamp DNA Mini Kit (QIAGEN), following the manufacturer’s instructions. DNA concentration was determined by fluorometry using the Qubit dsDNA BR Assay Kit (Thermo Fisher Scientific), and its integrity was assessed by electrophoresis on an Agilent 4150 TapeStation System (Agilent Technologies). Libraries were prepared using the Illumina DNA Prep Kit (Illumina). Sequencing was performed on a NextSeq 2000 System (Illumina), generating 5,117,651 paired-end reads with an average length of 301 base pairs.

Genome assembly was performed with Unicycler v. 0.5.1 (56), with the following parameters: --nominiasm, --spades_options “--isolate”, --no_simple_bridges e --no_long_read_aligment. Genome annotation was carried out using the NCBI Prokaryotic Genome Annotation Pipeline (PGAP, v. build7555) (57). Genome visualization was achieved through a local installation of Bandage (Bioinformatics Application for Navigating De novo Assembly Graphs Easily, v.0.9.0) (58). Plasmid identification was conducted using the PLSDB (Plasmid Database, available at: https://ccb-microbe.cs.uni-saarland.de/plsdb2025/) web interface, with the following parameters: max_pvalue = 0.01, min_ident = 99, winner_takes_all = false.

A set of 123 *A. baumannii* genomes, including the reference genome of *A. baumannii* ATCC 19606 (assembly ASM903584v1) and *Acinetobacter nosocomialis* (assembly XH1679) as an outgroup, were retrieved from the NCBI (National Center for Biotechnology Information) database. The selection criteria for these genomes included a deposition date between January 1st, 2024, and September 14th, 2024, complete genome status, exclusion of metagenomic assemblies, and exclusion of atypical genomes. The assembly codes of the 124 genomes are listed in **Table S2**.

Genome completeness was assessed using BUSCO v.5.5.0 (Benchmarking Universal Single-Copy Orthologs) (59), utilizing the gammaproteobacteria_odb10 database. 366 complete single-copy orthologous proteins were selected using the BUSCO_phylogenomics.py script (available at: https://github.com/jamiemcg/BUSCO_phylogenomics), which performs multiple sequence alignment with MUSCLE and removes unaligned ends with trimAL. IQ-TREE2 v. 2.0.7 (60) was employed to construct the maximum likelihood phylogenetic tree. The robustness of the phylogenetic tree was evaluated through bootstrap with 1000 pseudoreplicates. The output in NEXUS format was visualized and edited using the ITOL web interface tool (61). Finally, 123 *A. baumannii* genomes and the reference genome of *A. nosocomialis*, as an outgroup, were used for the phylogenetic analysis.

### Identification of resistance genes and mobile genetic elements

Resistance genes were identified using RGI v. 6.0.3 (The Resistance Gene Identifier) (62), utilizing the CARD (Comprehensive Antibiotic Resistance Database) as a reference for bacterial resistome prediction. Mobile genetic elements were identified through BLASTn against TNCentral (63), a curated database of prokaryotic MGEs. The output generated by BLASTn was manually analyzed on the TNCentral and ISFinder (64) web pages to remove redundancies. Manual curation involved verifying the output hits and comparing the genomic coordinates of the identified genes with the expected sizes in the databases to remove redundancies.

### RNA extraction and sequencing

For transcriptomic analysis, the *A. baumannii* strain AC37 was grown in Mueller-Hinton broth at 37 ºC under three experimental conditions: untreated (control), treated with 3 µM of the LyeTx I mnΔK peptide, and treated with a combination of 1 µM of the LyeTx I mnΔK peptide and 6.5 µg/mL of meropenem. After control cultures reached an OD_600_ of 0.8, all sample groups were collected. Total RNA was then stabilized and purified using the RNeasy Protect Bacteria Mini Kit (QIAGEN), following the enzymatic and mechanical lysis protocol. Total RNA concentration was determined by fluorometry using the Qubit RNA BR Assay Kit (Thermo Fisher Scientific), and its integrity was assessed by electrophoresis on an Agilent 4150 TapeStation System (Agilent Technologies). Libraries were prepared using the Illumina Stranded Total RNA Prep, Ligation with Ribo-Zero Plus (Illumina). Sequencing was performed on a NextSeq 550 System (Illumina), generating between 5 and 9 million paired-end reads with an average length of 75 base pairs. Growth curves for each treatment are presented in Fig S3.

Public RNA-Seq data from the treatment of *A. baumannii* with meropenem were retrieved from the Sequence Read Archive (SRA) using the MeSH terms “Acinetobacter baumannii AND meropenem”. This search yielded 7 studies. The characteristics of the available libraries varied considerably, including the specific *A. baumannii* strain, read fragment length, meropenem concentration, sequencing depth, read type (single-end or paired-end), and the number of biological replicates, as summarized in **Table S3**. To ensure greater data homogeneity and facilitate direct comparisons with our in-house RNA-Seq data, we applied selection criteria based on paired-end read type and a minimum of three biological replicates per experimental condition. Following these criteria, a single SRA study (PRJNA787205) was deemed suitable for our comparative analysis.

Quality control of DNA-Seq and RNA-Seq libraries was performed using FastQC v. 0.11.9 (65) and MultiQC v. 1.13.dev0 (66).

### Transcriptome analyses

The read alignment from the raw RNA-Seq libraries was performed using Bowtie2 (67), with default parameters. Conversion and sorting of BAM to SAM files was performed using Samtools v. 1.16.1 (68). The number of reads mapped to each gene was counted using FeatureCounts v. 2.0.3 (69). To visualize the variance of samples treated with antimicrobials compared to the control, principal component analysis (PCA) was performed using the prcomp function on R 4.4.3 using the RStudio 2024,12.1.563 platform.

Differential gene expression analysis was performed in R using the DESeq2 v.1.40.2 package (70). The cutoff value used to consider a gene as differentially expressed (DEG) was an absolute log2FoldChange ≥ 1, i.e., at least 2 times higher expression compared to the untreated condition (control); and an adjusted p-value (padj) < 0.05. Functional enrichment analysis was performed with the gseKEGG function from the clusterProfiler v.4.8.3 package. This function ranks all genes using the pathways annotated in the KEGG (Kyoto Encyclopedia of Genes and Genomes) database (71).

## DATA AVAILABILITY

The raw transcriptomic data, genomic and the assembled genome have been deposited in NCBI’s Sequence Read Archive (SRA) under the BioProject accession PRJNA1260460.

## ACKNOWLEDGMENTS

We thank the team from the Laboratório de Genética e Bioquímica (LGB) at UFMG for constructive data discussion. Computational analyses were performed on the Gloriosos clusters (PPG Bioinfo UFMG) and Chapolin clusters (UFSB).

## AUTHOR CONTRIBUTIONS

**FGCO, KOB, MEL**, and **GRF** conceptualized the project. **KOB, WGL, AMV** and **GRF** advised on the bioinformatics analysis. **FGCO** and **DOV** analyzed data. **JRM, JCD, DLS**, and **FAS** performed all wet lab experiments. **IM** submitted DNA-Seq and RNA-Seq raw data to SRA (NCBI). **TMB and IM** provided technical support and infrastructure maintenance for bioinformatics analyses. **FGCO** wrote and edited the manuscript. **CAR, VLS, MEL**, and **GRF** secured funding for the research. **FGCO, KOB, DOV, JRM, JCD, DLS, IM, RGM, RSA, FAS, VLS, AMV, TMB, CRM, WGL CAR, MEL**, and **GRF** reviewed, edited, and approved the manuscript.

## REFERENCES

1. Ramirez, M. S., Bonomo, R. A., and Tolmasky, M. E. (2020) Carbapenemases: Transforming Acinetobacter baumannii into a Yet More Dangerous Menace. Biomolecules. 10, 720

2. Ibrahim, S., Al-Saryi, N., Al-Kadmy, I. M. S., and Aziz, S. N. (2021) Multidrug-resistant Acinetobacter baumannii as an emerging concern in hospitals. Mol. Biol. Rep. 48, 6987–6998

3. Müller, C., Reuter, S., Wille, J., Xanthopoulou, K., Stefanik, D., Grundmann, H., Higgins, P. G., and Seifert, H. (2023) A global view on carbapenem-resistant Acinetobacter baumannii. mBio. 14, e02260–23

4. Antunes, L. C. S., Visca, P., and Towner, K. J. (2014) Acinetobacter baumannii : evolution of a global pathogen. Pathog. Dis. 71, 292–301

5. Lee, C.-R., Lee, J. H., Park, M., Park, K. S., Bae, I. K., Kim, Y. B., Cha, C.-J., Jeong, B. C., and Lee, S. H. (2017) Biology of Acinetobacter baumannii: Pathogenesis, Antibiotic Resistance Mechanisms, and Prospective Treatment Options. Front. Cell. Infect. Microbiol. 10.3389/fcimb.2017.00055

6. Gordon, N. C., and Wareham, D. W. (2010) Multidrug-resistant Acinetobacter baumannii: mechanisms of virulence and resistance. Int. J. Antimicrob. Agents. 35, 219–226

7. Cerceo, E., Deitelzweig, S. B., Sherman, B. M., and Amin, A. N. (2016) Multidrug-Resistant Gram-Negative Bacterial Infections in the Hospital Setting: Overview, Implications for Clinical Practice, and Emerging Treatment Options. Microb. Drug Resist. 22, 412–431

8. Giammanco, A., Calà, C., Fasciana, T., and Dowzicky, M. J. (2017) Global Assessment of the Activity of Tigecycline against Multidrug-Resistant Gram-Negative Pathogens between 2004 and 2014 as Part of the Tigecycline Evaluation and Surveillance Trial. mSphere. 2, e00310–16

9. Doi, Y. (2019) Treatment Options for Carbapenem-resistant Gram-negative Bacterial Infections. Clin. Infect. Dis. Off. Publ. Infect. Dis. Soc. Am. 69, S565–S575

10. Nordmann, P., and Poirel, L. (2019) Epidemiology and Diagnostics of Carbapenem Resistance in Gram-negative Bacteria. Clin. Infect. Dis. 69, S521–S528

11. Kyriakidis, I., Vasileiou, E., Pana, Z. D., and Tragiannidis, A. (2021) Acinetobacter baumannii Antibiotic Resistance Mechanisms. Pathog. Basel Switz. 10, 373

12. Matsuzaki, K. (2001) Why and how are peptide-lipid interactions utilized for self defence? Biochem. Soc. Trans. 29, 598–601

13. Mallapragada, S., Wadhwa, A., and Agrawal, P. (2017) Antimicrobial peptides: The miraculous biological molecules. J. Indian Soc. Periodontol. 21, 434

14. Li, X., Zuo, S., Wang, B., Zhang, K., and Wang, Y. (2022) Antimicrobial Mechanisms and Clinical Application Prospects of Antimicrobial Peptides. Molecules. 27, 2675

15. Fuscaldi, L. L., De Avelar Júnior, J. T., Dos Santos, D. M., Boff, D., De Oliveira, V. L. S., Gomes, K. A. G. G., Cruz, R. D. C., De Oliveira, P. L., Magalhães, P. P., Cisalpino, P. S., Farias, L. D. M., De Souza-Fagundes, E. M., Delp, J., Leist, M., Resende, J. M., Amaral, F. A., Pimenta, A. M. D. C., Fernandes, S. O. A., Cardoso, V. N., and De Lima, M. E. (2021) Shortened derivatives from native antimicrobial peptide LyeTx I: In vitro and in vivo biological activity assessment. Exp. Biol. Med. 246, 414–425

16. Lima, W. G., Brito, J. C. M., De Lima, M. E., Pizarro, A. C. S. T., Vianna, M. A. M. D. M., De Paiva, M. C., De Assis, D. C. S., Cardoso, V. N., and Fernandes, S. O. A. (2021) A short synthetic peptide, based on LyeTx I from Lycosa erythrognatha venom, shows potential to treat pneumonia caused by carbapenem-resistant Acinetobacter baumannii without detectable resistance. J. Antibiot. (Tokyo). 74, 425–434

17. Heaton, B. E., Herrou, J., Blackwell, A. E., Wysocki, V. H., and Crosson, S. (2012) Molecular Structure and Function of the Novel BrnT/BrnA Toxin-Antitoxin System of Brucella abortus. J. Biol. Chem. 287, 12098–12110

18. Jurėnas, D., Fraikin, N., Goormaghtigh, F., and Van Melderen, L. (2022) Biology and evolution of bacterial toxin–antitoxin systems. Nat. Rev. Microbiol. 20, 335–350

19. Tokuda, M., and Shintani, M. (2024) Microbial evolution through horizontal gene transfer by mobile genetic elements. Microb. Biotechnol. 17, e14408

20. Jurado-Martín, I., Sainz-Mejías, M., and McClean, S. (2021) Pseudomonas aeruginosa: An Audacious Pathogen with an Adaptable Arsenal of Virulence Factors. Int. J. Mol. Sci. 22, 3128

21. Diggle, S. P., and Whiteley, M. (2020) Microbe Profile: Pseudomonas aeruginosa: opportunistic pathogen and lab rat. Microbiology. 166, 30–33

22. Zhanel, G. G., Wiebe, R., Dilay, L., Thomson, K., Rubinstein, E., Hoban, D. J., Noreddin, M., and Karlowsky, J. A. (2007) Comparative Review of the Carbapenems: Drugs. 67, 1027–1052

23. Steffens, N. A., Zimmermann, E. S., Nichelle, S. M., and Brucker, N. (2021) Meropenem use and therapeutic drug monitoring in clinical practice: a literature review. J. Clin. Pharm. Ther. 46, 610–621

24. Lobato-Márquez, D., Díaz-Orejas, R., and García-del Portillo, F. (2016) Toxin-antitoxins and bacterial virulence. FEMS Microbiol. Rev. 40, 592–609

25. Lewis, K. (2010) Persister Cells. Annu. Rev. Microbiol. 64, 357–372

26. Tu, Q., Pu, M., Li, Y., Wang, Y., Li, M., Song, L., Li, M., An, X., Fan, H., and Tong, Y. (2023) Acinetobacter Baumannii Phages: Past, Present and Future. Viruses. 15, 673

27. Mea, H. J., Yong, P. V. C., and Wong, E. H. (2021) An overview of Acinetobacter baumannii pathogenesis: Motility, adherence and biofilm formation. Microbiol. Res. 247, 126722

28. Yang, C.-H., Su, P.-W., Moi, S.-H., and Chuang, L.-Y. (2019) Biofilm Formation in Acinetobacter Baumannii: Genotype-Phenotype Correlation. Molecules. 24, 1849

29. Hernández-González, I. L., Mateo-Estrada, V., and Castillo-Ramirez, S. (2022) The promiscuous and highly mobile resistome of Acinetobacter baumannii. Microb. Genomics. 8, 000762

30. Vetting, M. W., Park, C. H., Hegde, S. S., Jacoby, G. A., Hooper, D. C., and Blanchard, J. S. (2008) Mechanistic and Structural Analysis of Aminoglycoside N-Acetyltransferase AAC(6′)-Ib and Its Bifunctional, Fluoroquinolone-Active AAC(6′)-Ib-cr Variant. Biochemistry. 47, 9825–9835

31. Green, K. D., Chen, W., Houghton, J. L., Fridman, M., and Garneau-Tsodikova, S. (2010) Exploring the Substrate Promiscuity of Drug-Modifying Enzymes for the Chemoenzymatic Generation of N-Acylated Aminoglycosides. ChemBioChem. 11, 119–126

32. Ramirez, M. S., and Tolmasky, M. E. (2010) Aminoglycoside Modifying Enzymes. Drug Resist. Updat. Rev. Comment. Antimicrob. Anticancer Chemother. 13, 151–171

33. Ramirez, M. S., and Tolmasky, M. E. (2017) Amikacin: Uses, Resistance, and Prospects for Inhibition. Mol. J. Synth. Chem. Nat. Prod. Chem. 22, 2267

34. Ramirez, M. S., Nikolaidis, N., and Tolmasky, M. E. (2013) Rise and dissemination of aminoglycoside resistance: the aac(6′)-Ib paradigm. Front. Microbiol. 4, 121

35. Karah, N., Jolley, K. A., Hall, R. M., and Uhlin, B. E. (2017) Database for the ampC alleles in Acinetobacter baumannii. PLOS ONE. 12, e0176695

36. Zhao, W.-H., and Hu, Z.-Q. (2012) Acinetobacter : A potential reservoir and dispenser for β-lactamases. Crit. Rev. Microbiol. 38, 30–51

37. Bush, K. (2018) Past and Present Perspectives on β-Lactamases. Antimicrob. Agents Chemother. 62, e01076–18

38. Arcangioli, M.-A., Leroy-Sétrin, S., Martel, J.-L., and Chaslus-Dancla, E. (1999) A new chloramphenicol and florfenicol resistance gene flanked by two integron structures in Salmonella typhimurium DT104. FEMS Microbiol. Lett. 174, 327–332

39. Lu, J., Zhang, J., Xu, L., Liu, Y., Li, P., Zhu, T., Cheng, C., Lu, S., Xu, T., Yi, H., Li, K., Zhou, W., Li, P., Ni, L., and Bao, Q. (2018) Spread of the florfenicol resistance floR gene among clinical Klebsiella pneumoniae isolates in China. Antimicrob. Resist. Infect. Control. 7, 127

40. Boiten, K. E., Kuijper, E. J., Schuele, L., Van Prehn, J., Bode, L. G. M., Maat, I., Van Asten, S. A. V., Notermans, D. W., Rossen, J. W. A., and Veloo, A. C. M. (2023) Characterization of mobile genetic elements in multidrug-resistant Bacteroides fragilis isolates from different hospitals in the Netherlands. Anaerobe. 81, 102722

41. Coyne, S., Courvalin, P., and Périchon, B. (2011) Efflux-Mediated Antibiotic Resistance in Acinetobacter spp. Antimicrob. Agents Chemother. 55, 947–953

42. Athar, M., Gervasoni, S., Catte, A., Basciu, A., Malloci, G., Ruggerone, P., and Vargiu, A. V. (2023) Tripartite efflux pumps of the RND superfamily: what did we learn from computational studies? Microbiol. Read. Engl. 169, 001307

43. Xu, C. F., Bilya, S. R., and Xu, W. (2019) adeABC efflux gene in Acinetobacter baumannii. New Microbes New Infect. 30, 100549

44. Ruzin, A., Keeney, D., and Bradford, P. A. (2007) AdeABC multidrug efflux pump is associated with decreased susceptibility to tigecycline in Acinetobacter calcoaceticus–Acinetobacter baumannii complex. J. Antimicrob. Chemother. 59, 1001–1004

45. Rumbo, C., Gato, E., López, M., Ruiz De Alegría, C., Fernández-Cuenca, F., Martínez-Martínez, L., Vila, J., Pachón, J., Cisneros, J. M., Rodríguez-Baño, J., Pascual,, Bou, G., and Tomás, M. (2013) Contribution of Efflux Pumps, Porins, and β-Lactamases to Multidrug Resistance in Clinical Isolates of Acinetobacter baumannii. Antimicrob. Agents Chemother. 57, 5247–5257

46. Yuhan, Y., Ziyun, Y., Yongbo, Z., Fuqiang, L., and Qinghua, Z. (2016) Over expression of AdeABC and AcrAB-TolC efflux systems confers tigecycline resistance in clinical isolates of Acinetobacter baumannii and Klebsiella pneumoniae. Rev. Soc. Bras. Med. Trop. 49, 165–171

47. Su, C.-C., Morgan, C. E., Kambakam, S., Rajavel, M., Scott, H., Huang, W., Emerson, C. C., Taylor, D. J., Stewart, P. L., Bonomo, R. A., and Yu, E. W. (2019) Cryo-Electron Microscopy Structure of an Acinetobacter baumannii Multidrug Efflux Pump. mBio. 10, 10.1128/mbio.01295-19

48. Marchand, I., Damier-Piolle, L., Courvalin, P., and Lambert, T. (2004) Expression of the RND-Type Efflux Pump AdeABC in Acinetobacter baumannii Is Regulated by the AdeRS Two-Component System. Antimicrob. Agents Chemother. 48, 3298–3304

49. Tambat, R., Kinthada, R. K., Saral Sariyer, A., Leus, I. V., Sariyer, E., D’Cunha, N., Zhou, H., Leask, M., Walker, J. K., and Zgurskaya, H. I. (2024) AdeIJK Pump-Specific Inhibitors Effective against Multidrug Resistant Acinetobacter baumannii. ACS Infect. Dis. 10, 2239–2249

50. Green, E. R., and Mecsas, J. (2016) Bacterial Secretion Systems: An Overview. Microbiol. Spectr. 4, 4.1.13

51. Tsirigotaki, A., De Geyter, J., Šoštarić, N., Economou, A., and Karamanou, S. (2017) Protein export through the bacterial Sec pathway. Nat. Rev. Microbiol. 15, 21–36

52. Qin, H., Lo, N. W.-S., Loo, J., Lin, X., Yim, A. K.-Y., Tsui, S. K.-W., Lau, T. C.-K., Ip, M., and Chan, T.-F. (2018) Comparative transcriptomics of multidrug-resistant Acinetobacter baumannii in response to antibiotic treatments. Sci. Rep. 8, 3515

53. Hua, X., Liu, L., Fang, Y., Shi, Q., Li, X., Chen, Q., Shi, K., Jiang, Y., Zhou, H., and Yu, Y. (2017) Colistin Resistance in Acinetobacter baumannii MDR-ZJ06 Revealed by a Multiomics Approach. Front. Cell. Infect. Microbiol. 7, 45

54. Giuffrè, A., Borisov, V. B., Arese, M., Sarti, P., and Forte, E. (2014) Cytochrome bd oxidase and bacterial tolerance to oxidative and nitrosative stress. Biochim. Biophys. Acta BBA - Bioenerg. 1837, 1178–1187

55. Kashyap, S., Sharma, P., and Capalash, N. (2022) Tobramycin Stress Induced Differential Gene Expression in Acinetobacter baumannii. Curr. Microbiol. 79, 88

56. Wick, R. R., Judd, L. M., Gorrie, C. L., and Holt, K. E. (2017) Unicycler: Resolving bacterial genome assemblies from short and long sequencing reads. PLOS Comput. Biol. 13, e1005595

57. Tatusova, T., DiCuccio, M., Badretdin, A., Chetvernin, V., Nawrocki, E. P., Zaslavsky, L., Lomsadze, A., Pruitt, K. D., Borodovsky, M., and Ostell, J. (2016) NCBI prokaryotic genome annotation pipeline. Nucleic Acids Res. 44, 6614–6624

58. Wick, R. R., Schultz, M. B., Zobel, J., and Holt, K. E. (2015) Bandage: interactive visualization of de novo genome assemblies. Bioinformatics. 31, 3350–3352

59. Simão, F. A., Waterhouse, R. M., Ioannidis, P., Kriventseva, E. V., and Zdobnov, E. M. (2015) BUSCO: assessing genome assembly and annotation completeness with single-copy orthologs. Bioinformatics. 31, 3210–3212

60. Minh, B. Q., Schmidt, H. A., Chernomor, O., Schrempf, D., Woodhams, M. D., von Haeseler, A., and Lanfear, R. (2020) IQ-TREE 2: New Models and Efficient Methods for Phylogenetic Inference in the Genomic Era. Mol. Biol. Evol. 37, 1530–1534

61. Letunic, I., and Bork, P. (2024) Interactive Tree of Life (iTOL) v6: recent updates to the phylogenetic tree display and annotation tool. Nucleic Acids Res. 52, W78–W82

62. Alcock, B. P., Raphenya, A. R., Lau, T. T. Y., Tsang, K. K., Bouchard, M., Edalatmand, A., Huynh, W., Nguyen, A.-L. V., Cheng, A. A., Liu, S., Min, S. Y., Miroshnichenko, A., Tran, H.-K., Werfalli, R. E., Nasir, J. A., Oloni, M., Speicher, D. J., Florescu, A., Singh, B., Faltyn, M., Hernandez-Koutoucheva, A., Sharma, A. N., Bordeleau, E., Pawlowski, A. C., Zubyk, H. L., Dooley, D., Griffiths, E., Maguire, F., Winsor, G. L., Beiko, R. G., Brinkman, F. S. L., Hsiao, W. W. L., Domselaar, G. V., and McArthur, A. G. (2019) CARD 2020: antibiotic resistome surveillance with the comprehensive antibiotic resistance database. Nucleic Acids Res. 10.1093/nar/gkz935

63. Ross, K., Varani, A. M., Snesrud, E., Huang, H., Alvarenga, D. O., Zhang, J., Wu, C., McGann, P., and Chandler, M. (2021) TnCentral: a Prokaryotic Transposable Element Database and Web Portal for Transposon Analysis. mBio. 12, 10.1128/mbio.02060-21

64. Siguier, P., Perochon, J., Lestrade, L., Mahillon, J., and Chandler, M. (2006) ISfinder: the reference centre for bacterial insertion sequences. Nucleic Acids Res. 34, D32–36

65. Babraham Bioinformatics - FastQC A Quality Control tool for High Throughput Sequence Data [online] https://www.bioinformatics.babraham.ac.uk/projects/fastqc/ (Accessed February 15, 2025)

66. Ewels, P., Magnusson, M., Lundin, S., and Käller, M. (2016) MultiQC: summarize analysis results for multiple tools and samples in a single report. Bioinformatics. 32, 3047–3048

67. Langmead, B., and Salzberg, S. L. (2012) Fast gapped-read alignment with Bowtie 2. Nat. Methods. 9, 357–359

68. Danecek, P., Bonfield, J. K., Liddle, J., Marshall, J., Ohan, V., Pollard, M. O., Whitwham, A., Keane, T., McCarthy, S. A., Davies, R. M., and Li, H. (2021) Twelve years of SAMtools and BCFtools. GigaScience. 10, giab008

69. Liao, Y., Smyth, G. K., and Shi, W. (2014) featureCounts: an efficient general purpose program for assigning sequence reads to genomic features. Bioinformatics. 30, 923–930

70. Love, M. I., Huber, W., and Anders, S. (2014) Moderated estimation of fold change and dispersion for RNA-seq data with DESeq2. Genome Biol. 15, 550

71. Kanehisa, M., and Goto, S. (2000) KEGG: kyoto encyclopedia of genes and genomes. Nucleic Acids Res. 28, 27–30

