## Supplementary material for "The peptide LyeTx I mnΔK induces transcriptomic reprogramming in a novel Multidrug-resistant *Acinetobacter baumannii*": Supplemetal Figures

*Running title: A. baumannii* Response to LyeTx I mnΔK Peptide Treatment

*Corresponding Author:

Maria E. de Lima:

Glória R. Franco:

**Table of Contents**

**Table S1.** Genomic features of *A. baumannii* AC37 strain
**Table S2.** Accession numbers for all *Acinetobacter* genomes used at NCBI
**Table S3.** Resistance genes in *A. baumannii* AC37 strain: antibiotic classes and mechanisms
**Table S4.** Differentially expressed genes in the LyeTx I mnΔK treatment
**Table S5.** Differentially expressed genes in the combined treatment withLyeTx I mnΔK and meropenem
**Table S6.** Characteristics of studies of *Acinetobacter baumannii* treated with Meropenem deposited in the SRA.
**Table S7.** Differentially expressed genes in the meropenem treatment of BioProject PRJNA787205
**Figure S1.** Visualization of the genome assembled by Unicycler using Bandage.
**Figure S2.** Growth curves of isolate AC37 under sub-MIC conditions in the presence of peptide LyeTx I mnΔK.
**Figure S3.** Growth curves of isolate AC37 under sub-MIC conditions in the presence of peptide LyeTx I mnΔK and the antibiotic Meropenem.
Figure S4. Sample variation profile.
**Figure S5.** Volcano plots from the PRJNA787205 study
**Figure S6.** Enriched biological processes in Meropenem treatments from the PRJNA787205 project.

**Table S1: Genomic features of *A. baumannii* AC37 strain**

| **Features** | ***A. baumannii* AC37** |
|---|---|
| Number of reads (2x301) | 5,117,651 |
| Genome size main chromosome | 3,843,875 bp |
| Plasmid NZ_CP131945.1 | 8,731 bp |
| Average Coverage | 750x |
| GC | 39% |
| Genes (total) | 3716 |
| CDS (total) | 3646 |
| Genes (coding) | 3588 |
| Genes (RNA) | 70 |
| rRNAs | 1 5S, 1 16S, 1 23S |
| tRNAs | 63 |
| ncRNAs | 4 |
| Pseudo Genes (total) | 58 |

**Table S2: Accession numbers for all *Acinetobacter* genomes used at NCBI**

| **Accession number** | ***Acinetobacter* strain** |
|---|---|
| GCF_041021905.1 (reference) | *A. nosocomialis* XH1679 |
| GCF_009035845.1 (reference) | *A. baumannii* ATCC 19606 |
| GCF_041154855.1 | *A. baumannii* JUNP543 |
| GCF_038987955.1 | *A. baumannii* SRM25 |
| GCF 041014305.1 | *A. baumannii* AB37-AUFP |
| GCF_036718155.1 | *A. baumannii* 2024CK-00130 |
| GCF 038441325.1 | *A. baumannii* CRAb2 |
| GCF 038442125.1 | *A. baumannii* Ab1 |
| GCF 038442095.1 | *A. baumannii* CRAb1 |
| GCF 038627335.1 | *A. baumannii* 2024CK-00546 |
| GCF 037008745.1 | *A. baumannii* 2024CK-00246 |
| GCF_041801535.1 | *A. baumannii* Y03 |
| GCF_041154805.1 | *A. baumannii* JUNP406 |
| GCF_040822435.1 | *A. baumannii* SIMBA061 |
| GCF_040930415.1 | *A. baumannii* SIMBA113 |
| GCF_040822385.1 | *A. baumannii* SIMBA035 |
| GCF_041154835.1 | *A. baumannii* JUNP499 |
| GCF_040822395.1 | *A. baumannii* SIMBA028 |
| GCF_040822125.1 | *A. baumannii* SIMBA003 |
| GCF_040357515.1 | *A. baumannii* XH2146 |
| GCF_040543095.1 | *A. baumannii* AB11-AUFP |
| GCF_040703085.1 | *A. baumannii* AB44-AUFP |
| GCF_041154825.1 | *A. baumannii* JUNP496 |
| GCF_038990075.1 | *A. baumannii* SRM3 |
| GCF 041154845.1 | *A. baumannii* JUNP514 |
| GCA_041154865.1 | *A. baumannii* JUNP586 |
| GCF 040834015.1 | *A. baumannii* SIMBA089 |
| GCF 041154875.1 | *A. baumannii* JUNP712 |
| GCF 041154775.1 | *A. baumannii* JUNP402 |
| GCF 038441195.1 | *A. baumannii* 2008S11-069 |
| GCF 038441235.1 | *A. baumannii* 2008N08-156 |
| GCF 038441245.1 | *A. baumannii* 2010C01-165 |
| GCF_037008665.1 | *A. baumannii* 2024CK-00250 |
| GCF_039654205.1 | *A. baumannii* 2024CK-00247 |
| GCF_034331205.1 | *A. baumannii* 2023CK-01512 |
| GCF_038418365.1 | *A. baumannii* 2024CK-00462 |
| GCF_041154795.1 | *A. baumannii* JUNP405 |
| GCF_041154785.1 | *A. baumannii* JUNP403 |
| GCF_040822405.1 | *A. baumannii* SIMBA034 |
| GCF_041154815.1 | *A. baumannii* JUNP419 |
| GCF_038985835.1 | *A. baumannii* SRM21 |
| GCF_038069255.1 | *A. baumannii* 2024CK-00376 |
| GCF_038995905.1 | *A. baumannii* YFY27 |
| GCF_038441255.1 | *A. baumannii* 2010N17-053 |
| GCF_038441215.1 | *A. baumannii* 2008C02-166 |
| GCF_034126645.1 | *A. baumannii* 2023CK-01487 |
| GCF_034126905.1 | *A. baumannii* 2023CK-01486 |
| GCF_038441225.1 | *A. baumannii* 2010N16-194 |

| | |
|---|---|
| GCF_038441205.1 | *A. baumannii* 2006S08-082 |
| GCF_038992355.1 | *A. baumannii* YFY21 |
| GCF_038627275.1 | *A. baumannii* 2024CK-00536 |
| GCF_038627255.1 | *A. baumannii* 2024CK-00538 |
| GCF_038522965.1 | *A. baumannii* 2024CK-00537 |
| GCF_038418265.1 | *A. baumannii* 2024CK-00452 |
| GCF_038069235.1 | *A. baumannii* 2024CK-00375 |
| GCF_041004195.1 | *A. baumannii* W155 |
| GCF_038998165.1 | *A. baumannii* YFY3 |
| GCF_038983805.1 | *A. baumannii* SRM1 |
| GCF_039518815.1 | *A. baumannii* 2024CK-00319 |
| GCF_039566025.1 | *A. baumannii* Aci44 |
| GCF_038993615.1 | *A. baumannii* YFY24 |
| GCA_040956925.1 | *A. baumannii* MTC0629 |
| GCA_040956955.1 | *A. baumannii* MTC0620 |
| GCA_040957165.1 | *A. baumannii* MTC0609 |
| GCA_040956905.1 | *A. baumannii* MTC1120 |
| GCF_037939845.1 | *A. baumannii* AB7276 |
| GCF_035666135.1 | *A. baumannii* EMB-1 |
| GCF_029847455.2 | *A. baumannii* PUMA0052 |
| GCF_038442235.1 | *A. baumannii* F19-10-CH8 |
| GCF_038442275.1 | *A. baumannii* F19-10-CH3 |
| GCF_038442305.1 | *A. baumannii* F19-10-9 |
| GCF_038442395.1 | *A. baumannii* F19-10-2 |
| GCF_038442375.1 | *A. baumannii* F19-10-8 |
| GCF_038442575.1 | *A. baumannii* F15-01B |
| GCF_038442265.1 | *A. baumannii* F19-10-CH6 |
| GCF_038442565.1 | *A. baumannii* F15-02 |
| GCF_038442555.1 | *A. baumannii* F15-06 |
| GCF_038442585.1 | *A. baumannii* F15-01A |
| GCF_038442385.1 | *A. baumannii* F19-10-4 |
| GCF_038442295.1 | *A. baumannii* F19-10-MH6 |
| GCF_038442285.1 | *A. baumannii* F19-10-MH12 |
| GCF_038442225.1 | *A. baumannii* 40288-CIRI |
| GCF_038442635.1 | *A. baumannii* F14-11 |
| GCF_038442405.1 | *A. baumannii* F19-02 |
| GCF_038442415.1 | *A. baumannii* F18-02 |
| GCF_038442425.1 | *A. baumannii* F16-05 |
| GCF 035937495.1 | *A. baumannii* A38 |
| GCA 036320755.1 | *A. baumannii* C20AB05 |
| GCF 039545565.1 | *A. baumannii* NCSR_106 |
| GCF 036601155.1 | *A. baumannii* AB5075 |
| GCF 036602705.1 | *A. baumannii* AB5075 |
| GCF_039779435.1 | *A. baumannii* D78 |
| GCF_040529295.1 | *A. baumannii* NAB01B-R7 |
| GCF_040536945.1 | *A. baumannii* NAB05B |
| GCF_040529225.1 | *A. baumannii* NAB07B |
| GCF_040529255.1 | *A. baumannii* NAB01B-R5 |
| GCF_040267665.1 | *A. baumannii* NAB02B |
| GCF_040529275.1 | *A. baumannii* NAB01B-R6 |

| | |
|---|---|
| GCF_040529235.1 | *A. baumannii* NAB01B-R1 |
| GCF_040529205.1 | *A. baumannii* NAB03B |
| GCF_040536965.1 | *A. baumannii* NAB01B-R1-2 |
| GCF_040529215.1 | *A. baumannii* NAB04B |
| GCF_040529305.1 | *A. baumannii* NAB01B-R9 |
| GCF_040529245.1 | *A. baumannii* NAB01B-R3 |
| GCF_040536955.1 | *A. baumannii* NAB06B |
| GCF_040529325.1 | *A. baumannii* NAB01B |
| GCF_040529285.1 | *A. baumannii* NAB01B-R8 |
| GCF_035754475.1 | *A. baumannii* C20AB01 |
| GCF_037039675.1 | *A. baumannii* G20AB08 |
| GCF_035753605.1 | *A. baumannii* A20AB13 |
| GCF_036870965.1 | *A. baumannii* A20AB02 |
| GCF_033378915.2 | *A. baumannii* SCCH68:Ab991128 |
| GCF_036287275.1 | *A. baumannii* B20AB10 |
| GCF_036287255.1 | *A. baumannii* B20AB01 |
| GCF_040529315.1 | *A. baumannii* NAB01B-R10 |
| GCF_040529265.1 | *A. baumannii* NAB01B-R4 |
| GCF_037215885.1 | *A. baumannii* LZFZ3604 |
| GCF_028548795.2 | *A. baumannii* AB191 |
| GCF_039701065.1 | *A. baumannii* TY918 |
| GCF_040536855.1 | *A. baumannii* Ab6-Co-2 |
| GCA_035755505.1 | *A. baumannii* B20AB06 |
| GCA_037039585.1 | *A. baumannii* F20AB03 |
| GCF 039780165.1 | *A. baumannii* D13 |

**Table S3: Resistance genes in *A. baumannii* AC37 strain: antibiotic classes and mechanisms**

| Gene | Drug Class | Antibiotic Resistance | Resistance Mechanism |
|---|---|---|---|
| AAC(6')-Ib' | Aminoglycosides | Amikacin and sobramycin | Aminoglycoside-modifying enzyme |
| AbaF | Phosphonic acid | Fosfomycin | Efflux pump |
| AbaQ | Fluoroquinolones | - | Efflux pump |
| abeM | Fluoroquinolones, disinfectants and antiseptics | Spectinomycin and streptomycin | Efflux pump |
| abeS | Macrolites and aminocoumarins | Erythromycin and novobiocin | Efflux pump |
| ADC-183 | Cephalosporin | - | β-lactamase |
| adeA | Glycylcyclices and tetracyclines | Tigecycline and tetracycline | Efflux pump |
| adeB | Glycylcyclices and tetracyclines | Tigecycline and tetracycline | Efflux pump |
| adeC | Glycylcyclices and tetracyclines | Tigecycline and tetracycline | Efflux pump |
| adeF | Fluoroquinolones and tetracyclines | Tetracycline | Efflux pump |
| adeG | Fluoroquinolones and tetracyclines | Tetracycline | Efflux pump |
| adeH | Fluoroquinolones and tetracyclines | Tetracycline | Efflux pump |
| adeI | Macrolides, fluoroquinolones, lincosamides, carbapenems, cephalosporins, tetracyclines, rifamycins, diaminopyrimidines and phenicols | Tetracycline, rifampicin, imipenem, trimethoprim, chloramphenicol and ticarcillin | Efflux pump |

| | | | |
|---|---|---|---|
| adeJ | Macrolides, fluoroquinolones, lincosamides, carbapenems, cephalosporins, tetracyclines, rifamycins, diaminopyrimidines and phenicols | Tetracycline, rifampicin, imipenem, trimethoprim, chloramphenicol and ticarcillin | Efflux pump |
| adeK | Macrolides, fluoroquinolones, lincosamides, carbapenems, cephalosporins, tetracyclines, rifamycins, diaminopyrimidines and phenicols | Tetracycline, rifampicin, imipenem, trimethoprim, chloramphenicol and ticarcillin | Efflux pump |
| adeL | Fluoroquinolones and tetracyclines | Tetracycline | adeFGH regulator |
| adeN | Macrolides, fluoroquinolones, lincosamides, carbapenems, cephalosporins, tetracyclines, rifamycins, diaminopyrimidines and phenicols | Tetracycline, rifampicin, imipenem, trimethoprim, chloramphenicol and ticarcillin | adeIJK regulator |
| adeR | Glycylcyclices and tetracyclines | Tigecycline and tetracycline | adeABC regulator |
| adeS | Glycylcyclices and tetracyclines | Tigecycline and tetracycline | adeABC regulator |
| AmvA | Macrolides, disinfectants and antiseptics | Erythromycin and acriflavine | Efflux pump |
| ANT(3’’)-llc | Aminoglycosides | Spectinomycin and streptomycin | Aminoglycoside-modifying enzime |
| APH(3’’)-lb | Aminoglycosides | Streptomycin | Aminoglycoside-modifying enzime |
| APH(6)-id | Aminoglycosides | Streptomycin | Aminoglycoside-modifying enzime |
| floR | Phenicols | Chloramphenicol and florfenicol | Efflux pump |

| | | | |
|---|---|---|---|
| gyrA | Fluoroquinolones | Enoxacin, ciprofloxacin, levofloxacin, moxifloxacion, gatifloxacin, lomefloxacin, nalidixic acid, norfloxacin, ofloxacin, trovafloxacin, grepafloxacin, sparfloxacin and perfloxacin | Structural alteration in DNA gyrase |
| LpsB | Peptides | Colistin A, colistin B and defesins | Mutation in outer membrane proteins |
| OXA-23 | Carbapems, cephalosporins | Cloxacilin, oxacilin and cephalothin | β-lactamase |
| OXA-63 | Carbapems, cephalosporins | Cloxacilin, oxacilin and cephalothin | β-lactamase |
| parC | Fluoroquinolones | Enoxacin, ciprofloxacin, levofloxacin, moxifloxacin, gatifloxacin, lomefloxacin, nalidixic acid, norfloxacin, ofloxacin, trovafloxacin, grepafloxacin, sparfloxacin and perfloxacin | Structural alteration in Topoisomerase IV |
| rsmA | Fluoroquinolones, diaminopyrimidines and phenicols | Trimethoprim and chloramphenicol | Efflux pump |
| sul2 | Sulfonamides | Sulfadiazine, sulfadimidine, | - |

sulfamethoxazole, fulfisoxazole,
sulfacetamide, mefenide,
sulfasalazine and sulfamethizole

**Table S4. Differentially expressed genes in the LyeTx I mnΔK treatment**

**Table S5. Differentially expressed genes in the combined treatment with LyeTx I mnΔK and meropenem**

**Table S6. Characteristics of studies of *Acinetobacter baumannii* treated with Meropenem deposited in the SRA.** In bold the identifier of the selected studies

| Bioproject | Strain | Read Length | Meropenem Concentration | Growth Time After Treatment | Library Depth | Sequencing Platform | Read Type | Number of Replicates |
|---|---|---|---|---|---|---|---|---|
| PRJNA321935 | R1 to R9 (resistent), S1 to S3 (sensible) | 101 pb | sub-MIC (1/8 to 1/64 x MIC) | - | 17-26M pb | Illumina HiSeq 2000 | Paired-end | Duplicate |
| PRJNA518730 | ATCC 17978 (resistent) | 68 pb | MIC | 10, 30 and 60 min | 8-21M pb | Illumina HiSeq2500 | Paired-end | Duplicate |
| PRJNA548006 | 37662 (resistent) | 150 pb | sub-MIC (0.5 x MIC) | 30 days | 10-12M pb | Illumina HiSeq2500 | Paired-end | Triplicate |
| PRJNA686448 | ATCC 17978 (resistent) | 58 pb | - | 30 and 90 min | 3-9M pb | Illumina NextSeq 500 | Single-end | Triplicate |
| **PRJNA787205** | **ATCC 17978 (resistent)** | **50 pb** | **MIC** | **30 min, 3h and 9h** | **10-16M pb** | **Illumina NextSeq 550** | **Paired-end** | **Triplicate** |
| PRJNA797559 | AB5116 (sensible) | 150 pb | sub-MIC (0.5 x MIC, 0.0625 µg/mL) | 30 min | 10-14M pb | Illumina HiSeq X Ten | Paired-end | Triplicate |
| PRJNA876061 | AB075 (resistent) | 102 pb | 25, 100 and 200 µg/mL | 15 min | 1-3M pb | Illumina HiSeq 4000 | Paired-end | Duplicate |

**Table S7. Differentially expressed genes in the meropenem treatment of BioProject PRJNA787205**

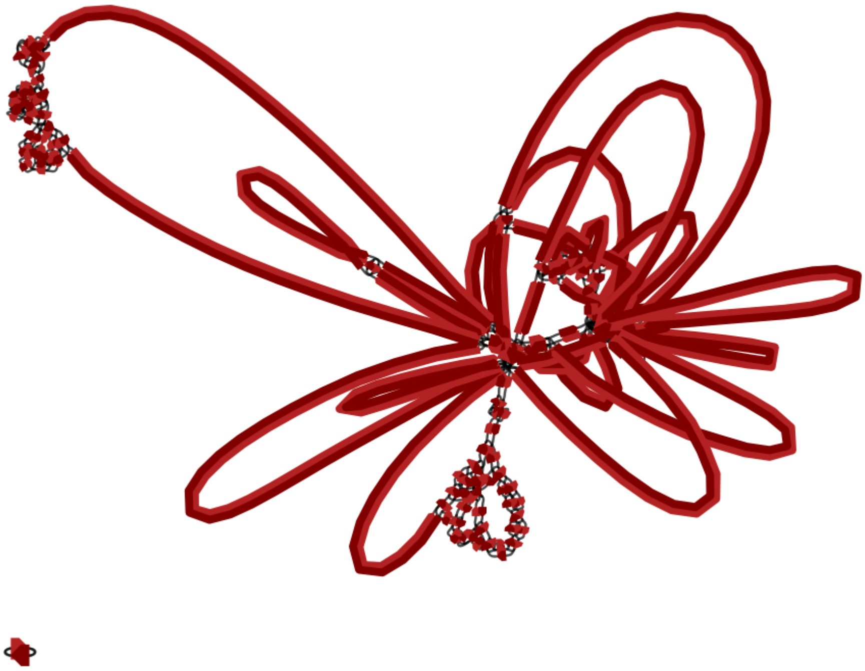

**Figure S1. Visualization of the genome assembled by Unicycler using Bandage.** Unicycler attempts to form bridges to link contigs and circularize the bacterial genome. Bandage aids in visualizing the assembled genome and identifying extrachromosomal structures, such as plasmids.

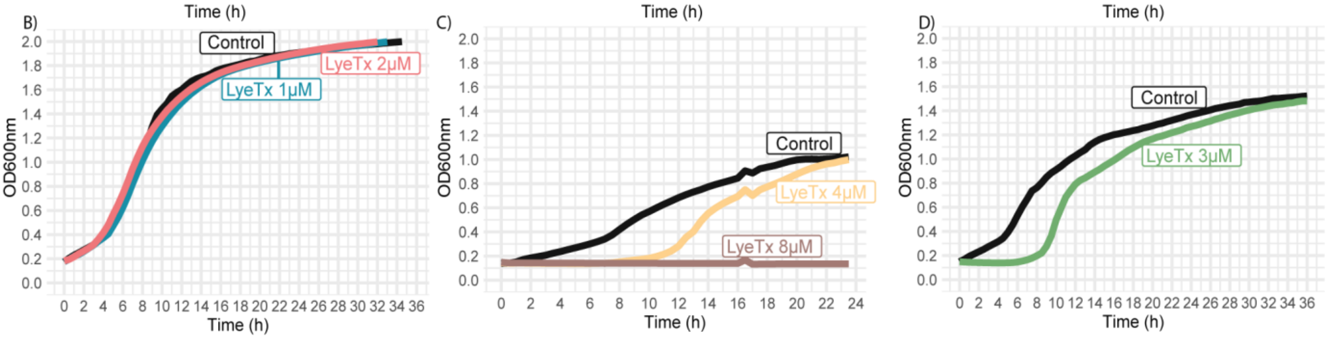


**Figure S2. Growth curves of isolate AC37 under sub-MIC conditions in the presence of peptide LyeTx I mnΔK.** Experiments conducted by collaborators. The selected concentration for cultivation and RNA-seq library preparation was 3 μM. MIC of LyeTx I mnΔK = 4 μM.

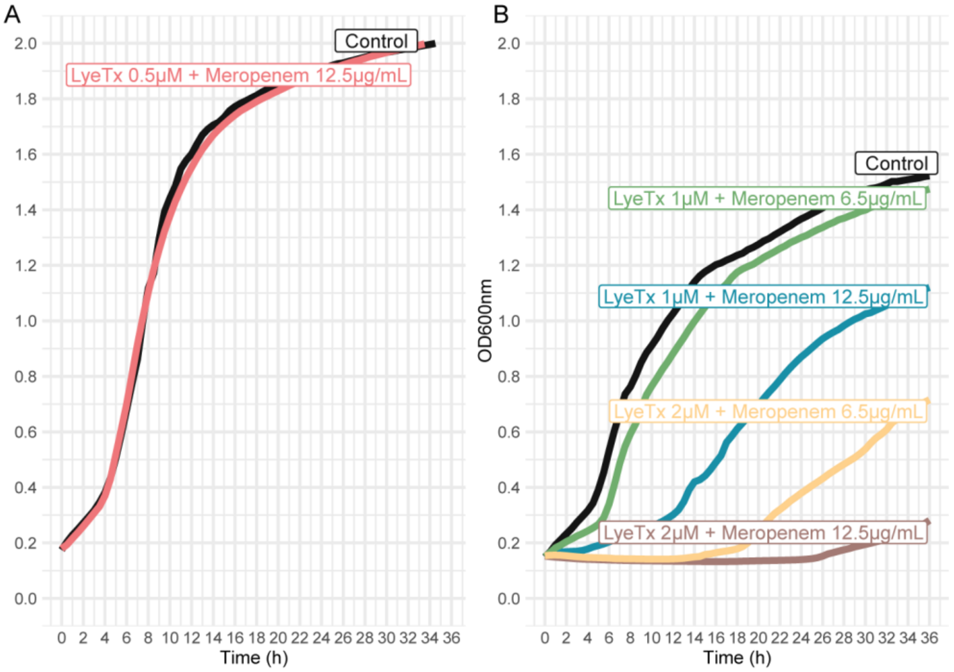


**Figure S3. Growth curves of isolate AC37 under sub-MIC conditions in the presence of peptide LyeTx I mnΔK and the antibiotic Meropenem.** Experiments performed by collaborators. The selected concentrations for cultivation and RNA-seq library preparation were 1 μM of the peptide and 6.5 μg/mL of Meropenem. MIC values: LyeTx I mnΔK = 4 μM, Meropenem ≥ 16 μg/mL

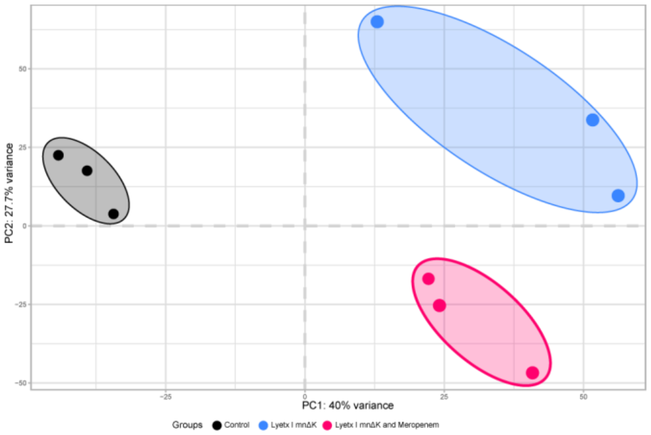


**Figure S4. Sample variation profile.** Biplot showing the results of the principal component analysis performed with the *prcomp* function in R. Each circle represents a sample from each experimental condition. Controls are shown in black, LyeTx I mnΔK in blue, and the synergistic effect in pink.

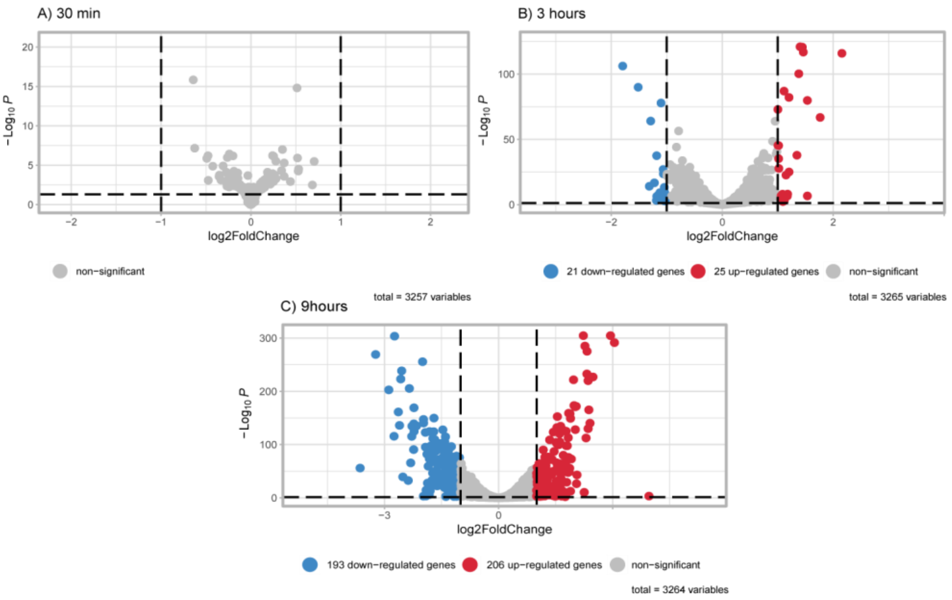


**Figure S5. Volcano plots from the PRJNA787205 study.** Representation of the number of differentially expressed genes identified. Blue, red, and gray circles represent downregulated, upregulated, and non-differentially expressed genes, respectively, based on an absolute log2FoldChange ≥ 1 and adjusted p-value < 0.05.

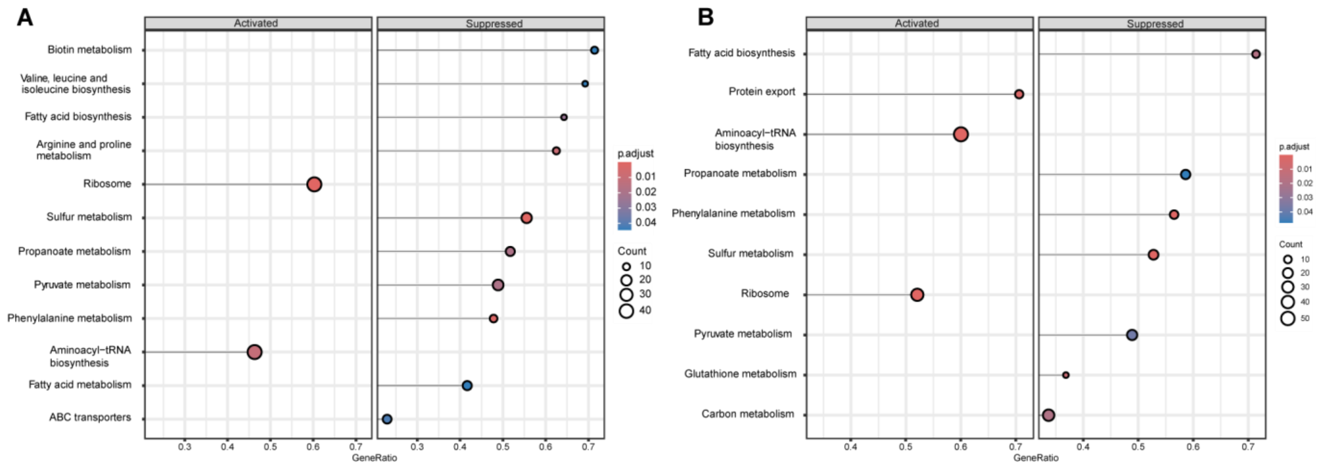


**Figure S6. Enriched biological processes in Meropenem treatments from the PRJNA787205 project.** Lollipop plot of activated and suppressed pathways under treatment conditions using the gseKEGG function. Color spectrum represents the significance value (adjusted p-value) of biological pathways. The size of each circle indicates the count of enriched genes in each pathway. Panels (A) and (B) represent Meropenem incubation times of 3 hours and 9 hours, respectively.
