## Supplemental table 4 for "The peptide LyeTx I mnΔK induces transcriptomic reprogramming in a novel Multidrug-resistant *Acinetobacter baumannii*"

| | pgap_annot | log2FoldChange | pvalue | gene_name |
|---|---|---|---|---|
| 1 | cds-pgaptmp_000039 | 1,19587939208073 | 2,65744445602524e-15 | NA |
| 2 | cds-pgaptmp_000040 | 1,09655014785856 | 1,08833391743684e-28 | NA |
| 3 | cds-pgaptmp_000053 | -1,10088190915218 | 9,65567568826493e-14 | NA |
| 4 | cds-pgaptmp_000054 | 1,35583225009182 | 2,24624124920354e-49 | putP |
| 5 | cds-pgaptmp_000068 | -1,05191914808406 | 1,76618875637058e-35 | NA |
| 6 | cds-pgaptmp_000084 | -5,05419950771527 | 0 | NA |
| 7 | cds-pgaptmp_000085 | -4,00560298811504 | 2,46562790428069e-242 | NA |
| 8 | cds-pgaptmp_000093 | 1,60168816910134 | 1,35035185177673e-39 | NA |
| 9 | cds-pgaptmp_000094 | 1,05422232132263 | 2,54040269995567e-31 | NA |
| 10 | cds-pgaptmp_000097 | -2,06063104255788 | 5,77614026979504e-56 | NA |
| 11 | cds-pgaptmp_000098 | -2,05937017576753 | 5,45746867964088e-45 | NA |
| 12 | cds-pgaptmp_000099 | -2,11645274488162 | 1,216029553431e-60 | NA |
| 13 | cds-pgaptmp_000100 | -2,19525860899942 | 6,12794911805897e-130 | NA |
| 14 | cds-pgaptmp_000109 | 1,37744080370776 | 2,14778150548547e-43 | ychF |
| 15 | cds-pgaptmp_000125 | -1,92370963488083 | 5,52857919662869e-77 | NA |
| 16 | cds-pgaptmp_000140 | -1,14058253494748 | 2,03604300648327e-15 | NA |
| 17 | cds-pgaptmp_000149 | 1,47764778842673 | 2,95530183872476e-18 | tauA |
| 18 | cds-pgaptmp_000154 | -1,38382270038566 | 3,55002275444942e-18 | NA |
| 19 | cds-pgaptmp_000155 | -2,03510029712553 | 1,6019426087035e-86 | NA |
| 20 | cds-pgaptmp_000156 | -2,07615437273278 | 1,08441049237496e-89 | cydB |
| 21 | cds-pgaptmp_000157 | -2,16616420243514 | 5,28735557338262e-25 | NA |
| 22 | cds-pgaptmp_000166 | -1,06433781012277 | 8,17286690851513e-05 | NA |
| 23 | cds-pgaptmp_000167 | -1,41858802607635 | 5,78685231562968e-15 | NA |
| 24 | cds-pgaptmp_000168 | -1,65248086505549 | 1,28285055368572e-25 | mdcA |
| 25 | cds-pgaptmp_000174 | 1,4447954693789 | 0,000300227461601996 | NA |
| 26 | cds-pgaptmp_000176 | -1,38980527448746 | 1,01008277917763e-33 | yfcF |
| 27 | cds-pgaptmp_000177 | -1,47631136349322 | 2,35187913888489e-53 | yghU |
| 28 | cds-pgaptmp_000190 | 1,05631171356643 | 1,45997684889055e-05 | NA |
| 29 | cds-pgaptmp_000191 | 2,17619171588552 | 6,40001360570384e-11 | NA |
| 30 | cds-pgaptmp_000192 | 1,10966157640874 | 1,50145275458017e-06 | NA |
| 31 | cds-pgaptmp_000199 | -1,33312454987694 | 1,44710778122367e-17 | NA |
| 32 | cds-pgaptmp_000202 | -3,34967903006486 | 5,44439262065075e-162 | NA |
| 33 | cds-pgaptmp_000203 | -2,76243836861445 | 2,03685665261385e-15 | NA |
| 34 | cds-pgaptmp_000204 | -4,404890294486 | 0 | NA |
| 35 | cds-pgaptmp_000205 | -4,05088811164666 | 0 | katE |
| 36 | cds-pgaptmp_000206 | -3,77741768068886 | 5,18177532962291e-229 | NA |
| 37 | cds-pgaptmp_000207 | -2,88443037248375 | 1,60982154386109e-12 | NA |
| 38 | cds-pgaptmp_000208 | -5,38297794641797 | 4,53733810166529e-50 | NA |
| 39 | cds-pgaptmp_000209 | -4,75150407017122 | 3,62816432754348e-267 | surA1 |
| 40 | cds-pgaptmp_000210 | -2,17055747094527 | 4,47571258252326e-32 | NA |
| 41 | cds-pgaptmp_000211 | -3,17816720136068 | 9,07353054470437e-92 | NA |
| 42 | cds-pgaptmp_000212 | -3,90812835271703 | 4,97318286167281e-87 | NA |
| 43 | cds-pgaptmp_000216 | -1,59614786161372 | 3,96348991863215e-10 | NA |
| 44 | cds-pgaptmp_000217 | -3,57040609097344 | 5,40276211972561e-74 | NA |
| 45 | cds-pgaptmp_000218 | -3,67148272472501 | 7,06419489075637e-176 | NA |
| 46 | cds-pgaptmp_000219 | -3,21541582370166 | 1,87596466596837e-198 | NA |
| 47 | cds-pgaptmp_000220 | -3,12022904939372 | 1,63004627580644e-180 | NA |
| 48 | cds-pgaptmp_000221 | -3,01836844104179 | 1,21029245492535e-153 | NA |

| | | | | |
|---|---|---|---|---|
| 49 | cds-pgaptmp_000222 | -3,02883079399572 | 1,53072968479167e-139 | NA |
| 50 | cds-pgaptmp_000223 | -2,9681091445778 | 8,81763160263602e-150 | NA |
| 51 | cds-pgaptmp_000224 | -2,77222916236483 | 2,6771075713357e-119 | NA |
| 52 | cds-pgaptmp_000226 | -3,3669118689971 | 9,66490359367887e-230 | NA |
| 53 | cds-pgaptmp_000271 | -1,36623292410477 | 2,63544125537185e-22 | NA |
| 54 | cds-pgaptmp_000272 | -1,33551548651402 | 1,10012835069998e-29 | NA |
| 55 | cds-pgaptmp_000278 | -2,52258024090713 | 6,83874841491818e-09 | NA |
| 56 | cds-pgaptmp_000279 | -3,09319349695725 | 8,02734906336347e-29 | NA |
| 57 | cds-pgaptmp_000280 | -3,11731027116855 | 1,1680255003118e-33 | NA |
| 58 | cds-pgaptmp_000281 | -2,18863059423983 | 1,93725930991197e-20 | NA |
| 59 | cds-pgaptmp_000282 | -2,50324343749196 | 3,276568333424e-10 | NA |
| 60 | cds-pgaptmp_000283 | -1,62922701029078 | 1,03610680238669e-07 | NA |
| 61 | cds-pgaptmp_000284 | -1,48527177500385 | 8,43705213226885e-20 | NA |
| 62 | cds-pgaptmp_000286 | -1,81205777698789 | 4,54591351543442e-25 | NA |
| 63 | cds-pgaptmp_000290 | -2,02415470853193 | 1,4136677618324e-07 | NA |
| 64 | cds-pgaptmp_000291 | -1,9136356861039 | 8,20094714949009e-63 | NA |
| 65 | cds-pgaptmp_000292 | -1,56595220121691 | 1,37578328422106e-59 | paaX |
| 66 | cds-pgaptmp_000293 | -3,03601483385162 | 1,18172141897307e-191 | paaK |
| 67 | cds-pgaptmp_000294 | -3,82621696390308 | 1,30804174505646e-273 | pcaF |
| 68 | cds-pgaptmp_000295 | -4,45622842073135 | 1,63724896641021e-289 | NA |
| 69 | cds-pgaptmp_000296 | -5,12284739282986 | 1,34770295853543e-248 | paaG |
| 70 | cds-pgaptmp_000297 | -5,29513110927012 | 1,38722917576412e-202 | NA |
| 71 | cds-pgaptmp_000298 | -4,87270582861026 | 3,27723713862199e-289 | paaE |
| 72 | cds-pgaptmp_000299 | -4,60521994646234 | 7,19111917848856e-167 | paaD |
| 73 | cds-pgaptmp_000300 | -4,74035917913701 | 1,07546477519442e-37 | paaC |
| 74 | cds-pgaptmp_000301 | -4,70784665109864 | 5,61790620202347e-181 | paaB |
| 75 | cds-pgaptmp_000302 | -4,59865360586642 | 5,92946859151377e-39 | paaA |
| 76 | cds-pgaptmp_000303 | -4,28030369248253 | 2,69680424600712e-68 | paaZ |
| 77 | cds-pgaptmp_000311 | -1,28859888621718 | 3,41964231169035e-06 | NA |
| 78 | cds-pgaptmp_001410 | 1,22162681636446 | 4,37880856570257e-36 | NA |
| 79 | cds-pgaptmp_001411 | 1,17845334374959 | 6,34799032624036e-34 | lpxO |
| 80 | exon-pgaptmp_001435-1 | 1,03038231874226 | 9,45555645387109e-25 | NA |
| 81 | cds-pgaptmp_001441 | 1,21911967025291 | 1,15686094485e-36 | NA |
| 82 | cds-pgaptmp_001448 | -1,07224328967225 | 1,14725313517349e-33 | NA |
| 83 | cds-pgaptmp_001465 | -1,66372637882448 | 5,14182396065949e-42 | NA |
| 84 | cds-pgaptmp_001466 | -1,26571733939785 | 1,29752551107475e-27 | NA |
| 85 | cds-pgaptmp_001486 | 1,09803252277657 | 6,73586687241335e-20 | NA |
| 86 | exon-pgaptmp_001501-1 | 1,30344165531994 | 9,02817475491662e-16 | NA |
| 87 | cds-pgaptmp_001502 | 1,00298878487242 | 8,23158556168553e-29 | NA |
| 88 | cds-pgaptmp_001508 | -1,18891097169903 | 3,12127408413732e-25 | NA |
| 89 | cds-pgaptmp_001548 | -1,82488378376633 | 7,59734964136511e-82 | NA |
| 90 | cds-pgaptmp_001550 | 1,33074114814726 | 1,73334069534794e-35 | rpmG |
| 91 | cds-pgaptmp_001551 | 1,32629586586242 | 1,16665650686484e-33 | rpmB |
| 92 | cds-pgaptmp_001552 | -2,31337955045975 | 1,44481277402591e-100 | NA |
| 93 | cds-pgaptmp_001576 | -1,35827375354884 | 1,71838504662455e-55 | leuA |
| 94 | cds-pgaptmp_001578 | 1,36659673734622 | 1,30635905844171e-24 | NA |
| 95 | cds-pgaptmp_001579 | 1,42866917772822 | 7,64333129623286e-49 | tig |
| 96 | cds-pgaptmp_001585 | -1,60257994344788 | 5,06717062652072e-48 | pta |
| 97 | cds-pgaptmp_001586 | -1,69924322401386 | 1,79942445822901e-41 | NA |

| | | | | |
|---|---|---|---|---|
| 98 | cds-pgaptmp_001587 | 1,06229288140784 | 7,63827823518597e-39 | edd |
| 99 | cds-pgaptmp_001588 | 1,1366781131264 | 6,79995814449782e-40 | eda |
| 100 | cds-pgaptmp_001589 | 1,18056189303847 | 2,75190413886446e-46 | NA |
| 101 | cds-pgaptmp_001590 | 1,05639015886048 | 1,19484562194185e-28 | NA |
| 102 | cds-pgaptmp_001593 | -1,03280065727031 | 2,01482407708624e-18 | NA |
| 103 | cds-pgaptmp_001598 | -1,96066658705152 | 1,06107622180943e-34 | NA |
| 104 | cds-pgaptmp_001616 | 1,91467759863706 | 1,7801352951871e-57 | NA |
| 105 | cds-pgaptmp_001618 | 1,05322392262388 | 8,78865866393074e-24 | NA |
| 106 | cds-pgaptmp_001619 | 1,3171830136118 | 6,73300544204365e-38 | NA |
| 107 | cds-pgaptmp_001635 | -1,99256460887058 | 2,10668718901962e-85 | acnA |
| 108 | cds-pgaptmp_001637 | 1,79965568566078 | 0,00615687587078157 | NA |
| 109 | cds-pgaptmp_001657 | -2,40315963015188 | 2,07343206495594e-120 | NA |
| 110 | cds-pgaptmp_001660 | -1,86687663587664 | 1,00224669850547e-31 | NA |
| 111 | cds-pgaptmp_001661 | -1,69365751610303 | 6,82348873794215e-73 | NA |
| 112 | cds-pgaptmp_001662 | -1,44356335590623 | 3,00871881298046e-30 | NA |
| 113 | cds-pgaptmp_001668 | -1,50244894232674 | 4,97306205029059e-37 | NA |
| 114 | cds-pgaptmp_001669 | -1,5552952662188 | 4,63550729815133e-27 | NA |
| 115 | cds-pgaptmp_001670 | -1,51181160830903 | 4,88844990346748e-39 | NA |
| 116 | cds-pgaptmp_001675 | -1,16146561957467 | 1,79308702918309e-31 | NA |
| 117 | cds-pgaptmp_001691 | -1,56013278048548 | 1,01877678240151e-29 | NA |
| 118 | cds-pgaptmp_002086 | -2,64603656157985 | 7,06116081070189e-186 | NA |
| 119 | cds-pgaptmp_002101 | 1,04144613065212 | 4,72961505248761e-25 | sdhD |
| 120 | cds-pgaptmp_002105 | 1,37887860508947 | 3,82891748592744e-54 | NA |
| 121 | cds-pgaptmp_002111 | -1,76658467534769 | 5,59513505986762e-88 | NA |
| 122 | cds-pgaptmp_002112 | -1,53884060696332 | 3,82116125578515e-73 | NA |
| 123 | cds-pgaptmp_002121 | 1,1112387944539 | 8,4601562352642e-28 | rpmA |
| 124 | cds-pgaptmp_002122 | 1,06068263320304 | 2,91980160720035e-25 | rplU |
| 125 | cds-pgaptmp_002126 | 2,12029995862092 | 6,2755318920413e-95 | NA |
| 126 | cds-pgaptmp_002127 | 1,66750927388927 | 8,27515109578589e-73 | adeI |
| 127 | cds-pgaptmp_002128 | 1,46338901609779 | 5,31670969932646e-55 | adeJ |
| 128 | cds-pgaptmp_002129 | 1,23826007824668 | 8,92079215041309e-37 | adeK |
| 129 | cds-pgaptmp_002133 | -1,04457846171477 | 9,50102699763952e-28 | NA |
| 130 | cds-pgaptmp_002168 | -1,36616650299605 | 1,34879716049456e-37 | NA |
| 131 | cds-pgaptmp_002177 | -1,09983801136315 | 3,58900939955227e-23 | NA |
| 132 | cds-pgaptmp_002184 | 1,57473850378593 | 4,61130274470901e-40 | NA |
| 133 | cds-pgaptmp_002185 | 1,64801887791905 | 2,0552024131188e-34 | NA |
| 134 | cds-pgaptmp_002186 | 1,66391743449468 | 8,51349753068649e-43 | NA |
| 135 | cds-pgaptmp_002191 | 1,10041696678402 | 2,20018628640707e-24 | NA |
| 136 | cds-pgaptmp_002192 | 1,03122931988899 | 4,03708727791841e-12 | NA |
| 137 | exon-pgaptmp_002193-1 | 1,01922465307976 | 4,82954909942644e-11 | NA |
| 138 | exon-pgaptmp_002194-1 | 1,17106877870097 | 8,83684752649995e-25 | NA |
| 139 | exon-pgaptmp_002195-1 | 1,20973649115726 | 1,24304793568814e-37 | NA |
| 140 | exon-pgaptmp_002196-1 | 1,24852593573547 | 6,92331509785818e-34 | NA |
| 141 | exon-pgaptmp_002197-1 | 1,1093804094641 | 6,90191973022835e-26 | NA |
| 142 | exon-pgaptmp_002198-1 | 1,25002478165574 | 1,39965556565312e-35 | NA |
| 143 | cds-pgaptmp_002202 | -1,94674414822981 | 7,87222157275785e-11 | NA |
| 144 | cds-pgaptmp_002216 | -1,83338067239956 | 3,06205847108262e-72 | NA |
| 145 | cds-pgaptmp_002217 | 1,85320469582013 | 3,34974902911649e-28 | NA |
| 146 | cds-pgaptmp_002219 | -1,94426371714396 | 8,79753405773582e-42 | NA |

| | | | | |
|---|---|---|---|---|
| 147 | cds-pgaptmp_002221 | 1,39028336562073 | 1,33334040443916e-50 | NA |
| 148 | cds-pgaptmp_002230 | 1,09054809960319 | 4,4662672279057e-28 | mscL |
| 149 | cds-pgaptmp_002231 | 1,2709336000128 | 1,93860140408208e-28 | typA |
| 150 | cds-pgaptmp_002232 | 1,30869347087323 | 2,75073186866818e-32 | NA |
| 151 | cds-pgaptmp_002234 | 1,03142338694938 | 1,61262058315189e-21 | NA |
| 152 | cds-pgaptmp_002240 | -1,62811366550197 | 1,15842119867126e-47 | NA |
| 153 | cds-pgaptmp_002258 | -1,19113910199433 | 1,23541986732955e-27 | NA |
| 154 | cds-pgaptmp_002279 | 3,09710668963456 | 2,46735449267589e-92 | NA |
| 155 | cds-pgaptmp_002281 | -1,18288326364116 | 9,6018958584576e-35 | NA |
| 156 | cds-pgaptmp_002283 | 2,57402895897818 | 6,12053444731967e-57 | NA |
| 157 | cds-pgaptmp_002301 | -2,20005944245098 | 3,42323564163051e-102 | NA |
| 158 | cds-pgaptmp_002304 | -1,20678292497663 | 1,34459733465306e-17 | NA |
| 159 | cds-pgaptmp_002305 | -1,10538350277381 | 5,08016335287233e-16 | NA |
| 160 | cds-pgaptmp_002306 | -1,25753365643859 | 2,17808901980097e-15 | gpM |
| 161 | cds-pgaptmp_002309 | -1,14770299660581 | 6,26450708487731e-05 | NA |
| 162 | cds-pgaptmp_002310 | -1,02643086170513 | 0,00204987946409474 | NA |
| 163 | cds-pgaptmp_002319 | -1,62222535844982 | 1,85689147319003e-30 | NA |
| 164 | cds-pgaptmp_002320 | -1,74303389415105 | 9,30442905842817e-24 | NA |
| 165 | cds-pgaptmp_002347 | 1,21313068260354 | 7,88289617392963e-28 | queA |
| 166 | cds-pgaptmp_002422 | -1,36130045440329 | 8,26090236821267e-38 | NA |
| 167 | cds-pgaptmp_002423 | -1,27320100216422 | 1,76623092734176e-39 | NA |
| 168 | cds-pgaptmp_002440 | 1,00412985198229 | 1,60267498966725e-06 | NA |
| 169 | exon-pgaptmp_002444-1 | 1,16329391103351 | 1,82588517450436e-14 | NA |
| 170 | exon-pgaptmp_002445-1 | 1,26261590986536 | 2,28311024341554e-05 | NA |
| 171 | exon-pgaptmp_002446-1 | 1,20905848849309 | 3,43512266310073e-06 | NA |
| 172 | cds-pgaptmp_002448 | -1,45214233331165 | 1,21974084476172e-33 | NA |
| 173 | cds-pgaptmp_002458 | 1,03212431939843 | 6,83400672034423e-20 | rplQ |
| 174 | cds-pgaptmp_002463 | 1,02873607735058 | 2,07099951794087e-20 | rpmJ |
| 175 | cds-pgaptmp_002464 | 1,05360903756068 | 3,62085335788965e-21 | secY |
| 176 | cds-pgaptmp_002465 | 1,05170144457496 | 1,34512641923886e-19 | rplO |
| 177 | cds-pgaptmp_002466 | 1,06994664273324 | 2,73298608225109e-20 | rpmD |
| 178 | cds-pgaptmp_002467 | 1,06459422586087 | 5,37541767785802e-21 | rpsE |
| 179 | cds-pgaptmp_002470 | 1,0363910549897 | 1,54341002316908e-18 | rpsH |
| 180 | cds-pgaptmp_002471 | 1,00720610189215 | 1,12784046627623e-18 | rpsN |
| 181 | cds-pgaptmp_002474 | 1,11358444220452 | 1,62812532525958e-27 | rplN |
| 182 | cds-pgaptmp_002476 | 1,01557103048807 | 1,41953544189211e-19 | rpmC |
| 183 | cds-pgaptmp_002513 | -1,17447255904395 | 8,84216635800101e-42 | hemF |
| 184 | cds-pgaptmp_002523 | 1,10007750466888 | 5,69973748981473e-33 | glyQ |
| 185 | cds-pgaptmp_002533 | -1,07260560968351 | 3,94771840726574e-05 | NA |
| 186 | cds-pgaptmp_002534 | 1,74910643199132 | 1,33177899045986e-100 | NA |
| 187 | cds-pgaptmp_002540 | 1,17287922419372 | 8,33837577753875e-31 | NA |
| 188 | cds-pgaptmp_002542 | 1,25512804647895 | 5,25427096115922e-30 | NA |
| 189 | cds-pgaptmp_002559 | -1,1647426884113 | 3,3697122516553e-30 | NA |
| 190 | cds-pgaptmp_002560 | -2,56882767760562 | 1,09911639012073e-137 | NA |
| 191 | cds-pgaptmp_002565 | 1,34125679768247 | 1,93755004760637e-33 | rplS |
| 192 | cds-pgaptmp_002566 | 1,1673874177537 | 1,49946312985757e-29 | trmD |
| 193 | cds-pgaptmp_002567 | 1,01082725889487 | 2,00342406399998e-23 | rimM |
| 194 | cds-pgaptmp_002568 | 1,13180904519972 | 4,78299929297283e-27 | rpsP |
| 195 | cds-pgaptmp_002597 | 1,01628169037891 | 1,09567073552774e-08 | NA |

| | | | | |
|---|---|---|---|---|
| 196 | cds-pgaptmp_002609 | 1,0100113488705 | 3,24086243958383e-09 | NA |
| 197 | cds-pgaptmp_002614 | 1,18640103867454 | 7,14963006598657e-30 | NA |
| 198 | cds-pgaptmp_002625 | 1,10014672958925 | 1,85349211252194e-14 | NA |
| 199 | cds-pgaptmp_002626 | -1,35031167857115 | 1,0026537208888e-37 | NA |
| 200 | cds-pgaptmp_002629 | 1,03047483857959 | 1,35747245095852e-20 | NA |
| 201 | cds-pgaptmp_002630 | 1,29543498933926 | 6,17932819800254e-38 | NA |
| 202 | cds-pgaptmp_002647 | 1,11945898327985 | 1,26400388038794e-21 | NA |
| 203 | cds-pgaptmp_002648 | 1,0654222040495 | 8,53236832410434e-29 | NA |
| 204 | cds-pgaptmp_002654 | 1,13160659197095 | 2,8702738657225e-39 | aqpZ |
| 205 | cds-pgaptmp_002658 | 1,75536697915263 | 5,57824492251277e-73 | NA |
| 206 | cds-pgaptmp_002694 | -3,12171476550325 | 1,07656259312527e-247 | NA |
| 207 | cds-pgaptmp_002704 | 1,21107859083892 | 1,60460685542773e-19 | NA |
| 208 | cds-pgaptmp_002706 | -1,07918992447324 | 1,10632661536212e-32 | ycaC |
| 209 | cds-pgaptmp_002708 | -1,22742626919318 | 2,42691385579427e-55 | NA |
| 210 | cds-pgaptmp_002709 | -1,10958902956075 | 4,16975991864909e-39 | gabT |
| 211 | cds-pgaptmp_002717 | 1,01485177559607 | 1,25739600945925e-25 | NA |
| 212 | cds-pgaptmp_002725 | 2,88075071051866 | 4,38438052465408e-185 | omp33-36 |
| 213 | cds-pgaptmp_002728 | -2,00775270181917 | 5,90774814970238e-56 | NA |
| 214 | cds-pgaptmp_002729 | -2,0465944032075 | 1,4096349715562e-65 | NA |
| 215 | cds-pgaptmp_002731 | -2,59635524911547 | 8,36651692632436e-115 | NA |
| 216 | cds-pgaptmp_002737 | -1,21729812031194 | 4,25434363502944e-40 | acs |
| 217 | cds-pgaptmp_002772 | -1,25617611349437 | 3,47493994060915e-54 | NA |
| 218 | cds-pgaptmp_002783 | 2,31163970473036 | 4,78483472373305e-93 | NA |
| 219 | cds-pgaptmp_002795 | 1,20244860852741 | 4,09963965204627e-43 | NA |
| 220 | cds-pgaptmp_002796 | 1,06157169100113 | 6,79663520446517e-25 | NA |
| 221 | cds-pgaptmp_002803 | 1,78850584304252 | 3,42426294618157e-36 | NA |
| 222 | cds-pgaptmp_002807 | -1,10556922021397 | 6,69884442900069e-29 | NA |
| 223 | cds-pgaptmp_002835 | 1,129838340914 | 1,73848150878666e-28 | hutH |
| 224 | cds-pgaptmp_002836 | 1,32595998409793 | 7,67211460440805e-27 | hutU |
| 225 | cds-pgaptmp_002837 | 1,35186024046743 | 1,2232520291278e-14 | NA |
| 226 | cds-pgaptmp_002839 | 1,13160167025064 | 2,15078538307564e-21 | NA |
| 227 | cds-pgaptmp_002843 | -1,24587304134192 | 2,63556402020271e-24 | fahA |
| 228 | cds-pgaptmp_002844 | -1,48819506177851 | 2,00223053399602e-26 | maiA |
| 229 | cds-pgaptmp_002845 | -1,4727951398324 | 4,33234124313143e-24 | NA |
| 230 | cds-pgaptmp_002856 | -1,02265035218194 | 6,13015481976453e-18 | NA |
| 231 | cds-pgaptmp_002858 | -2,37371325201286 | 1,00979742712125e-98 | NA |
| 232 | cds-pgaptmp_002860 | -2,58196911025253 | 8,15763508426324e-122 | NA |
| 233 | cds-pgaptmp_002878 | 1,24650926324419 | 6,22567641575396e-19 | NA |
| 234 | cds-pgaptmp_002879 | 1,27223910430614 | 0,000474834242334831 | NA |
| 235 | cds-pgaptmp_002891 | 1,56164088399836 | 4,55977203719839e-45 | NA |
| 236 | cds-pgaptmp_002907 | -1,16320906630876 | 1,0732284776523e-19 | NA |
| 237 | cds-pgaptmp_002935 | -2,44889504959043 | 1,32063443975171e-101 | NA |
| 238 | cds-pgaptmp_002936 | -2,93195131208697 | 1,03812087179857e-230 | NA |
| 239 | cds-pgaptmp_002946 | -2,15228328917519 | 5,33184115486986e-92 | NA |
| 240 | cds-pgaptmp_002948 | 1,81399266184171 | 1,77792405007571e-59 | NA |
| 241 | cds-pgaptmp_002954 | -1,0208966694014 | 2,04437217403532e-10 | NA |
| 242 | cds-pgaptmp_002969 | -5,66797555089216 | 0 | NA |
| 243 | cds-pgaptmp_002970 | -4,68366252510939 | 3,48990910787772e-198 | otsB |
| 244 | cds-pgaptmp_002973 | 1,00763903339166 | 5,27067876813571e-15 | NA |

| | | | | |
|---|---|---|---|---|
| 245 | cds-pgaptmp_002975 | 1,40498053120454 | 5,99320982907541e-33 | bioD |
| 246 | cds-pgaptmp_002983 | 1,27062617768162 | 1,25912256526357e-31 | fabD |
| 247 | cds-pgaptmp_002984 | 1,25203828179374 | 1,04837457419883e-27 | fabG |
| 248 | exon-pgaptmp_002996-1 | 1,36935029778086 | 2,37339183162919e-40 | NA |
| 249 | exon-pgaptmp_002997-1 | 1,97113228692213 | 6,45508801089918e-60 | NA |
| 250 | exon-pgaptmp_002998-1 | 2,31538448619733 | 4,13735794812586e-87 | NA |
| 251 | exon-pgaptmp_002999-1 | 1,81465888583629 | 3,84561377668387e-64 | NA |
| 252 | cds-pgaptmp_003000 | 1,46973467481364 | 1,0126721765946e-53 | ispE |
| 253 | cds-pgaptmp_003027 | 1,1016345916269 | 1,62647359589738e-22 | hemJ |
| 254 | cds-pgaptmp_003028 | 1,21907063507064 | 2,07555100355974e-25 | NA |
| 255 | cds-pgaptmp_003029 | -1,34983725735026 | 1,03548543306019e-46 | NA |
| 256 | cds-pgaptmp_003051 | -1,05181450912925 | 2,16856381969809e-32 | NA |
| 257 | cds-pgaptmp_003055 | 1,65339103568661 | 3,59713245724415e-24 | NA |
| 258 | cds-pgaptmp_003066 | -1,26781969651361 | 1,05731719350661e-21 | NA |
| 259 | cds-pgaptmp_003078 | 1,18006887037883 | 0,000700809158106293 | NA |
| 260 | exon-pgaptmp_003109-1 | -1,13010637496702 | 1,06074605431304e-14 | NA |
| 261 | cds-pgaptmp_003130 | -1,50499098144944 | 2,46741121067965e-29 | NA |
| 262 | cds-pgaptmp_003142 | 1,01344605608922 | 3,90619144723679e-23 | NA |
| 263 | cds-pgaptmp_003158 | 1,25592849804492 | 9,63908097735964e-36 | NA |
| 264 | cds-pgaptmp_003160 | -1,23945030705408 | 4,79226101663634e-40 | NA |
| 265 | cds-pgaptmp_003208 | -1,20689551175319 | 0,00115090423680532 | NA |
| 266 | cds-pgaptmp_003237 | 1,07826270728214 | 4,11088626987467e-30 | NA |
| 267 | cds-pgaptmp_003238 | 1,15204488425845 | 5,91961364042402e-36 | rpsU |
| 268 | exon-pgaptmp_003246-1 | 1,2162374877234 | 3,5411077705403e-22 | NA |
| 269 | exon-pgaptmp_003247-1 | 1,42596008124853 | 2,19118075726453e-11 | NA |
| 270 | cds-pgaptmp_003249 | -2,26821688334557 | 3,1225455363299e-61 | NA |
| 271 | cds-pgaptmp_003250 | -2,17463963933214 | 5,67771290932492e-54 | NA |
| 272 | cds-pgaptmp_003251 | -1,7173488035624 | 2,95494021697072e-14 | NA |
| 273 | cds-pgaptmp_003252 | -4,72439511959596 | 0 | NA |
| 274 | cds-pgaptmp_003253 | -2,61195716999726 | 3,89411873401935e-72 | NA |
| 275 | cds-pgaptmp_003254 | -3,21305166883151 | 5,31252876872947e-115 | NA |
| 276 | cds-pgaptmp_003255 | -3,24393643918514 | 8,42847208806929e-115 | NA |
| 277 | cds-pgaptmp_003256 | -3,12852841793416 | 1,20785007102046e-75 | NA |
| 278 | cds-pgaptmp_003257 | -3,49273112603398 | 5,3710352016611e-102 | NA |
| 279 | cds-pgaptmp_003263 | 1,68512627520182 | 5,34331098231968e-87 | gltS |
| 280 | cds-pgaptmp_003265 | 1,02284631904301 | 1,40921862067619e-16 | NA |
| 281 | cds-pgaptmp_003271 | 1,45670559743633 | 1,11531050391962e-05 | NA |
| 282 | cds-pgaptmp_003272 | 1,43619148357094 | 1,78086126175623e-06 | NA |
| 283 | cds-pgaptmp_003273 | 2,40395973352262 | 2,57813131216346e-146 | csuAB |
| 284 | cds-pgaptmp_003274 | 1,56924214427166 | 2,32597477181824e-40 | csuA |
| 285 | cds-pgaptmp_003275 | 2,06376065502491 | 1,62047289174746e-47 | csuB |
| 286 | cds-pgaptmp_003276 | 2,44653084289235 | 1,05941327602437e-123 | csuC |
| 287 | cds-pgaptmp_003277 | 2,37476981783726 | 2,62651324255098e-162 | csuD |
| 288 | cds-pgaptmp_003278 | 2,41791035299059 | 5,90779280444624e-119 | csuE |
| 289 | cds-pgaptmp_003294 | 2,53880458319593 | 5,52067460117795e-23 | NA |
| 290 | cds-pgaptmp_003295 | 2,27718861618521 | 4,2430360862779e-15 | NA |
| 291 | cds-pgaptmp_003296 | 2,05225663419395 | 1,48393095637854e-12 | NA |
| 292 | cds-pgaptmp_003297 | 1,29650987141463 | 7,87279759053493e-05 | NA |
| 293 | cds-pgaptmp_003298 | 1,08874741733467 | 1,1427373459248e-07 | NA |

| | | | | |
|---|---|---|---|---|
| 294 | cds-pgaptmp_003306 | -1,49663143176085 | 2,52000716085283e-27 | NA |
| 295 | cds-pgaptmp_003310 | 1,60401500765807 | 3,23277352705091e-25 | NA |
| 296 | cds-pgaptmp_003313 | -2,2385569765638 | 2,92255400734825e-105 | NA |
| 297 | cds-pgaptmp_003323 | -3,21935729311074 | 1,33895076152872e-205 | NA |
| 298 | cds-pgaptmp_003326 | -2,2726385868087 | 5,47921426328341e-65 | NA |
| 299 | cds-pgaptmp_003327 | -1,6778970314648 | 1,73298157949757e-33 | NA |
| 300 | cds-pgaptmp_003328 | -1,73180357241332 | 1,63485721897103e-55 | NA |
| 301 | cds-pgaptmp_003329 | -1,11400090280856 | 1,34508773963423e-29 | NA |
| 302 | cds-pgaptmp_003332 | 1,01627094513167 | 8,03758588229591e-17 | rplI |
| 303 | cds-pgaptmp_003333 | 1,02790792520056 | 7,13844690683291e-17 | rpsR |
| 304 | cds-pgaptmp_003334 | 1,14509015869183 | 8,95464032957175e-23 | rpsF |
| 305 | cds-pgaptmp_003350 | 1,0071026946142 | 2,45904157709541e-20 | NA |
| 306 | cds-pgaptmp_003355 | -1,0232060031161 | 2,97243366882542e-17 | NA |
| 307 | cds-pgaptmp_003356 | -1,87879392087762 | 5,92080799451287e-65 | NA |
| 308 | cds-pgaptmp_003388 | -1,61783712498724 | 1,18281053182239e-41 | NA |
| 309 | cds-pgaptmp_003389 | -1,0947627746672 | 4,2150949393178e-07 | NA |
| 310 | cds-pgaptmp_003391 | -1,24614730399686 | 0,000424066615840152 | NA |
| 311 | cds-pgaptmp_003422 | -2,16413060931444 | 4,27660692082281e-18 | clpB |
| 312 | cds-pgaptmp_003423 | -1,33043223460997 | 2,96299197200614e-21 | NA |
| 313 | cds-pgaptmp_003429 | -1,82119261621088 | 1,04629259855301e-69 | NA |
| 314 | cds-pgaptmp_003432 | 1,62154445789636 | 7,02766852927355e-42 | trmB |
| 315 | cds-pgaptmp_003435 | -1,45469948058397 | 2,10332843341912e-56 | NA |
| 316 | cds-pgaptmp_003439 | -2,29842141726394 | 4,87829151736797e-53 | NA |
| 317 | cds-pgaptmp_003453 | -1,17918085186211 | 5,98957841423538e-31 | NA |
| 318 | cds-pgaptmp_003456 | -4,35098341872298 | 3,06866428493619e-150 | NA |
| 319 | cds-pgaptmp_003457 | -3,75567963816089 | 5,80353892661993e-105 | NA |
| 320 | cds-pgaptmp_003460 | -1,35471002871068 | 9,22165663714277e-13 | xseB |
| 321 | cds-pgaptmp_003461 | -1,31597750548025 | 1,74520486217825e-32 | xseA |
| 322 | cds-pgaptmp_003472 | -2,88279707793826 | 4,74197943875286e-78 | NA |
| 323 | cds-pgaptmp_003473 | -2,23511023763953 | 2,03028494397426e-16 | NA |
| 324 | cds-pgaptmp_003474 | -2,85794069118363 | 9,02774550976592e-78 | NA |
| 325 | cds-pgaptmp_003475 | -1,17931730945406 | 1,47732937590746e-10 | NA |
| 326 | cds-pgaptmp_003479 | -4,00428441378747 | 1,54324574069251e-224 | NA |
| 327 | cds-pgaptmp_003484 | 1,26576891842859 | 3,4555072529102e-33 | NA |
| 328 | cds-pgaptmp_003485 | -1,01117226422831 | 3,14283399896481e-06 | NA |
| 329 | cds-pgaptmp_003486 | -3,40954777714216 | 1,16901098768055e-202 | NA |
| 330 | cds-pgaptmp_003487 | -1,42226470851596 | 9,2246367063718e-06 | NA |
| 331 | cds-pgaptmp_003490 | -1,58819444583502 | 1,86701251841309e-06 | terL |
| 332 | cds-pgaptmp_003493 | 1,76133727878674 | 1,20323220630443e-26 | NA |
| 333 | cds-pgaptmp_003494 | 1,3863810644645 | 2,31225979841478e-28 | NA |
| 334 | cds-pgaptmp_003500 | -1,33258195297855 | 4,82278080776693e-19 | NA |
| 335 | cds-pgaptmp_003519 | -1,99079042124374 | 8,73229589341169e-90 | NA |
| 336 | cds-pgaptmp_003527 | -1,3708395677889 | 1,12674551355541e-20 | NA |
| 337 | cds-pgaptmp_003534 | -1,2246689594548 | 4,11187641876691e-15 | NA |
| 338 | cds-pgaptmp_003536 | -1,76233931833432 | 1,14357332219189e-68 | NA |
| 339 | cds-pgaptmp_003540 | -1,96661049452487 | 1,49432654710789e-91 | NA |
| 340 | cds-pgaptmp_003545 | -1,4894320376881 | 1,98621854561192e-44 | NA |
| 341 | cds-pgaptmp_003550 | -1,09965227035355 | 1,10889966811583e-24 | NA |
| 342 | exon-pgaptmp_003561-1 | 1,22758430207287 | 1,64889830274727e-08 | NA |

| | | | | |
|---|---|---|---|---|
| 343 | cds-pgaptmp_003567 | -1,61655981830794 | 3,51917113736049e-43 | glpD |
| 344 | cds-pgaptmp_003581 | 1,0454409868934 | 1,50865425456842e-23 | ttcA |
| 345 | cds-pgaptmp_003583 | 1,03769323538915 | 6,74171158921827e-24 | NA |
| 346 | cds-pgaptmp_003598 | -1,76430585558639 | 5,19544448025986e-63 | NA |
| 347 | cds-pgaptmp_003599 | -1,64335751942641 | 6,73016299768196e-26 | fdhD |
| 348 | cds-pgaptmp_003635 | 1,0809444700281 | 1,52901498819925e-28 | purN |
| 349 | cds-pgaptmp_003645 | 1,13755368289096 | 1,36739796332184e-33 | NA |
| 350 | cds-pgaptmp_003658 | -1,62136987077212 | 9,68987537019662e-64 | NA |
| 351 | cds-pgaptmp_003659 | -1,5753716530016 | 3,95827668259614e-16 | NA |
| 352 | cds-pgaptmp_003660 | -1,06242336903842 | 8,27464642638521e-13 | NA |
| 353 | cds-pgaptmp_003661 | -1,60883141132191 | 6,79832521211022e-50 | NA |
| 354 | cds-pgaptmp_003663 | -1,02555170003317 | 3,37325349230727e-16 | NA |
| 355 | cds-pgaptmp_003670 | -1,34349022909387 | 3,29436705575501e-75 | carO |
| 356 | cds-pgaptmp_003675 | 1,11669449243714 | 2,09426815960299e-07 | NA |
| 357 | cds-pgaptmp_003676 | 1,96554807780226 | 7,77867770107739e-34 | NA |
| 358 | cds-pgaptmp_003702 | -1,1762491359511 | 1,0695607898801e-32 | NA |
| 359 | cds-pgaptmp_003703 | -1,8413889169632 | 5,51476416971974e-74 | uvrB |
| 360 | cds-pgaptmp_003704 | -1,72603919541891 | 8,01886991625391e-72 | NA |
| 361 | cds-pgaptmp_003716 | -1,33009583882406 | 7,39073408949028e-43 | NA |
| 362 | cds-pgaptmp_000334 | -1,07042819451192 | 1,65827082579526e-17 | NA |
| 363 | cds-pgaptmp_000336 | 1,54640083495693 | 2,51722174416404e-38 | rpsT |
| 364 | cds-pgaptmp_000337 | -1,52771659752005 | 1,30718146528994e-32 | NA |
| 365 | cds-pgaptmp_000361 | -1,61949646187093 | 2,30113237003562e-45 | NA |
| 366 | cds-pgaptmp_000375 | -1,37294129339694 | 1,58005121552955e-30 | mtlD |
| 367 | cds-pgaptmp_000383 | 1,98575163855031 | 2,10024855912377e-58 | NA |
| 368 | cds-pgaptmp_000396 | -1,79772578467751 | 7,43832945399019e-51 | NA |
| 369 | cds-pgaptmp_000398 | 1,45440869127579 | 5,42543273511251e-27 | NA |
| 370 | cds-pgaptmp_000402 | -1,66495576668773 | 5,32965525314267e-21 | NA |
| 371 | cds-pgaptmp_000406 | -1,02054305186457 | 4,93683029912084e-05 | NA |
| 372 | cds-pgaptmp_000408 | -1,88253230255394 | 1,14990557351456e-40 | NA |
| 373 | cds-pgaptmp_000414 | 1,05902969213413 | 3,84726807880231e-27 | prmB |
| 374 | cds-pgaptmp_000416 | -1,04649958889618 | 2,77533502732694e-25 | NA |
| 375 | cds-pgaptmp_000418 | -1,1247563824146 | 7,86941600686897e-13 | NA |
| 376 | cds-pgaptmp_000420 | -2,54864634988831 | 3,97881772512507e-18 | NA |
| 377 | cds-pgaptmp_000421 | -2,19390577941328 | 1,59049878006243e-22 | NA |
| 378 | cds-pgaptmp_000422 | -1,79355111457112 | 4,50415800656446e-19 | NA |
| 379 | cds-pgaptmp_000423 | -1,93272980267197 | 3,67396752472439e-17 | lpdA |
| 380 | cds-pgaptmp_000424 | -1,25601039851145 | 4,38974114505215e-07 | NA |
| 381 | cds-pgaptmp_000425 | -1,66407846749986 | 2,3095825895667e-36 | NA |
| 382 | cds-pgaptmp_000428 | -2,78420703222833 | 1,47136206243836e-126 | NA |
| 383 | cds-pgaptmp_000429 | -2,35846770971395 | 2,36559504567532e-123 | NA |
| 384 | cds-pgaptmp_000430 | -1,36112762975083 | 8,14997192862985e-16 | NA |
| 385 | cds-pgaptmp_000432 | -1,18420742859878 | 9,04339368905742e-21 | NA |
| 386 | cds-pgaptmp_000445 | -1,51108510665121 | 6,94339766532005e-19 | NA |
| 387 | cds-pgaptmp_000452 | -2,43006032818453 | 1,47335907415493e-140 | NA |
| 388 | cds-pgaptmp_000453 | -2,13820791475018 | 1,46338908789721e-44 | NA |
| 389 | cds-pgaptmp_000454 | -2,88728819912516 | 2,18400113956202e-107 | NA |
| 390 | cds-pgaptmp_000455 | -2,73491230108556 | 3,66130913416479e-81 | NA |
| 391 | cds-pgaptmp_000458 | -1,8999143856008 | 4,82868596754602e-49 | NA |

| | | | | |
|---|---|---|---|---|
| 392 | cds-pgaptmp_000459 | -2,17584468756925 | 4,23331447522102e-55 | NA |
| 393 | cds-pgaptmp_000460 | -1,19533542081103 | 8,84180425312095e-10 | NA |
| 394 | cds-pgaptmp_000472 | -1,615011674837 | 2,35698516727646e-43 | modA |
| 395 | cds-pgaptmp_000473 | -1,26197293629092 | 6,22228096380166e-17 | modB |
| 396 | cds-pgaptmp_000479 | -1,34726015257958 | 8,44200915543591e-32 | ppk1 |
| 397 | cds-pgaptmp_000486 | -2,34175953287256 | 7,37291279458677e-26 | NA |
| 398 | cds-pgaptmp_000489 | -1,67128322709893 | 3,10884392032926e-57 | NA |
| 399 | cds-pgaptmp_000490 | -3,42350619247791 | 1,27248235657552e-80 | NA |
| 400 | cds-pgaptmp_000491 | -1,38293045660654 | 5,11530506829582e-22 | NA |
| 401 | cds-pgaptmp_000494 | 1,74151915182981 | 4,466624089976e-46 | NA |
| 402 | cds-pgaptmp_000495 | 1,29569274560564 | 7,97542593334617e-36 | NA |
| 403 | cds-pgaptmp_000496 | 1,74564135911791 | 3,1055288619831e-67 | NA |
| 404 | cds-pgaptmp_000498 | 2,62287262175854 | 3,1828049169707e-114 | NA |
| 405 | cds-pgaptmp_000513 | 1,16164488787093 | 5,47464451891302e-33 | NA |
| 406 | cds-pgaptmp_000540 | -1,1221604916427 | 4,16905001098357e-34 | NA |
| 407 | cds-pgaptmp_000542 | -2,3186938321448 | 2,26937492591408e-53 | NA |
| 408 | cds-pgaptmp_000573 | -1,23823200090654 | 5,59422810301145e-43 | lon |
| 409 | cds-pgaptmp_000576 | 1,9475835606492 | 3,69414852017413e-109 | NA |
| 410 | cds-pgaptmp_000589 | -1,23005025426944 | 2,5794858829596e-17 | ureE |
| 411 | cds-pgaptmp_000594 | 1,16724275367224 | 2,53838679203392e-10 | NA |
| 412 | cds-pgaptmp_000595 | -1,02689721698731 | 1,57091700920392e-34 | NA |
| 413 | cds-pgaptmp_000596 | -1,11417655588879 | 1,85902900436382e-33 | NA |
| 414 | cds-pgaptmp_000603 | 1,08562267452674 | 9,5272184783594e-35 | cysD |
| 415 | cds-pgaptmp_000607 | -1,37962531598247 | 6,76910716003145e-17 | NA |
| 416 | cds-pgaptmp_000621 | 1,12298160759341 | 3,75031077842965e-18 | NA |
| 417 | cds-pgaptmp_000629 | -1,92664697118373 | 7,01598162766331e-36 | NA |
| 418 | cds-pgaptmp_000632 | 1,20269513828874 | 5,24061043722423e-29 | NA |
| 419 | cds-pgaptmp_000651 | -1,11133025617628 | 2,62311795362432e-05 | NA |
| 420 | cds-pgaptmp_000653 | -1,16253070435716 | 4,49043700239377e-07 | NA |
| 421 | cds-pgaptmp_000659 | 1,94525556580318 | 4,67598002993578e-05 | NA |
| 422 | cds-pgaptmp_000668 | 1,31967309222996 | 4,31406058229354e-19 | NA |
| 423 | cds-pgaptmp_000675 | 1,44342962611788 | 2,44075343046343e-49 | NA |
| 424 | cds-pgaptmp_000676 | 2,98382783498351 | 5,04213966914015e-133 | betI |
| 425 | cds-pgaptmp_000677 | 3,01936010094156 | 1,23316873002777e-133 | betB |
| 426 | cds-pgaptmp_000678 | 3,35947176499493 | 5,07987790169454e-181 | betA |
| 427 | cds-pgaptmp_000679 | 1,18452519930262 | 6,0432789915557e-26 | mqo |
| 428 | cds-pgaptmp_000681 | 2,46771002000967 | 6,6970540909918e-50 | NA |
| 429 | cds-pgaptmp_000682 | 1,67391967027674 | 1,69171934019612e-33 | NA |
| 430 | cds-pgaptmp_000685 | -1,22968243772835 | 3,50274217281888e-20 | NA |
| 431 | cds-pgaptmp_000693 | 1,92513669234491 | 0,000732349760290496 | NA |
| 432 | cds-pgaptmp_000701 | -3,78515817821282 | 8,82016251188775e-132 | NA |
| 433 | cds-pgaptmp_000702 | -1,2630680463827 | 1,94072143310889e-06 | NA |
| 434 | cds-pgaptmp_000707 | -2,85685906892175 | 4,4647770469058e-13 | NA |
| 435 | cds-pgaptmp_000708 | -1,80364815557085 | 1,46251076342166e-13 | NA |
| 436 | cds-pgaptmp_000710 | -2,07556501002143 | 1,05660823203852e-90 | NA |
| 437 | cds-pgaptmp_000721 | 1,05205611100692 | 5,07999575062717e-05 | aceI |
| 438 | cds-pgaptmp_000730 | -1,02042521877629 | 2,99592365784239e-12 | NA |
| 439 | cds-pgaptmp_000731 | -1,15036502135722 | 2,65968597785786e-27 | NA |
| 440 | cds-pgaptmp_000740 | -1,18368595617138 | 1,21499839080918e-21 | NA |

| | | | | |
|---|---|---|---|---|
| 441 | cds-pgaptmp_000744 | 1,02329832769576 | 4,51395374514946e-18 | accB |
| 442 | cds-pgaptmp_000756 | -1,45723123530972 | 9,76420453471924e-33 | moaA |
| 443 | cds-pgaptmp_000757 | -1,65971730322293 | 6,08972598559695e-23 | NA |
| 444 | cds-pgaptmp_000758 | -1,1769602806311 | 1,28209943742445e-25 | NA |
| 445 | cds-pgaptmp_000768 | -1,52621906618842 | 5,19097211435297e-57 | NA |
| 446 | cds-pgaptmp_000769 | -1,50159052810852 | 4,14631248659881e-64 | fumC |
| 447 | cds-pgaptmp_000770 | -1,25677441909244 | 1,30980199228796e-17 | NA |
| 448 | cds-pgaptmp_000773 | -2,82066080943584 | 3,36591488007515e-243 | NA |
| 449 | cds-pgaptmp_000797 | -1,03737751604312 | 4,55865569624823e-23 | NA |
| 450 | cds-pgaptmp_000799 | -1,18236619857777 | 7,4785988123489e-11 | NA |
| 451 | cds-pgaptmp_000800 | -1,01013334098561 | 1,6475918686987e-08 | NA |
| 452 | cds-pgaptmp_000808 | -1,12941781891297 | 6,13531015046657e-24 | NA |
| 453 | cds-pgaptmp_000810 | -2,63987481363184 | 3,89606673422398e-67 | NA |
| 454 | cds-pgaptmp_000812 | -2,50618504475222 | 7,85556566548005e-94 | NA |
| 455 | cds-pgaptmp_000816 | 2,10608485526205 | 2,82281067770786e-51 | NA |
| 456 | cds-pgaptmp_000824 | 1,20052991377678 | 6,31162865397537e-31 | brnQ |
| 457 | cds-pgaptmp_000830 | -1,71719103614165 | 3,79538721711167e-74 | NA |
| 458 | cds-pgaptmp_000833 | -2,40969168970274 | 4,73159896853501e-94 | NA |
| 459 | cds-pgaptmp_000834 | -1,15675969002484 | 9,30847334927935e-23 | NA |
| 460 | cds-pgaptmp_000838 | -1,00123777342358 | 6,55760571023055e-33 | NA |
| 461 | cds-pgaptmp_000839 | -1,00505155803705 | 1,24686059300386e-31 | cydX |
| 462 | cds-pgaptmp_000875 | -2,0379547916201 | 1,47053746713332e-24 | NA |
| 463 | cds-pgaptmp_000876 | -2,02306889535554 | 1,21671586657618e-26 | NA |
| 464 | cds-pgaptmp_000877 | -2,02779544527229 | 3,47050732427416e-33 | pcaF |
| 465 | cds-pgaptmp_000878 | -1,80339827931363 | 1,45507022081875e-24 | NA |
| 466 | cds-pgaptmp_000879 | -1,67858034480715 | 9,58533403989884e-13 | pcaD |
| 467 | cds-pgaptmp_000880 | -1,81100024919487 | 1,265503723766e-20 | NA |
| 468 | cds-pgaptmp_000881 | -1,6358900261688 | 3,26695125302518e-13 | pcaC |
| 469 | cds-pgaptmp_000882 | -1,63088267057943 | 4,58410795100275e-17 | pcaH |
| 470 | cds-pgaptmp_000883 | -1,51596006949274 | 4,33831252666222e-15 | pcaG |
| 471 | cds-pgaptmp_000884 | -1,24182479000045 | 3,63588800608414e-05 | aroD |
| 472 | cds-pgaptmp_000906 | -1,02692999055323 | 0,00952720208052033 | NA |
| 473 | cds-pgaptmp_000909 | -1,06096932888667 | 0,000529904537449128 | NA |
| 474 | cds-pgaptmp_000915 | -1,11456398087627 | 1,60225121828608e-07 | pcaD |
| 475 | cds-pgaptmp_000916 | -1,05104907497404 | 6,08926836720105e-07 | pcaF |
| 476 | cds-pgaptmp_000917 | -1,06232510229102 | 1,18363202744154e-05 | NA |
| 477 | cds-pgaptmp_000919 | -1,08688945770885 | 1,22055308536198e-11 | catA |
| 478 | cds-pgaptmp_000922 | -1,13586957532542 | 1,09465400688617e-20 | NA |
| 479 | cds-pgaptmp_000930 | -2,28639470958796 | 9,94852932463566e-161 | NA |
| 480 | cds-pgaptmp_000932 | -1,34735210018609 | 5,65071245482444e-37 | NA |
| 481 | cds-pgaptmp_000935 | -3,8576912549264 | 2,99973133621223e-156 | NA |
| 482 | cds-pgaptmp_000947 | -1,34494411251996 | 7,53791588741914e-21 | NA |
| 483 | cds-pgaptmp_000948 | -1,6703567329004 | 4,44266857837272e-61 | NA |
| 484 | cds-pgaptmp_000950 | -1,08094274190649 | 1,36618238591329e-32 | NA |
| 485 | cds-pgaptmp_000961 | 1,21685062100048 | 9,4395027791102e-16 | NA |
| 486 | cds-pgaptmp_000962 | 1,20218638670114 | 3,96521976708036e-25 | NA |
| 487 | cds-pgaptmp_000971 | 1,3636433582463 | 9,41115894029357e-68 | tviB |
| 488 | cds-pgaptmp_000975 | 1,12007531531799 | 1,91525254117419e-26 | NA |
| 489 | cds-pgaptmp_000976 | 1,53406973458774 | 7,82588456113676e-63 | NA |

| | | | | |
|---|---|---|---|---|
| 490 | cds-pgaptmp_000977 | 1,18626967854583 | 3,4224449743054e-56 | NA |
| 491 | cds-pgaptmp_000978 | 1,15751687684177 | 1,29245987374695e-42 | NA |
| 492 | cds-pgaptmp_000979 | 1,15402725277715 | 1,12117293312106e-34 | NA |
| 493 | cds-pgaptmp_000980 | 1,12561663327999 | 5,17531191415715e-47 | NA |
| 494 | cds-pgaptmp_000987 | 5,48203518890058 | 0 | lldP |
| 495 | cds-pgaptmp_000988 | 3,11876601074598 | 2,40914976669193e-177 | lldR |
| 496 | cds-pgaptmp_000989 | 3,24949119652472 | 9,34332130693336e-245 | lldD |
| 497 | cds-pgaptmp_000990 | 2,61486715626549 | 1,31964814246308e-131 | dld |
| 498 | cds-pgaptmp_000994 | -1,20628784538915 | 7,29942708160528e-44 | prpC |
| 499 | cds-pgaptmp_000995 | -1,15766003474982 | 1,13354753403966e-38 | acnD |
| 500 | cds-pgaptmp_001013 | -1,48144169522712 | 3,72091350081318e-35 | NA |
| 501 | cds-pgaptmp_001014 | -1,38227677374083 | 4,21322141806656e-06 | alr |
| 502 | cds-pgaptmp_001015 | -1,18906395867013 | 0,000683836341571858 | NA |
| 503 | cds-pgaptmp_001016 | -1,07169153040381 | 0,000734417205605664 | NA |
| 504 | cds-pgaptmp_001017 | 1,60777070596387 | 1,5162971192427e-46 | NA |
| 505 | cds-pgaptmp_001019 | -1,79933764976391 | 4,73276416140312e-23 | NA |
| 506 | cds-pgaptmp_001020 | -1,81047249487277 | 8,39655104671062e-14 | mmsB |
| 507 | exon-pgaptmp_001037-1 | 1,60315466274106 | 4,38722980045122e-05 | NA |
| 508 | exon-pgaptmp_001038-1 | 1,77567440149007 | 1,15492433036758e-06 | NA |
| 509 | cds-pgaptmp_001063 | 1,18775201341488 | 4,31316929373905e-33 | NA |
| 510 | cds-pgaptmp_001075 | 1,48154079573573 | 2,17410247734589e-34 | NA |
| 511 | cds-pgaptmp_001082 | 1,65820838496422 | 1,12672405770576e-49 | NA |
| 512 | cds-pgaptmp_001083 | 3,18547059905481 | 4,20594809391279e-159 | NA |
| 513 | cds-pgaptmp_001084 | 2,97582647444654 | 3,86990327530403e-125 | NA |
| 514 | cds-pgaptmp_001087 | 1,81518891371499 | 3,37989703321065e-48 | pgaA |
| 515 | cds-pgaptmp_001088 | 1,78598425215136 | 4,7353977003131e-31 | pgaB |
| 516 | cds-pgaptmp_001089 | 1,75899784262844 | 1,02988364828521e-25 | pgaC |
| 517 | cds-pgaptmp_001090 | 1,08767350035704 | 5,54683167340101e-07 | pgaD |
| 518 | cds-pgaptmp_001093 | 3,4845326391018 | 6,07897523438544e-230 | NA |
| 519 | cds-pgaptmp_001094 | 2,45084456249237 | 1,72021215843987e-147 | NA |
| 520 | cds-pgaptmp_001102 | -3,33207270897967 | 4,75014387429847e-124 | NA |
| 521 | cds-pgaptmp_001103 | -2,87042969373446 | 8,36713334458037e-59 | NA |
| 522 | cds-pgaptmp_001104 | -2,40694568390002 | 1,03420879070382e-52 | NA |
| 523 | cds-pgaptmp_001110 | 1,90737693615008 | 1,20026806415597e-39 | kdpA |
| 524 | cds-pgaptmp_001111 | 1,59019934706202 | 1,69134845345263e-39 | kdpB |
| 525 | cds-pgaptmp_001112 | 1,4635664482418 | 1,30999188250733e-31 | kdpC |
| 526 | cds-pgaptmp_001125 | 1,09322545546986 | 1,85661894022762e-19 | NA |
| 527 | cds-pgaptmp_001126 | 1,76960470738047 | 4,01757232746769e-17 | NA |
| 528 | cds-pgaptmp_001130 | -1,86997856579202 | 2,92729102288047e-44 | NA |
| 529 | cds-pgaptmp_001132 | -1,44763528319578 | 2,0796399525068e-37 | NA |
| 530 | cds-pgaptmp_001152 | 1,30264913013076 | 5,18464902069525e-21 | NA |
| 531 | cds-pgaptmp_001157 | -2,39823383849506 | 7,67201979722487e-33 | NA |
| 532 | cds-pgaptmp_001159 | -3,16362132174127 | 3,28174376106342e-119 | NA |
| 533 | cds-pgaptmp_001160 | -1,47182969246769 | 5,81024275683798e-65 | fadB |
| 534 | cds-pgaptmp_001161 | -1,41329190668887 | 3,33845554093105e-56 | fadA |
| 535 | cds-pgaptmp_001163 | -2,88368064023585 | 5,35745020214201e-111 | NA |
| 536 | cds-pgaptmp_001172 | 1,98645108464615 | 1,63949354393421e-81 | NA |
| 537 | cds-pgaptmp_001176 | 1,11108328215998 | 2,85543555039025e-08 | NA |
| 538 | cds-pgaptmp_001179 | 1,02904664679384 | 4,2128135697485e-23 | rplL |

| | | | | |
|---|---|---|---|---|
| 539 | cds-pgaptmp_001180 | 1,00486481148203 | 5,53091839272684e-22 | rplJ |
| 540 | exon-pgaptmp_001185-1 | 1,11526199949378 | 2,23534526370313e-20 | NA |
| 541 | exon-pgaptmp_001188-1 | 1,03961601738268 | 2,43717941994079e-31 | NA |
| 542 | exon-pgaptmp_001189-1 | 1,03233231241303 | 1,66798608988832e-23 | NA |
| 543 | cds-pgaptmp_001205 | 1,15685468289059 | 1,66345331270123e-34 | argH |
| 544 | cds-pgaptmp_001206 | 1,30732921932089 | 1,75891607835802e-35 | NA |
| 545 | cds-pgaptmp_001222 | 1,08321197035227 | 1,26604986221173e-18 | xerD |
| 546 | cds-pgaptmp_001223 | 1,17934839298611 | 3,31129365364501e-33 | NA |
| 547 | cds-pgaptmp_001237 | -1,25530999966241 | 1,29850366030343e-20 | NA |
| 548 | cds-pgaptmp_001271 | 1,0139750985288 | 1,92103014804642e-08 | arsC |
| 549 | cds-pgaptmp_001275 | 1,09492913366178 | 1,5262689758973e-15 | lspA |
| 550 | cds-pgaptmp_001276 | 1,044037814855 | 2,22150998060177e-16 | NA |
| 551 | cds-pgaptmp_001299 | -1,02442624871143 | 8,90271727291459e-07 | NA |
| 552 | cds-pgaptmp_001304 | 1,22697904121686 | 1,2322015756295e-35 | adeB |
| 553 | cds-pgaptmp_001305 | 1,6735597926901 | 1,21371161756414e-43 | adeA |
| 554 | cds-pgaptmp_001308 | -1,18780364720008 | 1,1643233715334e-09 | NA |
| 555 | cds-pgaptmp_001332 | -2,06914505459969 | 3,22496399396554e-39 | iaaH |
| 556 | cds-pgaptmp_001333 | -1,07180228979349 | 2,48604326333196e-05 | NA |
| 557 | cds-pgaptmp_001334 | 1,75244069973021 | 0,000639330056586274 | NA |
| 558 | cds-pgaptmp_001349 | 2,50515576281669 | 1,15297720390841e-92 | NA |
| 559 | cds-pgaptmp_001350 | 2,96423410140645 | 1,38444759790844e-153 | NA |
| 560 | cds-pgaptmp_001354 | 1,2142747432073 | 2,83590557291373e-30 | NA |
| 561 | cds-pgaptmp_001355 | 1,12648471942767 | 7,57053187425864e-20 | NA |
| 562 | cds-pgaptmp_001370 | -1,4129582665109 | 1,33424272050404e-27 | NA |
| 563 | cds-pgaptmp_001373 | -1,10916782120619 | 2,52545360603934e-21 | add |
| 564 | exon-pgaptmp_001374-1 | 1,57004441768485 | 1,45617849624864e-40 | NA |
| 565 | exon-pgaptmp_001375-1 | 1,70691655503115 | 3,0059313566954e-51 | NA |
| 566 | exon-pgaptmp_001376-1 | 1,73312172314536 | 9,96248407777516e-62 | NA |
| 567 | exon-pgaptmp_001377-1 | 1,54776836711934 | 3,51201133635779e-41 | NA |
| 568 | cds-pgaptmp_001393 | -1,00855680499067 | 7,68510016756793e-18 | NA |
| 569 | cds-pgaptmp_001408 | -2,8097206972239 | 1,63263847446443e-161 | raiA |
| 570 | cds-pgaptmp_001703 | -1,471589746875 | 3,19585964144239e-14 | NA |
| 571 | cds-pgaptmp_001706 | -1,12455292250079 | 2,68251028198043e-16 | NA |
| 572 | cds-pgaptmp_001716 | -1,44749820967739 | 1,28088128370085e-41 | NA |
| 573 | cds-pgaptmp_001733 | -1,43687518776262 | 6,20130558138056e-47 | NA |
| 574 | cds-pgaptmp_001765 | 1,02720536207844 | 0,00120319508294217 | NA |
| 575 | cds-pgaptmp_001770 | -3,7253268439093 | 8,56010448532981e-176 | NA |
| 576 | cds-pgaptmp_001773 | 1,22535590743686 | 8,65945877157017e-20 | NA |
| 577 | cds-pgaptmp_001774 | 1,2654341858219 | 1,66237933092241e-22 | NA |
| 578 | cds-pgaptmp_001778 | -3,22068602353926 | 7,70838653228015e-236 | NA |
| 579 | cds-pgaptmp_001779 | -3,18780147723609 | 1,31529352799309e-225 | NA |
| 580 | cds-pgaptmp_001780 | -3,11442639355351 | 9,43930040401441e-234 | NA |
| 581 | cds-pgaptmp_001781 | -3,24675344962926 | 0 | pxpA |
| 582 | cds-pgaptmp_001782 | -3,23598917405446 | 3,97892327412244e-289 | NA |
| 583 | cds-pgaptmp_001783 | -1,08251846885475 | 7,61090119075799e-14 | mumR |
| 584 | cds-pgaptmp_001833 | 1,04865710203246 | 5,2346536505377e-19 | NA |
| 585 | cds-pgaptmp_001842 | 1,5485025386746 | 1,28269899748778e-47 | NA |
| 586 | cds-pgaptmp_001848 | 1,90060928787884 | 1,46573853699643e-24 | NA |
| 587 | cds-pgaptmp_001856 | 1,53464720092619 | 2,09535881186905e-27 | NA |

| | | | | |
|---|---|---|---|---|
| 588 | cds-pgaptmp_001857 | 1,23505764173216 | 1,88951466653275e-12 | NA |
| 589 | cds-pgaptmp_001858 | 1,00796077325909 | 1,02999905949041e-09 | NA |
| 590 | cds-pgaptmp_001859 | 1,38981569475741 | 8,6147052868748e-20 | NA |
| 591 | cds-pgaptmp_001863 | -1,74087030406521 | 9,21629822521607e-59 | NA |
| 592 | cds-pgaptmp_001869 | 1,17336474101866 | 4,60950277576878e-34 | yidD |
| 593 | cds-pgaptmp_001880 | -1,39850817388699 | 2,71839638390787e-38 | NA |
| 594 | cds-pgaptmp_001885 | 1,1148951821807 | 1,47831885584635e-27 | tyrS |
| 595 | cds-pgaptmp_001890 | -4,53099559005275 | 0 | NA |
| 596 | cds-pgaptmp_001891 | -1,61645235187651 | 2,67585126212342e-18 | NA |
| 597 | cds-pgaptmp_001892 | -5,68232593556068 | 1,35723875098789e-284 | NA |
| 598 | cds-pgaptmp_001893 | -3,26776508320989 | 1,8383897553934e-80 | NA |
| 599 | cds-pgaptmp_001894 | 1,74892443150968 | 5,02505084282998e-22 | NA |
| 600 | cds-pgaptmp_001895 | 1,04675308421257 | 1,81397493937112e-14 | pstC |
| 601 | cds-pgaptmp_001909 | 1,40253776247352 | 2,21597500996541e-39 | surE |
| 602 | cds-pgaptmp_001916 | 1,0709732982316 | 2,41820001722706e-20 | rpmE |
| 603 | cds-pgaptmp_001920 | 1,14340054288498 | 1,43650432458727e-31 | efp |
| 604 | cds-pgaptmp_001925 | -1,04455404269452 | 1,45630763186468e-32 | NA |
| 605 | cds-pgaptmp_001932 | -1,0600536940074 | 1,33783120562044e-16 | NA |
| 606 | cds-pgaptmp_001935 | -1,31670212528716 | 3,42769825409549e-15 | NA |
| 607 | cds-pgaptmp_001947 | -1,47343366896085 | 7,21505977194199e-68 | icd |
| 608 | cds-pgaptmp_001955 | -1,56166588562309 | 9,26955231193881e-33 | NA |
| 609 | cds-pgaptmp_001956 | -1,84978428809269 | 4,28024611434102e-39 | NA |
| 610 | cds-pgaptmp_001957 | -1,87494959304737 | 1,12081286784755e-62 | NA |
| 611 | cds-pgaptmp_001958 | -2,34006915590338 | 2,09484391795419e-125 | NA |
| 612 | cds-pgaptmp_001959 | -1,00932546901328 | 7,57750876152935e-22 | NA |
| 613 | cds-pgaptmp_002005 | 2,3469339084202 | 1,93955483506887e-50 | NA |
| 614 | cds-pgaptmp_002018 | 1,38819757466857 | 1,0804298966421e-11 | NA |
| 615 | cds-pgaptmp_002051 | 1,09706849010114 | 4,82247890918336e-18 | NA |
| 616 | cds-pgaptmp_002058 | -1,28090645502799 | 1,33040964484837e-08 | NA |
| 617 | cds-pgaptmp_002059 | -1,33077394464336 | 7,62609429984606e-14 | NA |
| 618 | cds-pgaptmp_002071 | 1,42859839179094 | 2,23550404443606e-39 | rpsI |
| 619 | cds-pgaptmp_002072 | 1,30441413004965 | 3,09505560248471e-34 | rplM |
| 620 | cds-pgaptmp_002351 | 1,54639440526571 | 1,8186790619223e-39 | tgt |
| 621 | cds-pgaptmp_002352 | 1,23049604029275 | 4,8751037250183e-40 | yajC |
| 622 | cds-pgaptmp_002354 | 1,0165236637091 | 5,29012203781355e-25 | secF |
| 623 | exon-pgaptmp_002399-1 | 1,69401086647247 | 7,96704824825883e-42 | NA |
| 624 | exon-pgaptmp_002400-1 | 1,78794818513243 | 1,9916207740371e-47 | NA |
| 625 | exon-pgaptmp_002402-1 | 1,32149866061869 | 1,14057509716575e-05 | NA |
| 626 | cds-pgaptmp_002405 | -1,15822868321449 | 7,79798038012364e-19 | xdhB |

protein
hypothetical protein
phosphoglycerate kinase
NAD-dependent deacetylase
sodium/proline symporter PutP
FxsA family protein
DMT family transporter
hypothetical protein
1-acyl-sn-glycerol-3-phosphate acyltransferase
YebC/PmpR family DNA-binding transcriptional regulator
amino acid ABC transporter ATP-binding protein
amino acid ABC transporter permease
amino acid ABC transporter permease
amino acid ABC transporter substrate-binding protein
redox-regulated ATPase YchF
dicarboxylate/amino acid:cation symporter
amidohydrolase
taurine ABC transporter substrate-binding protein
hypothetical protein
cytochrome ubiquinol oxidase subunit I
cytochrome d ubiquinol oxidase subunit II
DUF2474 domain-containing protein
malonate decarboxylase subunit delta
triphosphoribosyl-dephospho-CoA synthase
malonate decarboxylase subunit alpha
CidA/LrgA family protein
glutathione transferase
glutathione-dependent disulfide-bond oxidoreductase
amino acid ABC transporter ATP-binding protein
amino acid ABC transporter permease
amino acid ABC transporter permease
LysE family translocator
KGG domain-containing protein
DUF6367 family protein
SDR family oxidoreductase
catalase HPII
iron-containing redox enzyme family protein
CinA family protein
hypothetical protein
surface antigen protein 1
hypothetical protein
hypothetical protein
hypothetical protein
DcaP family trimeric outer membrane transporter
class I SAM-dependent methyltransferase
AMP-binding protein
TetR/AcrR family transcriptional regulator
isovaleryl-CoA dehydrogenase
carboxyl transferase domain-containing protein

enoyl-CoA hydratase/isomerase family protein
acetyl-CoA carboxylase biotin carboxylase subunit
hydroxymethylglutaryl-CoA lyase
indolepyruvate ferredoxin oxidoreductase family protein
cytochrome b/b6 domain-containing protein
catalase family peroxidase
hypothetical protein
hypothetical protein
hypothetical protein
cold-shock protein
hypothetical protein
hypothetical protein
hypothetical protein
hypothetical protein
PaaI family thioesterase
DapH/DapD/GlmU-related protein
phenylacetic acid degradation operon negative regulatory protein PaaX
phenylacetate--CoA ligase PaaK
3-oxoadipyl-CoA thiolase
3-hydroxyacyl-CoA dehydrogenase
2-(1,2-epoxy-1,2-dihydrophenyl)acetyl-CoA isomerase PaaG
enoyl-CoA hydratase-related protein
1,2-phenylacetyl-CoA epoxidase subunit PaaE
1,2-phenylacetyl-CoA epoxidase subunit PaaD
1,2-phenylacetyl-CoA epoxidase subunit PaaC
1,2-phenylacetyl-CoA epoxidase subunit PaaB
1,2-phenylacetyl-CoA epoxidase subunit PaaA
phenylacetic acid degradation bifunctional protein PaaZ
FAD-dependent oxidoreductase
hypothetical protein
lipid A hydroxylase LpxO
NA
ribosome-binding factor A
AAA family ATPase
chaperone modulator CbpM
DnaJ C-terminal domain-containing protein
SAM-dependent methyltransferase
NA
long-chain-acyl-CoA synthetase
glutathione S-transferase family protein
hypothetical protein
50S ribosomal protein L33
50S ribosomal protein L28
coniferyl aldehyde dehydrogenase
2-isopropylmalate synthase
TonB-dependent receptor
trigger factor
phosphate acetyltransferase
acetate/propionate family kinase

phosphogluconate dehydratase
bifunctional 4-hydroxy-2-oxoglutarate aldolase/2-dehydro-3-deoxy-phosphogluconate aldolase
gluconate:H symporter
gluconokinase
alpha/beta hydrolase
phosphatase PAP2 family protein
enoyl-ACP reductase
MacB family efflux pump subunit
MacA family efflux pump subunit
aconitate hydratase AcnA
hypothetical protein
heavy metal translocating P-type ATPase
hypothetical protein
multicopper oxidase domain-containing protein
copper resistance protein B
Re/Si-specific NAD(P)() transhydrogenase subunit alpha
proton-translocating transhydrogenase family protein
NAD(P)() transhydrogenase (Re/Si-specific) subunit beta
crotonase/enoyl-CoA hydratase family protein
acyl-CoA synthetase
Ig-like domain-containing protein
succinate dehydrogenase, hydrophobic membrane anchor protein
rhodanese-related sulfurtransferase
hypothetical protein
iron-containing alcohol dehydrogenase
50S ribosomal protein L27
50S ribosomal protein L21
phosphatase PAP2 family protein
multidrug efflux RND transporter periplasmic adaptor subunit AdeI
multidrug efflux RND transporter permease subunit AdeJ
multidrug efflux RND transporter outer membrane channel subunit AdeK
SDR family NAD(P)-dependent oxidoreductase
PA1571 family protein
DJ-1/PfpI family protein
alanine/glycine:cation symporter family protein
hypothetical protein
hypothetical protein
PepSY domain-containing protein
YdcF family protein
NA
NA
NA
NA
NA
NA
entericidin A/B family lipoprotein
YegP family protein
zinc ribbon domain-containing protein YjdM
SRPBCC family protein

DsbC family protein
large conductance mechanosensitive channel protein MscL
translational GTPase TypA
uracil-xanthine permease family protein
amino acid permease
VOC family protein
peroxiredoxin
hypothetical protein
acyl-CoA dehydrogenase C-terminal domain-containing protein
RcnB family protein
HAMP domain-containing sensor histidine kinase
GPO family capsid scaffolding protein
phage major capsid protein, P2 family
phage terminase small subunit
putative holin
phage holin family protein
phage tail sheath protein
phage major tail tube protein
tRNA preQ1(34) S-adenosylmethionine ribosyltransferase-isomerase QueA
hypothetical protein
DUF2789 family protein
nitroreductase family protein
NA
NA
NA
M3 family metallopeptidase
50S ribosomal protein L17
50S ribosomal protein L36
preprotein translocase subunit SecY
50S ribosomal protein L15
50S ribosomal protein L30
30S ribosomal protein S5
30S ribosomal protein S8
30S ribosomal protein S14
50S ribosomal protein L14
50S ribosomal protein L29
oxygen-dependent coproporphyrinogen oxidase
glycine--tRNA ligase subunit alpha
hypothetical protein
hypothetical protein
Glu/Leu/Phe/Val dehydrogenase
hypothetical protein
EamA family transporter
hemerythrin domain-containing protein
50S ribosomal protein L19
tRNA (guanosine(37)-N1)-methyltransferase TrmD
ribosome maturation factor RimM
30S ribosomal protein S16
type 4a pilus biogenesis protein PilO

sulfate ABC transporter substrate-binding protein
amino acid permease
hypothetical protein
tetratricopeptide repeat protein
SDR family NAD(P)-dependent oxidoreductase
SulP family inorganic anion transporter
antibiotic biosynthesis monooxygenase
DUF4870 family protein
aquaporin Z
DNA/RNA non-specific endonuclease
MFS transporter
FAD-dependent oxidoreductase
isochorismate family cysteine hydrolase YcaC
NAD-dependent succinate-semialdehyde dehydrogenase
4-aminobutyrate--2-oxoglutarate transaminase
DUF475 domain-containing protein
porin Omp33-36
cation acetate symporter
DUF485 domain-containing protein
hypothetical protein
acetate--CoA ligase
DUF2147 domain-containing protein
outer membrane protein OmpK
type VI secretion system Vgr family protein
disulfide bond formation protein B
SDR family NAD(P)-dependent oxidoreductase
alpha/beta fold hydrolase
histidine ammonia-lyase
urocanate hydratase
HutD family protein
acyltransferase family protein
fumarylacetoacetase
maleylacetoacetate isomerase
VOC family protein
formate/nitrite transporter family protein
PQQ-dependent sugar dehydrogenase
DUF1328 domain-containing protein
efflux RND transporter permease subunit
efflux RND transporter periplasmic adaptor subunit
TIGR04219 family outer membrane beta-barrel protein
beta strand repeat-containing protein
hypothetical protein
mechanosensitive ion channel
hypothetical protein
5'-methylthioadenosine/S-adenosylhomocysteine nucleosidase
DUF6160 family protein
trehalose-6-phosphate synthase
trehalose-phosphatase
8-amino-7-oxononanoate synthase

dethiobiotin synthase
ACP S-malonyltransferase
3-oxoacyl-ACP reductase FabG
NA
NA
NA
NA
4-(cytidine 5'-diphospho)-2-C-methyl-D-erythritol kinase
protoporphyrinogen oxidase HemJ
beta-ketoacyl-ACP synthase II
EcsC family protein
phosphomannomutase/phosphoglucomutase
bacteriohemerythrin
VOC family protein
transcriptional regulator
NA
MgtC/SapB family protein
dicarboxylate/amino acid:cation symporter
cold-shock protein
hypothetical protein
HK97-gp10 family putative phage morphogenesis protein
GatB/YqeY domain-containing protein
30S ribosomal protein S21
NA
NA
aspartate aminotransferase family protein
CoA-acylating methylmalonate-semialdehyde dehydrogenase
amino acid permease
DUF2171 domain-containing protein
acyl-CoA dehydrogenase
PIG-L family deacetylase
class I SAM-dependent methyltransferase
glycosyltransferase
NirD/YgiW/YdeI family stress tolerance protein
sodium/glutamate symporter
LysR substrate-binding domain-containing protein
hypothetical protein
TetR/AcrR family transcriptional regulator
Csu fimbrial major subunit CsuAB
Csu fimbrial biogenesis protein CsuA
Csu fimbrial biogenesis protein CsuB
Csu fimbrial biogenesis chaperone CsuC
Csu fimbrial usher CsuD
Csu fimbrial tip adhesin CsuE
lipocalin family protein
acyl-CoA desaturase
NAD(P)/FAD-dependent oxidoreductase
DUF1365 domain-containing protein
class I SAM-dependent methyltransferase

aspartate/glutamate racemase family protein
MFS transporter
hypothetical protein
hypothetical protein
circularly permuted type 2 ATP-grasp protein
alpha-E domain-containing protein
transglutaminase domain-containing protein
proteasome-type protease
50S ribosomal protein L9
30S ribosomal protein S18
30S ribosomal protein S6
amino acid permease
nuclear transport factor 2 family protein
gamma-aminobutyraldehyde dehydrogenase
GMC oxidoreductase
alpha/beta hydrolase
flavin-containing monooxygenase
ATP-dependent chaperone ClpB
CinA family protein
OmpA family protein
tRNA (guanosine(46)-N7)-methyltransferase TrmB
hypothetical protein
KTSC domain-containing protein
heavy metal translocating P-type ATPase
four-helix bundle copper-binding protein
hypothetical protein
exodeoxyribonuclease VII small subunit
exodeoxyribonuclease VII large subunit
hypothetical protein
hypothetical protein
minor capsid protein
MFS transporter
DUF4142 domain-containing protein
hypothetical protein
hypothetical protein
hypothetical protein
hypothetical protein
phage terminase large subunit
DUF2158 domain-containing protein
hypothetical protein
DUF4882 family protein
universal stress protein
EAL domain-containing protein
3-hydroxyacyl-CoA dehydrogenase NAD-binding domain-containing protein
hypothetical protein
BapA/Bap/LapF family large adhesin
universal stress protein
hypothetical protein
NA

glycerol-3-phosphate dehydrogenase
tRNA 2-thiocytidine(32) synthetase TtcA
transglycosylase SLT domain-containing protein
FdhF/YdeP family oxidoreductase
formate dehydrogenase accessory sulfurtransferase FdhD
phosphoribosylglycinamide formyltransferase
class 1 fructose-bisphosphatase
multidrug effflux MFS transporter
KGW motif small protein
NGG1p interacting factor NIF3
enoyl-CoA hydratase-related protein
CAP domain-containing protein
ornithine uptake porin CarO type 2
RBBP9/YdeN family alpha/beta hydrolase
sulfate ABC transporter substrate-binding protein
hypothetical protein
excinuclease ABC subunit UvrB
lipocalin family protein
tetratricopeptide repeat protein
3'(2'),5'-bisphosphate nucleotidase CysQ
30S ribosomal protein S20
NAD(P)H-binding protein
hypothetical protein
bifunctional mannitol-1-phosphate dehydrogenase/phosphatase
sulfite exporter TauE/SafE family protein
hypothetical protein
TonB C-terminal domain-containing protein
hypothetical protein
hypothetical protein
hypothetical protein
50S ribosomal protein L3 N(5)-glutamine methyltransferase
GGDEF domain-containing protein
transcriptional regulator
thiamine pyrophosphate-dependent dehydrogenase E1 component subunit alpha
alpha-ketoacid dehydrogenase subunit beta
2-oxo acid dehydrogenase subunit E2
dihydrolipoyl dehydrogenase
acetoin reductase
2,3-butanediol dehydrogenase
MBL fold metallo-hydrolase
TIGR01244 family sulfur transferase
sulfite exporter TauE/SafE family protein
EamA family transporter
MFS transporter
acetyl-CoA C-acyltransferase
TIGR00366 family protein
3-oxoacid CoA-transferase subunit B
CoA transferase subunit A
GntP family permease

3-hydroxybutyrate dehydrogenase
AraC family transcriptional regulator
molybdate ABC transporter substrate-binding protein
molybdate ABC transporter permease subunit
polyphosphate kinase 1
hypothetical protein
hypothetical protein
hypothetical protein
catalase family protein
DUF4882 family protein
fimbrial protein
fimbria/pilus outer membrane usher protein
fimbrial protein
DUF4468 domain-containing protein
hypothetical protein
hypothetical protein
endopeptidase La
META domain-containing protein
urease accessory protein UreE
hypothetical protein
copper resistance protein NlpE
isocitrate lyase
sulfate adenylyltransferase subunit CysD
YqiA/YcfP family alpha/beta fold hydrolase
carbonic anhydrase
hypothetical protein
inorganic phosphate transporter
alpha/beta hydrolase
MFS transporter
non-heme iron oxygenase ferredoxin subunit
YaeQ family protein
BCCT family transporter
transcriptional regulator BetI
betaine-aldehyde dehydrogenase
choline dehydrogenase
malate dehydrogenase (quinone)
homocysteine S-methyltransferase family protein
basic amino acid/polyamine antiporter
molecular chaperone
hypothetical protein
hypothetical protein
hypothetical protein
hypothetical protein
DNA breaking-rejoining protein
universal stress protein
chlorhexidine efflux PACE transporter AceI
acyl-CoA thioesterase
iron-containing alcohol dehydrogenase
iron-containing redox enzyme family protein

acetyl-CoA carboxylase biotin carboxyl carrier protein
GTP 3',8-cyclase MoaA
MoaD/ThiS family protein
molybdenum cofactor biosynthesis protein MoaE
NAD-dependent epimerase/dehydratase family protein
class II fumarate hydratase
hypothetical protein
D-amino acid dehydrogenase
S4 domain-containing protein
YcgJ family protein
hypothetical protein
S8 family peptidase
SOS response-associated peptidase family protein
universal stress protein
PEGA domain-containing protein
branched-chain amino acid transport system II carrier protein
CBS domain-containing protein
hypothetical protein
hypothetical protein
cyd operon YbgE family protein
cytochrome bd-I oxidase subunit CydX
3-oxoacid CoA-transferase subunit A
3-oxoacid CoA-transferase subunit B
3-oxoadipyl-CoA thiolase
3-carboxy-cis,cis-muconate cycloisomerase
3-oxoadipate enol-lactonase
MFS transporter
4-carboxymuconolactone decarboxylase
protocatechuate 3,4-dioxygenase subunit beta
protocatechuate 3,4-dioxygenase subunit alpha
type I 3-dehydroquinate dehydratase
aromatic-ring-hydroxylating dioxygenase subunit beta
p-hydroxyphenylacetate 3-hydroxylase reductase component
3-oxoadipate enol-lactonase
3-oxoadipyl-CoA thiolase
3-oxoacid CoA-transferase subunit B
catechol 1,2-dioxygenase
LysR substrate-binding domain-containing protein
alpha/beta hydrolase
saccharopine dehydrogenase NADP-binding domain-containing protein
type 1 glutamine amidotransferase domain-containing protein
putative quinol monooxygenase
hydrolase
glucose/sorbosone family PQQ-dependent dehydrogenase
hypothetical protein
DUF2314 domain-containing protein
Vi polysaccharide biosynthesis UDP-N-acetylglucosamine C-6 dehydrogenase TviB
glycosyltransferase family 4 protein
O-antigen polymerase

glycosyltransferase family 4 protein
sugar transferase
acetyltransferase
DegT/DnrJ/EryC1/StrS family aminotransferase
L-lactate permease
transcriptional regulator LldR
FMN-dependent L-lactate dehydrogenase LldD
D-lactate dehydrogenase
2-methylcitrate synthase
Fe/S-dependent 2-methylisocitrate dehydratase AcnD
D-amino acid dehydrogenase
alanine racemase
RidA family protein
amino acid permease
amino acid permease
CoA-acylating methylmalonate-semialdehyde dehydrogenase
3-hydroxyisobutyrate dehydrogenase
NA
NA
ATP synthase subunit I
MFS transporter
DUF2946 family protein
TonB-dependent copper receptor
hypothetical protein
poly-beta-1,6 N-acetyl-D-glucosamine exporter porin PgaA
poly-beta-1,6-N-acetyl-D-glucosamine N-deacetylase PgaB
poly-beta-1,6-N-acetyl-D-glucosamine synthase
poly-beta-1,6-N-acetyl-D-glucosamine biosynthesis protein PgaD
hypothetical protein
glycine zipper domain-containing protein
3-hydroxyacyl-CoA dehydrogenase
acyl-CoA dehydrogenase family protein
AMP-binding protein
potassium-transporting ATPase subunit KdpA
potassium-transporting ATPase subunit KdpB
potassium-transporting ATPase subunit KdpC
MFS transporter
hypothetical protein
hypothetical protein
GNAT family N-acetyltransferase
iron-containing alcohol dehydrogenase
hypothetical protein
spore coat U domain-containing protein
fatty acid oxidation complex subunit alpha FadB
acetyl-CoA C-acyltransferase FadA
RBBP9/YdeN family alpha/beta hydrolase
Naقی� antiporter NhaC family protein
DNÄ transfer protein p32
50S ribosomal protein L7/L12

50S ribosomal protein L10
NA
NA
NA
argininosuccinate lyase
oxidative damage protection protein
site-specific tyrosine recombinase XerD
DsbC family protein
DUF2726 domain-containing protein
arsenate reductase (glutaredoxin)
signal peptidase II
ISL3-like element ISPpu12 family transposase
NAD(P)/FAD-dependent oxidoreductase
multidrug efflux RND transporter permease subunit AdeB
multidrug efflux RND transporter periplasmic adaptor subunit AdeA
hypothetical protein
indoleacetamide hydrolase
helix-turn-helix transcriptional regulator
hypothetical protein
MFS transporter
HlyD family secretion protein
methionine synthase
DUF1852 domain-containing protein
acyl-CoA dehydrogenase
adenosine deaminase
NA
NA
NA
NA
gamma carbonic anhydrase family protein
ribosome-associated translation inhibitor RaiA
Mpo1-like protein
helix-turn-helix domain-containing protein
acyl-CoA dehydrogenase
lipase family protein
winged helix-turn-helix transcriptional regulator
zinc-dependent alcohol dehydrogenase
hypothetical protein
hypothetical protein
biotin carboxylase N-terminal domain-containing protein
5-oxoprolinase/urea amidolyase family protein
putative hydro-lyase
5-oxoprolinase subunit PxpA
NRAMP family divalent metal transporter
LysR family transcriptional regulator MumR
FAD-dependent oxidoreductase
hypothetical protein
M57 family metalloprotease
undecaprenyl-diphosphatase

TerC family protein
hypothetical protein
glutathionylspermidine synthase family protein
GlsB/YeaQ/YmgE family stress response membrane protein
membrane protein insertion efficiency factor YidD
putative solute-binding protein
tyrosine--tRNA ligase
aldehyde dehydrogenase family protein
Lrp/AsnC family transcriptional regulator
thiamine pyrophosphate-binding protein
amino acid permease
substrate-binding domain-containing protein
phosphate ABC transporter permease subunit PstC
5'/3'-nucleotidase SurE
50S ribosomal protein L31
elongation factor P
hotdog fold domain-containing protein
phosphopantetheine-binding protein
Mpo1-like protein
NADP-dependent isocitrate dehydrogenase
hypothetical protein
hypothetical protein
glycosyltransferase
hypothetical protein
hypothetical protein
RcnB family protein
hypothetical protein
organic hydroperoxide resistance protein
GFA family protein
DUF4256 domain-containing protein
30S ribosomal protein S9
50S ribosomal protein L13
tRNA guanosine(34) transglycosylase Tgt
preprotein translocase subunit YajC
protein translocase subunit SecF
NA
NA
NA
xanthine dehydrogenase molybdopterin binding subunit
