## Supplemental table 5 for "The peptide LyeTx I mnΔK induces transcriptomic reprogramming in a novel Multidrug-resistant *Acinetobacter baumannii*"

| | pgap_annot | log2FoldChange | pvalue | gene_name |
|---|---|---|---|---|
| 1 | cds-pgaptmp_000059 | -1,10152687129219 | 2,39812306055334e-16 | NA |
| 2 | cds-pgaptmp_000067 | -1,05343109691621 | 1,59418974411077e-13 | NA |
| 3 | cds-pgaptmp_000077 | 1,08491222620839 | 0,004435859637365 | NA |
| 4 | cds-pgaptmp_000078 | 1,79396765699938 | 1,67888023859398e-39 | NA |
| 5 | cds-pgaptmp_000079 | 1,92075428055637 | 7,94146060537e-27 | NA |
| 6 | cds-pgaptmp_000084 | -4,45393897707631 | 0 | NA |
| 7 | cds-pgaptmp_000085 | -3,02203518938749 | 9,1842291414991e-199 | NA |
| 8 | cds-pgaptmp_000089 | 1,28470311701318 | 3,25825108921838e-13 | NA |
| 9 | cds-pgaptmp_000097 | -1,23245668248979 | 1,75160431182877e-35 | NA |
| 10 | cds-pgaptmp_000098 | -1,12693471112892 | 4,14506942337598e-35 | NA |
| 11 | cds-pgaptmp_000099 | -1,18391084328459 | 3,21128861417768e-38 | NA |
| 12 | cds-pgaptmp_000100 | -1,23729404063881 | 1,74350403039199e-63 | NA |
| 13 | cds-pgaptmp_000125 | -1,80004840361717 | 2,53955928518136e-65 | NA |
| 14 | cds-pgaptmp_000154 | -1,42110360858232 | 8,17342981339926e-22 | NA |
| 15 | cds-pgaptmp_000155 | -1,77462850558809 | 8,50012365271966e-99 | NA |
| 16 | cds-pgaptmp_000156 | -1,96708721247959 | 2,71378654815715e-109 | cydB |
| 17 | cds-pgaptmp_000157 | -2,1672421715821 | 1,26722287189609e-28 | NA |
| 18 | cds-pgaptmp_000174 | 1,19993634857481 | 0,00170699332923208 | NA |
| 19 | cds-pgaptmp_000177 | -1,14928124399668 | 1,40256649708849e-51 | yghU |
| 20 | cds-pgaptmp_000190 | 1,05817835583302 | 2,92424869956232e-05 | NA |
| 21 | cds-pgaptmp_000191 | 1,52450001766007 | 1,2045992230288e-05 | NA |
| 22 | cds-pgaptmp_000202 | -4,90186405713166 | 4,15878944303801e-151 | NA |
| 23 | cds-pgaptmp_000203 | -1,45404044711378 | 2,40887167033343e-07 | NA |
| 24 | cds-pgaptmp_000204 | -4,69127576603224 | 0 | NA |
| 25 | cds-pgaptmp_000205 | -3,73551330622624 | 0 | katE |
| 26 | cds-pgaptmp_000206 | -2,36568823127853 | 7,66726015508006e-132 | NA |
| 27 | cds-pgaptmp_000207 | -1,91496458282391 | 5,43198995118162e-09 | NA |
| 28 | cds-pgaptmp_000208 | -6,35586467073899 | 8,77386100979987e-55 | NA |
| 29 | cds-pgaptmp_000209 | -5,42777891720549 | 1,03175421983811e-128 | surA1 |
| 30 | cds-pgaptmp_000210 | -1,37604133613686 | 4,90295211348421e-22 | NA |
| 31 | cds-pgaptmp_000211 | -1,67654929018278 | 5,28869400441254e-32 | NA |
| 32 | cds-pgaptmp_000212 | -4,14255878023437 | 7,29255060012139e-99 | NA |
| 33 | cds-pgaptmp_000216 | -1,02660038404475 | 2,21723278049973e-06 | NA |
| 34 | cds-pgaptmp_000217 | -1,80832629868312 | 7,79570990500088e-41 | NA |
| 35 | cds-pgaptmp_000218 | -1,45055259097768 | 1,05906557506752e-67 | NA |
| 36 | cds-pgaptmp_000219 | -1,63288759080993 | 2,11563919101728e-95 | NA |
| 37 | cds-pgaptmp_000220 | -1,45124493283399 | 5,40993983700259e-72 | NA |
| 38 | cds-pgaptmp_000221 | -1,40448937938904 | 8,78609165628166e-52 | NA |
| 39 | cds-pgaptmp_000222 | -1,2759307765683 | 1,49321951825622e-46 | NA |
| 40 | cds-pgaptmp_000223 | -1,4363369166175 | 2,43486186930496e-72 | NA |
| 41 | cds-pgaptmp_000224 | -1,3272778527412 | 1,17400798508425e-45 | NA |
| 42 | cds-pgaptmp_000226 | -1,57200871448538 | 2,10583630779857e-117 | NA |
| 43 | cds-pgaptmp_000255 | 1,96559804083343 | 4,68453094783107e-52 | pobA |
| 44 | cds-pgaptmp_000278 | -2,20558369861585 | 5,57007301820642e-08 | NA |
| 45 | cds-pgaptmp_000279 | -1,88167099069402 | 1,67739518135656e-17 | NA |
| 46 | cds-pgaptmp_000280 | -2,17112013549611 | 4,41682192692459e-22 | NA |
| 47 | cds-pgaptmp_000281 | -1,82050444254031 | 1,54020067621508e-23 | NA |
| 48 | cds-pgaptmp_000282 | -1,68546760633793 | 7,89037541797754e-07 | NA |

| | | | | |
|---|---|---|---|---|
| 49 | cds-pgaptmp_000283 | -1,67878923547564 | 1,65282600472059e-07 | NA |
| 50 | cds-pgaptmp_000284 | -1,33941841995852 | 2,053738195512e-20 | NA |
| 51 | cds-pgaptmp_000288 | 1,39091404964603 | 7,1974741847954e-52 | NA |
| 52 | cds-pgaptmp_000289 | 1,18066329894137 | 1,82168894308581e-46 | NA |
| 53 | cds-pgaptmp_000291 | -1,26146094701846 | 4,45478162543748e-39 | NA |
| 54 | cds-pgaptmp_000292 | -1,15138400546336 | 1,13485572761855e-48 | paaX |
| 55 | cds-pgaptmp_000293 | -1,86150357056723 | 5,84547239439764e-89 | paaK |
| 56 | cds-pgaptmp_000294 | -2,50885543520488 | 2,18254226488209e-140 | pcaF |
| 57 | cds-pgaptmp_000295 | -2,50456992307674 | 6,14960900034436e-144 | NA |
| 58 | cds-pgaptmp_000296 | -2,4570634937297 | 3,01855934838383e-139 | paaG |
| 59 | cds-pgaptmp_000297 | -2,26888462899965 | 7,03987484351257e-74 | NA |
| 60 | cds-pgaptmp_000298 | -1,95868975744625 | 5,67222680663571e-86 | paaE |
| 61 | cds-pgaptmp_000299 | -1,53014126057535 | 1,6054576075149e-45 | paaD |
| 62 | cds-pgaptmp_000300 | -1,69696506795598 | 1,34567403422257e-55 | paaC |
| 63 | cds-pgaptmp_000301 | -1,71345525439826 | 6,68299897707635e-64 | paaB |
| 64 | cds-pgaptmp_000302 | -1,47474244919237 | 3,96690839326465e-65 | paaA |
| 65 | cds-pgaptmp_000303 | -1,34229424656778 | 5,19920986445033e-46 | paaZ |
| 66 | cds-pgaptmp_001552 | -2,20442783768059 | 1,03532429118933e-162 | NA |
| 67 | cds-pgaptmp_001578 | -1,23084566612076 | 1,04561697909055e-39 | NA |
| 68 | cds-pgaptmp_001590 | 1,02931261599673 | 3,45715334980913e-46 | NA |
| 69 | cds-pgaptmp_001598 | -1,52502986660374 | 1,3680892972617e-24 | NA |
| 70 | cds-pgaptmp_001617 | 1,028600008664 | 4,3191217615856e-28 | adeC |
| 71 | cds-pgaptmp_001618 | 1,23993087093405 | 2,35358727979211e-37 | NA |
| 72 | cds-pgaptmp_001619 | 1,9260812137757 | 1,14367029703607e-78 | NA |
| 73 | cds-pgaptmp_001635 | -1,03892633859877 | 4,95565913993337e-52 | acnA |
| 74 | cds-pgaptmp_001657 | -1,76193378105187 | 7,54152094826251e-73 | NA |
| 75 | cds-pgaptmp_001660 | -1,1941136454745 | 1,0227575958445e-18 | NA |
| 76 | cds-pgaptmp_001661 | -1,13851580119523 | 8,92842640511312e-48 | NA |
| 77 | cds-pgaptmp_001662 | -1,01429585794497 | 2,152630026557e-23 | NA |
| 78 | cds-pgaptmp_001668 | -1,96389465719916 | 1,29877334868476e-80 | NA |
| 79 | cds-pgaptmp_001669 | -2,01660483654423 | 7,83117334107361e-44 | NA |
| 80 | cds-pgaptmp_001670 | -1,99827156305027 | 4,915917945118e-73 | NA |
| 81 | cds-pgaptmp_002086 | -1,3383645804937 | 2,39715543525875e-128 | NA |
| 82 | cds-pgaptmp_002111 | -1,00926853190781 | 2,88648979658117e-48 | NA |
| 83 | cds-pgaptmp_002120 | 1,79238540547319 | 3,699970829576e-115 | lolA |
| 84 | cds-pgaptmp_002126 | 1,30017884299731 | 4,62433222225845e-82 | NA |
| 85 | cds-pgaptmp_002168 | -1,38986837890857 | 2,59031328566159e-72 | NA |
| 86 | cds-pgaptmp_002202 | -1,99031372979556 | 6,33222698444495e-12 | NA |
| 87 | cds-pgaptmp_002208 | 1,09463393562825 | 1,7181821775823e-42 | NA |
| 88 | cds-pgaptmp_002209 | 1,27154294220636 | 6,05979882970443e-27 | NA |
| 89 | cds-pgaptmp_002217 | 1,08783002190536 | 3,44975343131427e-10 | NA |
| 90 | cds-pgaptmp_002221 | 1,13353652868894 | 4,34189491630826e-53 | NA |
| 91 | cds-pgaptmp_002259 | 1,0923486863044 | 9,78678156810628e-21 | NA |
| 92 | cds-pgaptmp_002279 | 5,14457232232854 | 2,98478203477645e-275 | NA |
| 93 | cds-pgaptmp_002283 | 5,94900851645158 | 3,37403036588474e-165 | NA |
| 94 | cds-pgaptmp_002289 | 1,3138378871564 | 7,48449817610735e-15 | NA |
| 95 | cds-pgaptmp_002301 | -1,41334955673189 | 3,49322685353901e-72 | NA |
| 96 | cds-pgaptmp_002304 | -1,9201367254229 | 8,08955278716541e-39 | NA |
| 97 | cds-pgaptmp_002305 | -1,98124792349164 | 1,51954737370986e-39 | NA |

| | | | | |
|---|---|---|---|---|
| 98 | cds-pgaptmp_002306 | -1,71529688824634 | 1,11681235824484e-23 | gpM |
| 99 | cds-pgaptmp_002319 | -2,35492716271103 | 1,49895132586012e-57 | NA |
| 100 | cds-pgaptmp_002320 | -2,29757561576859 | 1,27414235463035e-37 | NA |
| 101 | cds-pgaptmp_002321 | -1,78764945449242 | 1,59312934728421e-13 | NA |
| 102 | cds-pgaptmp_002322 | -1,35260638576797 | 0,00068967941980782 | NA |
| 103 | cds-pgaptmp_002432 | 1,18495295122372 | 2,29464591534508e-68 | NA |
| 104 | cds-pgaptmp_002504 | 1,22942997474446 | 2,95639203660619e-70 | NA |
| 105 | cds-pgaptmp_002509 | -1,58516375017615 | 5,23064289206957e-106 | NA |
| 106 | cds-pgaptmp_002516 | 1,26547027614646 | 2,78852188869909e-89 | NA |
| 107 | cds-pgaptmp_002529 | 1,27048562876472 | 8,54050468680412e-16 | NA |
| 108 | cds-pgaptmp_002534 | 2,0132641215637 | 1,40099185460384e-102 | NA |
| 109 | cds-pgaptmp_002545 | 1,75923295051494 | 3,48345052925748e-67 | NA |
| 110 | cds-pgaptmp_002560 | -1,49214759696079 | 3,92083213311454e-73 | NA |
| 111 | cds-pgaptmp_002596 | 1,30608773877683 | 7,1543569574047e-11 | NA |
| 112 | cds-pgaptmp_002597 | 1,28680478699225 | 3,1049003517291e-15 | NA |
| 113 | cds-pgaptmp_002598 | 1,12555200499692 | 6,61740660238321e-09 | NA |
| 114 | cds-pgaptmp_002609 | 1,07985766024947 | 6,04320480541392e-12 | NA |
| 115 | cds-pgaptmp_002615 | 1,08226480161418 | 3,84744707768762e-19 | NA |
| 116 | cds-pgaptmp_002630 | 2,01187337932272 | 4,78135843361173e-129 | NA |
| 117 | cds-pgaptmp_002657 | 1,54999245029734 | 1,48023167957403e-13 | NA |
| 118 | cds-pgaptmp_002658 | 1,78761190385794 | 1,73729065541413e-102 | NA |
| 119 | cds-pgaptmp_002659 | 1,34233782655411 | 1,96774263887458e-87 | NA |
| 120 | cds-pgaptmp_002680 | 1,02152010644545 | 6,11502705365898e-15 | NA |
| 121 | cds-pgaptmp_002694 | -3,173373238361 | 0 | NA |
| 122 | cds-pgaptmp_002708 | -1,34989201183405 | 5,79186979081913e-69 | NA |
| 123 | cds-pgaptmp_002709 | -1,54103845339765 | 5,56476709959291e-67 | gabT |
| 124 | cds-pgaptmp_002711 | -1,17354435379205 | 1,8618595361e-25 | NA |
| 125 | cds-pgaptmp_002725 | 1,57036967471811 | 3,86297710128226e-79 | omp33-36 |
| 126 | cds-pgaptmp_002731 | -2,60275553670241 | 1,87388848214732e-208 | NA |
| 127 | cds-pgaptmp_002783 | 1,04019996367967 | 7,3096310033682e-27 | NA |
| 128 | cds-pgaptmp_002790 | 1,34106901436152 | 2,20944487495844e-12 | NA |
| 129 | cds-pgaptmp_002791 | 1,18464370197172 | 3,84993782354028e-16 | NA |
| 130 | cds-pgaptmp_002803 | 2,05953551356371 | 1,31791148798129e-71 | NA |
| 131 | cds-pgaptmp_002839 | 1,1019427423999 | 9,31321796443451e-23 | NA |
| 132 | cds-pgaptmp_002858 | -1,63453535260797 | 1,70066588108886e-53 | NA |
| 133 | cds-pgaptmp_002860 | -2,54205565986169 | 2,93213144213312e-136 | NA |
| 134 | cds-pgaptmp_002903 | 1,23067866941331 | 2,06973073359655e-14 | NA |
| 135 | cds-pgaptmp_002911 | 1,14053844356877 | 3,02540693881902e-42 | NA |
| 136 | cds-pgaptmp_002935 | -1,03560740658819 | 2,25696712031197e-23 | NA |
| 137 | cds-pgaptmp_002936 | -2,24862372154453 | 4,52147993927067e-265 | NA |
| 138 | cds-pgaptmp_002946 | -1,43921014571456 | 6,33029666112356e-61 | NA |
| 139 | cds-pgaptmp_002948 | 1,02531888508289 | 4,61262157159859e-25 | NA |
| 140 | cds-pgaptmp_002960 | 1,0957056225052 | 5,11786081870898e-34 | NA |
| 141 | cds-pgaptmp_002968 | -1,84616456702154 | 4,44178774379557e-97 | NA |
| 142 | cds-pgaptmp_002969 | -5,06529471168115 | 0 | NA |
| 143 | cds-pgaptmp_002970 | -3,33338519576772 | 1,3730736932724e-148 | otsB |
| 144 | exon-pgaptmp_002998-1 | 1,1340119993131 | 2,3680187043687e-37 | NA |
| 145 | cds-pgaptmp_003001 | 1,02549829253862 | 1,11814668576419e-36 | lolB |
| 146 | exon-pgaptmp_003012-1 | -1,06061851561369 | 7,25980583201205e-31 | NA |

| | | | | |
|---|---|---|---|---|
| 147 | cds-pgaptmp_003055 | 2,08719712918268 | 3,44660972863427e-67 | NA |
| 148 | cds-pgaptmp_003089 | 1,62243657657974 | 8,54170569745452e-87 | NA |
| 149 | cds-pgaptmp_003090 | 1,37087866613045 | 7,87305560986712e-57 | NA |
| 150 | cds-pgaptmp_003091 | 1,07507153323802 | 6,35602914974099e-37 | qatC |
| 151 | cds-pgaptmp_003094 | 1,01033523943495 | 0,00179693304438044 | NA |
| 152 | cds-pgaptmp_003102 | -1,38379362845353 | 2,37852803557437e-06 | NA |
| 153 | cds-pgaptmp_003107 | 1,13895787663654 | 0,000311448133315418 | NA |
| 154 | cds-pgaptmp_003130 | -1,12091961716741 | 3,02056440683656e-25 | NA |
| 155 | cds-pgaptmp_003132 | 1,04927372885608 | 9,66567962024625e-17 | NA |
| 156 | cds-pgaptmp_003135 | -1,16943818035543 | 2,38225471089931e-19 | hemP |
| 157 | cds-pgaptmp_003252 | -4,29479239013585 | 7,50758210516162e-76 | NA |
| 158 | cds-pgaptmp_003253 | -1,24574949321253 | 6,66088519008055e-25 | NA |
| 159 | cds-pgaptmp_003254 | -1,87132408075264 | 2,31372468193422e-68 | NA |
| 160 | cds-pgaptmp_003255 | -2,2946148602628 | 4,24766287193093e-58 | NA |
| 161 | cds-pgaptmp_003256 | -2,51358305341055 | 3,47538292229943e-77 | NA |
| 162 | cds-pgaptmp_003257 | -2,70384616985931 | 2,1166884578209e-85 | NA |
| 163 | cds-pgaptmp_003263 | 1,09427949568472 | 1,06404806588537e-44 | gltS |
| 164 | cds-pgaptmp_003271 | 1,13287298344177 | 0,00022096339068713 | NA |
| 165 | cds-pgaptmp_003272 | 1,39900867909469 | 2,85604852720406e-06 | NA |
| 166 | cds-pgaptmp_003275 | 1,04253135788515 | 7,28327812376937e-12 | csuB |
| 167 | cds-pgaptmp_003276 | 1,17689988831519 | 2,21589595086488e-30 | csuC |
| 168 | cds-pgaptmp_003277 | 1,02844363749051 | 1,30441517101112e-36 | csuD |
| 169 | cds-pgaptmp_003278 | 1,00765309754378 | 3,80243929027808e-20 | csuE |
| 170 | cds-pgaptmp_003285 | 1,31340107429226 | 2,20616884318066e-16 | NA |
| 171 | cds-pgaptmp_003294 | 1,69518275681272 | 7,26304772882746e-10 | NA |
| 172 | cds-pgaptmp_003295 | 1,59395545018187 | 1,43454339298554e-07 | NA |
| 173 | cds-pgaptmp_003296 | 1,06951876695927 | 0,000302432721292348 | NA |
| 174 | cds-pgaptmp_003310 | 1,48700361721857 | 6,08887855354644e-22 | NA |
| 175 | cds-pgaptmp_003323 | -2,08274979119676 | 8,03123765661831e-150 | NA |
| 176 | cds-pgaptmp_003326 | -1,28157977480559 | 1,25376066509107e-44 | NA |
| 177 | cds-pgaptmp_003361 | 1,34110247577674 | 6,86309255426518e-13 | NA |
| 178 | cds-pgaptmp_003362 | 1,02340105935855 | 0,000164637297610482 | NA |
| 179 | cds-pgaptmp_003402 | -1,64447505012683 | 3,29482637615103e-35 | NA |
| 180 | cds-pgaptmp_003435 | -1,47354185487776 | 1,85679615165713e-76 | NA |
| 181 | cds-pgaptmp_003439 | -1,35273515483467 | 8,47035188830214e-33 | NA |
| 182 | cds-pgaptmp_003456 | -3,20524032145972 | 1,54206762285385e-119 | NA |
| 183 | cds-pgaptmp_003457 | -2,27098273203792 | 2,03456679116656e-72 | NA |
| 184 | exon-pgaptmp_003462-1 | -1,3222404379727 | 8,27666024317194e-38 | NA |
| 185 | cds-pgaptmp_003463 | -1,41916015553173 | 1,26098615605035e-35 | NA |
| 186 | cds-pgaptmp_003472 | -1,40115419601775 | 3,83488040056128e-29 | NA |
| 187 | cds-pgaptmp_003474 | -1,59673427089062 | 8,09380463967944e-31 | NA |
| 188 | cds-pgaptmp_003479 | -3,95494820282813 | 0 | NA |
| 189 | cds-pgaptmp_003486 | -1,47600021726279 | 1,61887298057422e-75 | NA |
| 190 | cds-pgaptmp_003490 | -1,14585259030978 | 7,82371042270627e-05 | terL |
| 191 | cds-pgaptmp_003493 | 2,56912618133624 | 5,34262688067429e-86 | NA |
| 192 | cds-pgaptmp_003494 | 2,05395215489805 | 7,21945522054855e-76 | NA |
| 193 | cds-pgaptmp_003503 | 1,91276372311368 | 2,5222268261055e-90 | NA |
| 194 | cds-pgaptmp_003529 | 1,24646526771942 | 0,0103172144523283 | NA |
| 195 | cds-pgaptmp_003532 | 1,42733430308161 | 5,16799543732837e-33 | NA |

| | | | | |
|---|---|---|---|---|
| 196 | cds-pgaptmp_003540 | -2,11316618567969 | 4,52148606924537e-272 | NA |
| 197 | cds-pgaptmp_003541 | 1,02525648291172 | 1,59214482121325e-22 | NA |
| 198 | cds-pgaptmp_003566 | 1,04363963562232 | 2,63942265248243e-25 | glpK |
| 199 | cds-pgaptmp_003571 | 1,06691181115272 | 1,19474491684916e-71 | blp1 |
| 200 | cds-pgaptmp_003589 | 2,06845437249745 | 0,000655194725104285 | NA |
| 201 | cds-pgaptmp_003658 | -1,25660054046974 | 3,1105735509778e-55 | NA |
| 202 | cds-pgaptmp_003659 | -1,01262425668738 | 1,18850409135904e-09 | NA |
| 203 | cds-pgaptmp_003676 | 1,29766596641111 | 1,53302147217671e-15 | NA |
| 204 | cds-pgaptmp_000337 | -1,13773291638084 | 3,92502950900681e-23 | NA |
| 205 | cds-pgaptmp_000364 | -1,05890321666898 | 6,85882909176594e-12 | NA |
| 206 | cds-pgaptmp_000365 | -1,3762843272951 | 1,65683661446573e-13 | NA |
| 207 | cds-pgaptmp_000371 | -1,19558704777153 | 1,73487035444896e-28 | NA |
| 208 | cds-pgaptmp_000372 | -1,20445338648834 | 3,55768944410615e-52 | NA |
| 209 | cds-pgaptmp_000375 | -1,47400002867872 | 2,4866555143504e-44 | mtlD |
| 210 | cds-pgaptmp_000396 | -1,04626877472017 | 5,18409824028691e-21 | NA |
| 211 | cds-pgaptmp_000420 | -1,32410904104953 | 8,61500639706313e-08 | NA |
| 212 | cds-pgaptmp_000422 | -1,21510808909299 | 5,54628289005848e-09 | NA |
| 213 | cds-pgaptmp_000423 | -1,21624545932755 | 3,99586707116051e-09 | lpdA |
| 214 | cds-pgaptmp_000425 | -1,14699888026428 | 8,4732555175116e-24 | NA |
| 215 | cds-pgaptmp_000428 | -1,98319585951477 | 3,774235146546e-62 | NA |
| 216 | cds-pgaptmp_000429 | -1,52972509669675 | 1,5137772147558e-79 | NA |
| 217 | cds-pgaptmp_000432 | -1,74316945738153 | 6,37940882878711e-42 | NA |
| 218 | cds-pgaptmp_000452 | -1,27137825207004 | 1,84950169385901e-58 | NA |
| 219 | cds-pgaptmp_000453 | -1,177548219745 | 1,21280901175588e-16 | NA |
| 220 | cds-pgaptmp_000454 | -1,96128245237626 | 2,48953286291622e-71 | NA |
| 221 | cds-pgaptmp_000455 | -1,73054362264247 | 2,52528680281195e-65 | NA |
| 222 | cds-pgaptmp_000458 | -1,15554955173586 | 1,8942156836353e-27 | NA |
| 223 | cds-pgaptmp_000459 | -1,10942005923151 | 2,24843358874301e-26 | NA |
| 224 | cds-pgaptmp_000486 | -1,22012755762374 | 4,04116082990915e-11 | NA |
| 225 | cds-pgaptmp_000490 | -1,49638411543375 | 3,67325668643409e-34 | NA |
| 226 | cds-pgaptmp_000494 | 1,28367750377808 | 1,5253551362188e-23 | NA |
| 227 | cds-pgaptmp_000498 | 1,10406432264829 | 8,27093134287284e-11 | NA |
| 228 | cds-pgaptmp_000499 | 1,10616179268874 | 9,21790503989241e-08 | NA |
| 229 | cds-pgaptmp_000528 | -1,3071112936027 | 0,000312854486448108 | NA |
| 230 | cds-pgaptmp_000542 | -1,5357454249168 | 2,69801299596528e-28 | NA |
| 231 | cds-pgaptmp_000545 | 1,06632919844167 | 4,10576083365045e-06 | NA |
| 232 | cds-pgaptmp_000568 | 1,04106503722272 | 6,18939630923574e-08 | NA |
| 233 | cds-pgaptmp_000569 | 1,29708219835574 | 9,36398056479723e-08 | NA |
| 234 | cds-pgaptmp_000576 | 1,27464694979961 | 1,28531618578899e-101 | NA |
| 235 | cds-pgaptmp_000614 | 1,40833639495158 | 0,00507293847771129 | NA |
| 236 | cds-pgaptmp_000656 | 1,478687333049 | 2,6431473211927e-07 | NA |
| 237 | cds-pgaptmp_000657 | 1,21918364513271 | 1,99531254987898e-08 | NA |
| 238 | cds-pgaptmp_000658 | 2,66485025212273 | 3,69397760039104e-15 | NA |
| 239 | cds-pgaptmp_000659 | 2,68606090653106 | 3,60320073033597e-09 | NA |
| 240 | cds-pgaptmp_000676 | 2,20206212427454 | 4,99183906566453e-90 | betI |
| 241 | cds-pgaptmp_000677 | 2,26817718664264 | 3,50274662290445e-136 | betB |
| 242 | cds-pgaptmp_000678 | 2,68542732673853 | 4,58643159814369e-245 | betA |
| 243 | cds-pgaptmp_000695 | 1,02380460579291 | 5,85807831905298e-05 | NA |
| 244 | cds-pgaptmp_000696 | 1,00547620202426 | 3,97444519972892e-06 | NA |

| | | | | |
|---|---|---|---|---|
| 245 | cds-pgaptmp_000698 | -1,46497667850607 | 7,51888109340764e-40 | NA |
| 246 | cds-pgaptmp_000701 | -2,74269094664158 | 5,49598449815857e-115 | NA |
| 247 | cds-pgaptmp_000707 | -1,28087857903625 | 6,92475101596945e-05 | NA |
| 248 | cds-pgaptmp_000710 | -1,44399952019644 | 1,25201563645822e-61 | NA |
| 249 | cds-pgaptmp_000712 | 1,76732308918578 | 0,000777602139320283 | NA |
| 250 | cds-pgaptmp_000773 | -1,80615809498842 | 6,51364648430709e-175 | NA |
| 251 | cds-pgaptmp_000810 | -1,43666538064571 | 2,33629664339222e-36 | NA |
| 252 | cds-pgaptmp_000812 | -1,27837833833593 | 5,3192421258921e-40 | NA |
| 253 | cds-pgaptmp_000816 | 1,70403869164104 | 3,0366188121654e-38 | NA |
| 254 | cds-pgaptmp_000838 | -1,14679347006282 | 1,8328385181915e-50 | NA |
| 255 | cds-pgaptmp_000839 | -1,05295030013604 | 1,24449337868586e-46 | cydX |
| 256 | cds-pgaptmp_000840 | -1,03456812086802 | 2,98612167056896e-48 | cydB |
| 257 | cds-pgaptmp_000841 | -1,07333569719513 | 2,3864189089129e-52 | NA |
| 258 | cds-pgaptmp_000863 | 1,24887563776558 | 2,5178532377499e-31 | NA |
| 259 | cds-pgaptmp_000864 | 1,07509315477514 | 1,22332520376787e-10 | NA |
| 260 | cds-pgaptmp_000930 | -1,73859993799142 | 3,54062670652454e-122 | NA |
| 261 | cds-pgaptmp_000935 | -2,23524016176796 | 7,46235699688773e-92 | NA |
| 262 | cds-pgaptmp_000938 | 1,49422686242782 | 1,33543479570351e-16 | NA |
| 263 | cds-pgaptmp_000939 | 1,17557119588497 | 4,15869746054497e-11 | NA |
| 264 | cds-pgaptmp_000947 | -1,31332968078795 | 1,22870137754159e-23 | NA |
| 265 | cds-pgaptmp_000948 | -1,15377063912609 | 1,06383027174101e-41 | NA |
| 266 | cds-pgaptmp_000962 | 1,13811101544543 | 5,50176665369345e-31 | NA |
| 267 | cds-pgaptmp_000975 | 1,37968496563636 | 8,61369209880175e-67 | NA |
| 268 | cds-pgaptmp_000976 | 1,76817921892791 | 5,59184874662404e-152 | NA |
| 269 | cds-pgaptmp_000977 | 1,14842770229606 | 1,32496612849645e-91 | NA |
| 270 | cds-pgaptmp_000978 | 1,21965521919601 | 4,36758798993434e-74 | NA |
| 271 | cds-pgaptmp_000979 | 1,13305668258865 | 1,18677564616065e-48 | NA |
| 272 | cds-pgaptmp_000980 | 1,01434428027136 | 4,96589835947309e-58 | NA |
| 273 | cds-pgaptmp_000987 | 5,84813782175311 | 0 | lldP |
| 274 | cds-pgaptmp_000988 | 3,64426669565119 | 2,80770000767986e-303 | lldR |
| 275 | cds-pgaptmp_000989 | 3,38547558451987 | 5,47340478369038e-248 | lldD |
| 276 | cds-pgaptmp_000990 | 3,08810081465287 | 1,62655487737835e-288 | dld |
| 277 | cds-pgaptmp_001003 | 1,40729056603347 | 2,65413935485733e-47 | NA |
| 278 | cds-pgaptmp_001026 | 1,59069699748156 | 1,48994389289608e-06 | abaI |
| 279 | cds-pgaptmp_001027 | 1,35000684388003 | 1,23993356748104e-07 | NA |
| 280 | cds-pgaptmp_001029 | 1,73505616518371 | 8,95800093365863e-12 | NA |
| 281 | cds-pgaptmp_001030 | 1,46522640119942 | 1,75172460064159e-10 | NA |
| 282 | cds-pgaptmp_001032 | 1,31763632903748 | 6,71557017648331e-17 | NA |
| 283 | cds-pgaptmp_001072 | 1,52320278892421 | 1,42487046849746e-49 | NA |
| 284 | cds-pgaptmp_001075 | 1,45100570613685 | 6,725534338318e-41 | NA |
| 285 | cds-pgaptmp_001084 | 1,3835742833325 | 5,37191174905553e-31 | NA |
| 286 | cds-pgaptmp_001093 | 4,94749330955088 | 8,73329286264165e-131 | NA |
| 287 | cds-pgaptmp_001094 | 3,06320840894589 | 1,96416508463852e-210 | NA |
| 288 | cds-pgaptmp_001102 | -2,18886055050779 | 2,72678994199563e-91 | NA |
| 289 | cds-pgaptmp_001103 | -2,02455879339281 | 5,64907455822259e-44 | NA |
| 290 | cds-pgaptmp_001104 | -1,54670719893288 | 1,33862223878905e-42 | NA |
| 291 | cds-pgaptmp_001126 | 1,05615551152123 | 1,16343817257173e-06 | NA |
| 292 | cds-pgaptmp_001128 | -1,04838469340461 | 1,12025133125858e-36 | NA |
| 293 | cds-pgaptmp_001130 | -1,23890719900794 | 1,95272452334964e-26 | NA |

| | | | | |
|---|---|---|---|---|
| 294 | cds-pgaptmp_001149 | 1,07185305148007 | 5,00560116545507e-28 | NA |
| 295 | cds-pgaptmp_001152 | 1,32102738992049 | 4,92816726945459e-35 | NA |
| 296 | cds-pgaptmp_001159 | -1,71416994960115 | 1,04185440608264e-49 | NA |
| 297 | cds-pgaptmp_001163 | -2,42347733821363 | 5,06431794644587e-90 | NA |
| 298 | cds-pgaptmp_001168 | 1,20274739594768 | 5,69554483992208e-08 | NA |
| 299 | cds-pgaptmp_001175 | 1,85330951607541 | 6,74992873162271e-54 | NA |
| 300 | cds-pgaptmp_001176 | 1,6915233571792 | 1,40338942180814e-07 | NA |
| 301 | cds-pgaptmp_001210 | 1,14020219771457 | 2,86443539390358e-07 | NA |
| 302 | cds-pgaptmp_001222 | 1,27707500875389 | 2,87482607381779e-30 | xerD |
| 303 | cds-pgaptmp_001279 | 1,47694356272541 | 2,31213807219445e-128 | NA |
| 304 | cds-pgaptmp_001328 | 1,02494457649337 | 2,5714959674246e-08 | NA |
| 305 | cds-pgaptmp_001329 | 1,07120740302078 | 2,69491326317056e-06 | NA |
| 306 | cds-pgaptmp_001330 | 1,48667142182272 | 1,73327973411609e-09 | NA |
| 307 | cds-pgaptmp_001331 | 1,1233144899334 | 1,85305293857096e-07 | NA |
| 308 | cds-pgaptmp_001349 | 1,86807633473078 | 4,70276066553083e-66 | NA |
| 309 | cds-pgaptmp_001350 | 2,26105766379297 | 5,1270411076626e-126 | NA |
| 310 | cds-pgaptmp_001372 | 1,10031902328827 | 4,73382513161676e-11 | NA |
| 311 | cds-pgaptmp_001408 | -1,55749416958363 | 8,45197738010354e-83 | raiA |
| 312 | cds-pgaptmp_001711 | 1,13055576661374 | 4,47135607932996e-27 | NA |
| 313 | cds-pgaptmp_001770 | -3,81407365829097 | 1,72529974776256e-218 | NA |
| 314 | cds-pgaptmp_001773 | 1,3571172602442 | 5,82471332147602e-27 | NA |
| 315 | cds-pgaptmp_001774 | 1,2154701011052 | 5,8083550465541e-22 | NA |
| 316 | cds-pgaptmp_001777 | -1,18252623613914 | 4,83622983941066e-31 | NA |
| 317 | cds-pgaptmp_001778 | -1,98961934130829 | 5,60806205569937e-95 | NA |
| 318 | cds-pgaptmp_001779 | -1,63511130638631 | 6,04092212630454e-77 | NA |
| 319 | cds-pgaptmp_001780 | -1,68136671875105 | 2,72517085417765e-78 | NA |
| 320 | cds-pgaptmp_001781 | -1,792232422125 | 1,61232612087836e-109 | pxpA |
| 321 | cds-pgaptmp_001782 | -1,92229842539425 | 8,05334074646813e-165 | NA |
| 322 | cds-pgaptmp_001804 | 1,0025208982181 | 1,19488746598826e-08 | NA |
| 323 | cds-pgaptmp_001833 | 1,40896808179647 | 4,55507689692559e-40 | NA |
| 324 | cds-pgaptmp_001841 | 2,7022728230831 | 4,6020690454703e-26 | NA |
| 325 | cds-pgaptmp_001842 | 1,04073319819052 | 2,4962207765773e-33 | NA |
| 326 | cds-pgaptmp_001856 | 1,20288185702933 | 9,1334407612498e-19 | NA |
| 327 | cds-pgaptmp_001859 | 1,33047808757601 | 1,37778647138808e-21 | NA |
| 328 | cds-pgaptmp_001863 | -1,2604461141428 | 7,98567109807421e-42 | NA |
| 329 | cds-pgaptmp_001880 | -1,19322032182243 | 2,24618496479169e-66 | NA |
| 330 | cds-pgaptmp_001890 | -2,03213056297077 | 3,34749946051046e-207 | NA |
| 331 | cds-pgaptmp_001892 | -2,2989626325422 | 2,30239081221379e-141 | NA |
| 332 | cds-pgaptmp_001893 | -1,97212893802731 | 4,3250124584668e-53 | NA |
| 333 | cds-pgaptmp_001935 | -1,11128708009109 | 3,50712459603993e-13 | NA |
| 334 | cds-pgaptmp_001958 | -1,2721861702262 | 5,10338901079647e-61 | NA |
| 335 | cds-pgaptmp_002005 | 6,16328574368554 | 0 | NA |
| 336 | cds-pgaptmp_002006 | 1,2223965955656 | 1,16486964926869e-38 | NA |
| 337 | cds-pgaptmp_002018 | 2,33724619437 | 1,44915253715227e-35 | NA |
| 338 | cds-pgaptmp_002037 | -1,13763210049135 | 2,31157199586919e-34 | NA |
| 339 | cds-pgaptmp_002370 | 1,04826440947539 | 1,90943679728366e-26 | NA |

protein
hypothetical protein
hypothetical protein
hypothetical protein
fimbrial protein
molecular chaperone
DMT family transporter
hypothetical protein
tyrosine-type recombinase/integrase
amino acid ABC transporter ATP-binding protein
amino acid ABC transporter permease
amino acid ABC transporter permease
amino acid ABC transporter substrate-binding protein
dicarboxylate/amino acid:cation symporter
hypothetical protein
cytochrome ubiquinol oxidase subunit I
cytochrome d ubiquinol oxidase subunit II
DUF2474 domain-containing protein
CidA/LrgA family protein
glutathione-dependent disulfide-bond oxidoreductase
amino acid ABC transporter ATP-binding protein
amino acid ABC transporter permease
KGG domain-containing protein
DUF6367 family protein
SDR family oxidoreductase
catalase HPII
iron-containing redox enzyme family protein
CinA family protein
hypothetical protein
surface antigen protein 1
hypothetical protein
hypothetical protein
hypothetical protein
DcaP family trimeric outer membrane transporter
class I SAM-dependent methyltransferase
AMP-binding protein
TetR/AcrR family transcriptional regulator
isovaleryl-CoA dehydrogenase
carboxyl transferase domain-containing protein
enoyl-CoA hydratase/isomerase family protein
acetyl-CoA carboxylase biotin carboxylase subunit
hydroxymethylglutaryl-CoA lyase
indolepyruvate ferredoxin oxidoreductase family protein
4-hydroxybenzoate 3-monooxygenase
hypothetical protein
hypothetical protein
hypothetical protein
cold-shock protein
hypothetical protein

hypothetical protein
hypothetical protein
ATP-dependent helicase
AAA family ATPase
DapH/DapD/GlmU-related protein
phenylacetic acid degradation operon negative regulatory protein PaaX
phenylacetate--CoA ligase PaaK
3-oxoadipyl-CoA thiolase
3-hydroxyacyl-CoA dehydrogenase
2-(1,2-epoxy-1,2-dihydrophenyl)acetyl-CoA isomerase PaaG
enoyl-CoA hydratase-related protein
1,2-phenylacetyl-CoA epoxidase subunit PaaE
1,2-phenylacetyl-CoA epoxidase subunit PaaD
1,2-phenylacetyl-CoA epoxidase subunit PaaC
1,2-phenylacetyl-CoA epoxidase subunit PaaB
1,2-phenylacetyl-CoA epoxidase subunit PaaA
phenylacetic acid degradation bifunctional protein PaaZ
coniferyl aldehyde dehydrogenase
TonB-dependent receptor
gluconokinase
phosphatase PAP2 family protein
AdeC/AdeK/OprM family multidrug efflux complex outer membrane factor
MacB family efflux pump subunit
MacA family efflux pump subunit
aconitate hydratase AcnA
heavy metal translocating P-type ATPase
hypothetical protein
multicopper oxidase domain-containing protein
copper resistance protein B
Re/Si-specific NAD(P)() transhydrogenase subunit alpha
proton-translocating transhydrogenase family protein
NAD(P)() transhydrogenase (Re/Si-specific) subunit beta
Ig-like domain-containing protein
hypothetical protein
outer membrane lipoprotein chaperone LolA
phosphatase PAP2 family protein
PA1571 family protein
entericidin A/B family lipoprotein
methyl-accepting chemotaxis protein
chemotaxis protein CheW
zinc ribbon domain-containing protein YjdM
DsbC family protein
MBL fold metallo-hydrolase
hypothetical protein
RcnB family protein
stealth conserved region 3 domain-containing protein
HAMP domain-containing sensor histidine kinase
GPO family capsid scaffolding protein
phage major capsid protein, P2 family

phage terminase small subunit
phage tail sheath protein
phage major tail tube protein
phage tail assembly protein
GpE family phage tail protein
DUF3108 domain-containing protein
phospholipid-binding protein MlaC
DEAD/DEAH box helicase
acyl-CoA dehydrogenase C-terminal domain-containing protein
LysR family transcriptional regulator
hypothetical protein
hypothetical protein
hemerythrin domain-containing protein
pilus assembly protein PilP
type 4a pilus biogenesis protein PilO
PilN domain-containing protein
sulfate ABC transporter substrate-binding protein
hypothetical protein
SulP family inorganic anion transporter
LysE family translocator
DNA/RNA non-specific endonuclease
hypothetical protein
hypothetical protein
MFS transporter
NAD-dependent succinate-semialdehyde dehydrogenase
4-aminobutyrate--2-oxoglutarate transaminase
amino acid permease
porin Omp33-36
hypothetical protein
outer membrane protein OmpK
SH3 domain-containing protein
lysozyme inhibitor LprI family protein
SDR family NAD(P)-dependent oxidoreductase
acyltransferase family protein
PQQ-dependent sugar dehydrogenase
DUF1328 domain-containing protein
hypothetical protein
hypothetical protein
hypothetical protein
mechanosensitive ion channel
hypothetical protein
5'-methylthioadenosine/S-adenosylhomocysteine nucleosidase
pantothenate kinase
MFS transporter
trehalose-6-phosphate synthase
trehalose-phosphatase
NA
lipoprotein insertase outer membrane protein LolB
NA

bacteriohemerythrin
P-loop NTPase fold protein
hypothetical protein
Qat anti-phage system QueC-like protein QatC
hypothetical protein
Slam-dependent surface lipoprotein
hypothetical protein
MgtC/SapB family protein
NirD/YgiW/YdeI family stress tolerance protein
hemin uptake protein HemP
DUF2171 domain-containing protein
acyl-CoA dehydrogenase
PIG-L family deacetylase
class I SAM-dependent methyltransferase
glycosyltransferase
NirD/YgiW/YdeI family stress tolerance protein
sodium/glutamate symporter
hypothetical protein
TetR/AcrR family transcriptional regulator
Csu fimbrial biogenesis protein CsuB
Csu fimbrial biogenesis chaperone CsuC
Csu fimbrial usher CsuD
Csu fimbrial tip adhesin CsuE
DMT family transporter
lipocalin family protein
acyl-CoA desaturase
NAD(P)/FAD-dependent oxidoreductase
MFS transporter
hypothetical protein
circularly permuted type 2 ATP-grasp protein
hypothetical protein
DUF4265 domain-containing protein
hypothetical protein
hypothetical protein
KTSC domain-containing protein
four-helix bundle copper-binding protein
hypothetical protein
NA
transposase
hypothetical protein
minor capsid protein
DUF4142 domain-containing protein
hypothetical protein
phage terminase large subunit
DUF2158 domain-containing protein
hypothetical protein
hypothetical protein
TetR/AcrR family transcriptional regulator
exo-alpha-sialidase

BapA/Bap/LapF family large adhesin
poly alpha-glucosyltransferase
glycerol kinase GlpK
biofilm-associated Ig-like repeat protein Blp1
hypothetical protein
multidrug effflux MFS transporter
KGW motif small protein
sulfate ABC transporter substrate-binding protein
NAD(P)H-binding protein
SidA/IucD/PvdA family monooxygenase
DHA2 family efflux MFS transporter permease subunit
hypothetical protein
PepSY-associated TM helix domain-containing protein
bifunctional mannitol-1-phosphate dehydrogenase/phosphatase
hypothetical protein
thiamine pyrophosphate-dependent dehydrogenase E1 component subunit alpha
2-oxo acid dehydrogenase subunit E2
dihydrolipoyl dehydrogenase
2,3-butanediol dehydrogenase
MBL fold metallo-hydrolase
TIGR01244 family sulfur transferase
EamA family transporter
acetyl-CoA C-acyltransferase
TIGR00366 family protein
3-oxoacid CoA-transferase subunit B
CoA transferase subunit A
GntP family permease
3-hydroxybutyrate dehydrogenase
hypothetical protein
hypothetical protein
DUF4882 family protein
fimbrial protein
hypothetical protein
Ish1 domain-containing protein
hypothetical protein
hypothetical protein
hypothetical protein
hypothetical protein
META domain-containing protein
hypothetical protein
SDR family oxidoreductase
Rieske 2Fe-2S domain-containing protein
recombinase-like helix-turn-helix domain-containing protein
non-heme iron oxygenase ferredoxin subunit
transcriptional regulator BetI
betaine-aldehyde dehydrogenase
choline dehydrogenase
hypothetical protein
hypothetical protein

TonB-dependent receptor
hypothetical protein
hypothetical protein
universal stress protein
hypothetical protein
D-amino acid dehydrogenase
SOS response-associated peptidase family protein
universal stress protein
PEGA domain-containing protein
cyd operon YbgE family protein
cytochrome bd-I oxidase subunit CydX
cytochrome d ubiquinol oxidase subunit II
cytochrome ubiquinol oxidase subunit I
FAD-dependent oxidoreductase
nuclear transport factor 2 family protein
alpha/beta hydrolase
type 1 glutamine amidotransferase domain-containing protein
MBL fold metallo-hydrolase
TetR family transcriptional regulator
putative quinol monooxygenase
hydrolase
DUF2314 domain-containing protein
glycosyltransferase family 4 protein
O-antigen polymerase
glycosyltransferase family 4 protein
sugar transferase
acetyltransferase
DegT/DnrJ/EryC1/StrS family aminotransferase
L-lactate permease
transcriptional regulator LldR
FMN-dependent L-lactate dehydrogenase LldD
D-lactate dehydrogenase
SDR family NAD(P)-dependent oxidoreductase
acyl-homoserine-lactone synthase AbaI
DUF4902 domain-containing protein
fatty acyl-AMP ligase
acyl-CoA dehydrogenase
amino acid adenylation domain-containing protein
hypothetical protein
MFS transporter
hypothetical protein
hypothetical protein
glycine zipper domain-containing protein
3-hydroxyacyl-CoA dehydrogenase
acyl-CoA dehydrogenase family protein
AMP-binding protein
hypothetical protein
LysR family transcriptional regulator
hypothetical protein

aldehyde dehydrogenase family protein
iron-containing alcohol dehydrogenase
spore coat U domain-containing protein
RBBP9/YdeN family alpha/beta hydrolase
selenocysteine synthase
YMGG-like glycine zipper-containing protein
DNA transfer protein p32
TetR/AcrR family transcriptional regulator
site-specific tyrosine recombinase XerD
hypothetical protein
GH3 auxin-responsive promoter family protein
hypothetical protein
DHA2 family efflux MFS transporter permease subunit
HlyD family secretion protein
MFS transporter
HlyD family secretion protein
zinc-binding dehydrogenase
ribosome-associated translation inhibitor RaiA
OmpW family outer membrane protein
zinc-dependent alcohol dehydrogenase
hypothetical protein
hypothetical protein
chloride channel protein
biotin carboxylase N-terminal domain-containing protein
5-oxoprolinase/urea amidolyase family protein
putative hydro-lyase
5-oxoprolinase subunit PxpA
NRAMP family divalent metal transporter
LysE/ArgO family amino acid transporter
FAD-dependent oxidoreductase
hypothetical protein
hypothetical protein
undecaprenyl-diphosphatase
glutathionylspermidine synthase family protein
GlsB/YeaQ/YmgE family stress response membrane protein
putative solute-binding protein
aldehyde dehydrogenase family protein
thiamine pyrophosphate-binding protein
amino acid permease
Mpo1-like protein
hypothetical protein
RcnB family protein
RcnB family protein
hypothetical protein
hypothetical protein
hypothetical protein
