## Supplemental table 6 for "The peptide LyeTx I mnΔK induces transcriptomic reprogramming in a novel Multidrug-resistant *Acinetobacter baumannii*"

### 9 hours treatment

| | pgap_annot | log2FoldChan | pvalue | gene_name | protein |
|---|---|---|---|---|---|
| 1 | cds-pgaptmp_ | -1,466830173 | 1,6039282263 | NA | malate synthase G |
| 2 | cds-pgaptmp_ | 1,2119028601 | 4,3755659796 | NA | hypothetical protein |
| 3 | cds-pgaptmp_ | -1,239995791 | 2,0877988703 | putA | trifunctional transcriptional re |
| 4 | cds-pgaptmp_ | -1,365053455 | 4,4843958840 | NA | hypothetical protein |
| 5 | cds-pgaptmp_ | 1,2950639999 | 2,1198964949 | NA | fimbria/pilus outer membrane |
| 6 | cds-pgaptmp_ | 1,2567640894 | 2,7896395121 | NA | fimbrial protein |
| 7 | cds-pgaptmp_ | 1,1954618898 | 4,0529549922 | NA | DcaP family trimeric outer m |
| 8 | cds-pgaptmp_ | -1,017144346 | 7,8929553926 | NA | amino acid ABC transporter |
| 9 | cds-pgaptmp_ | -1,367635790 | 3,8633572958 | NA | glutathione S-transferase fan |
| 10 | cds-pgaptmp_ | -1,438982525 | 3,5452960792 | NA | tautomerase family protein |
| 11 | cds-pgaptmp_ | -1,221534389 | 2,3559751230 | NA | glutathione peroxidase |
| 12 | cds-pgaptmp_ | -1,077400104 | 3,9290400533 | msrB | peptide-methionine (R)-S-oxi |
| 13 | cds-pgaptmp_ | -1,822374279 | 2,5962222200 | tauD | taurine dioxygenase |
| 14 | cds-pgaptmp_ | -2,282635116 | 3,7692336244 | tauC | taurine ABC transporter perm |
| 15 | cds-pgaptmp_ | -2,636114333 | 5,0381407340 | NA | taurine ABC transporter ATP |
| 16 | cds-pgaptmp_ | -2,573182004 | 6,9372797834 | tauA | taurine ABC transporter subs |
| 17 | cds-pgaptmp_ | 1,0981231869 | 0,0022936400 | NA | malonate decarboxylase sub |
| 18 | cds-pgaptmp_ | 1,6606475422 | 4,2948636070 | NA | triphosphoribosyl-dephospho |
| 19 | cds-pgaptmp_ | 1,1847388631 | 1,9636427019 | mdcA | malonate decarboxylase sub |
| 20 | cds-pgaptmp_ | -1,061363201 | 4,1180539478 | NA | LrgB family protein |
| 21 | cds-pgaptmp_ | -1,174094842 | 2,1896683672 | NA | CidA/LrgA family protein |
| 22 | cds-pgaptmp_ | -1,433538475 | 1,6654412196 | NA | cysteine ABC transporter sub |
| 23 | cds-pgaptmp_ | -1,487971625 | 9,0135106336 | NA | transporter substrate-binding |
| 24 | cds-pgaptmp_ | -1,751020619 | 1,2384888898 | NA | amino acid ABC transporter |
| 25 | cds-pgaptmp_ | -1,616242829 | 4,2213422983 | NA | amino acid ABC transporter |
| 26 | cds-pgaptmp_ | -1,438398634 | 1,5440106617 | NA | amino acid ABC transporter |
| 27 | cds-pgaptmp_ | 3,9541439307 | 0,0005244400 | NA | hypothetical protein |
| 28 | cds-pgaptmp_ | -1,800740633 | 8,7953435576 | NA | LysE family translocator |
| 29 | cds-pgaptmp_ | 1,6949662900 | 3,0256405554 | NA | CinA family protein |
| 30 | cds-pgaptmp_ | -1,824410110 | 1,0534012162 | NA | hypothetical protein |
| 31 | cds-pgaptmp_ | -1,138914541 | 9,0382018505 | NA | 3-hydroxyacyl-CoA dehydrog |
| 32 | cds-pgaptmp_ | -1,218963165 | 5,1799431622 | paaG | 2-(1,2-epoxy-1,2-dihydrophe |
| 33 | cds-pgaptmp_ | -1,089431547 | 2,8638876383 | NA | enoyl-CoA hydratase-related |
| 34 | cds-pgaptmp_ | -1,240773871 | 1,8431278216 | paaE | 1,2-phenylacetyl-CoA epoxid |
| 35 | cds-pgaptmp_ | -1,257324404 | 8,4774123636 | paaD | 1,2-phenylacetyl-CoA epoxid |
| 36 | cds-pgaptmp_ | -1,599615830 | 2,3711437319 | paaC | 1,2-phenylacetyl-CoA epoxid |
| 37 | cds-pgaptmp_ | -1,418867582 | 2,8221449299 | paaB | 1,2-phenylacetyl-CoA epoxid |
| 38 | cds-pgaptmp_ | -1,485238131 | 1,5881123995 | paaA | 1,2-phenylacetyl-CoA epoxid |
| 39 | cds-pgaptmp_ | -1,470980281 | 1,4896767507 | paaZ | phenylacetic acid degradatio |
| 40 | cds-pgaptmp_ | -1,047230278 | 1,4780708738 | NA | epoxyqueuosine reductase C |
| 41 | cds-pgaptmp_ | -1,156923542 | 1,5200794272 | NA | GNAT family N-acetyltransfe |
| 42 | cds-pgaptmp_ | -1,522845662 | 2,9471489607 | NA | pyrimidine/purine nucleoside |
| 43 | cds-pgaptmp_ | 1,0747063947 | 3,0674243661 | infB | translation initiation factor IF- |
| 44 | cds-pgaptmp_ | -1,473606590 | 4,4875241006 | gspG | type II secretion system majc |
| 45 | cds-pgaptmp_ | -2,222344681 | 1,2884819780 | NA | hypothetical protein |
| 46 | cds-pgaptmp_ | -1,119446161 | 1,8607275154 | hcaR | DNA-binding transcriptional r |
| 47 | cds-pgaptmp_ | 1,3745898547 | 2,3640375198 | NA | energy transducer TonB |
| 48 | cds-pgaptmp_ | 1,3995001768 | 6,6609246538 | NA | MotA/TolQ/ExbB proton char |

| | | | | | |
|---|---|---|---|---|---|
| 49 | cds-pgaptmp_ | 1,2033395463 | 1,9492375660 | NA | ExbD/TolR family protein |
| 50 | cds-pgaptmp_ | 1,5921360976 | 3,4537985408 | tatC | twin-arginine translocase sub |
| 51 | cds-pgaptmp_ | 1,9118779658 | 1,3603119394 | tatB | Sec-independent protein trar |
| 52 | cds-pgaptmp_ | 1,7673449266 | 1,5685250117 | NA | Sec-independent protein trar |
| 53 | cds-pgaptmp_ | 1,0182330336 | 1,0093396020 | NA | TonB-dependent receptor |
| 54 | cds-pgaptmp_ | 2,4814518412 | 9,6178047172 | adeC | AdeC/AdeK/OprM family mul |
| 55 | cds-pgaptmp_ | 2,4034351990 | 5,8557289550 | NA | MacB family efflux pump sub |
| 56 | cds-pgaptmp_ | 2,0147433615 | 1,2945299367 | NA | MacA family efflux pump sub |
| 57 | cds-pgaptmp_ | 1,0743319591 | 0,0022873902 | NA | hypothetical protein |
| 58 | cds-pgaptmp_ | -1,001807738 | 9,9622178033 | NA | NAD(P)-dependent oxidored |
| 59 | cds-pgaptmp_ | -1,180888795 | 3,6484056896 | NA | neutral zinc metallopeptidase |
| 60 | cds-pgaptmp_ | -1,485409593 | 4,3621194784 | NA | winged helix-turn-helix trans |
| 61 | cds-pgaptmp_ | 2,3259143662 | 5,3784563725 | lolA | outer membrane lipoprotein |
| 62 | cds-pgaptmp_ | -1,051458917 | 6,6547893604 | NA | YbjQ family protein |
| 63 | cds-pgaptmp_ | -1,699935511 | 1,6057500358 | NA | DcaP family trimeric outer m |
| 64 | cds-pgaptmp_ | -1,289126464 | 2,1005120061 | hchA | glyoxalase III HchA |
| 65 | cds-pgaptmp_ | -1,207843639 | 2,9310907928 | NA | PaaX family transcriptional re |
| 66 | cds-pgaptmp_ | -1,388526425 | 2,4664534696 | NA | OmpP1/FadL family transpor |
| 67 | cds-pgaptmp_ | -1,849387811 | 4,5426140134 | NA | DJ-1/PfpI family protein |
| 68 | cds-pgaptmp_ | 1,8303209125 | 1,1260238218 | NA | PepSY domain-containing pr |
| 69 | cds-pgaptmp_ | 1,0971680879 | 1,7692205168 | NA | YdcF family protein |
| 70 | exon-pgaptmp | 1,6674837382 | 3,1020965811 | NA | NA |
| 71 | exon-pgaptmp | 1,8400535204 | 1,0300134011 | NA | NA |
| 72 | exon-pgaptmp | 1,8057289795 | 1,5849342638 | NA | NA |
| 73 | exon-pgaptmp | 1,7706302510 | 2,5312833532 | NA | NA |
| 74 | exon-pgaptmp | 1,7795849264 | 1,5329669902 | NA | NA |
| 75 | exon-pgaptmp | 1,5347188812 | 6,5134784903 | NA | NA |
| 76 | cds-pgaptmp_ | -1,113137868 | 4,1135934533 | NA | entericidin A/B family lipopro |
| 77 | cds-pgaptmp_ | -1,280004855 | 1,3448257690 | NA | response regulator |
| 78 | cds-pgaptmp_ | -1,382123407 | 3,8917877901 | pilG | twitching motility response re |
| 79 | cds-pgaptmp_ | -1,222891035 | 1,6553578356 | mscL | large conductance mechanos |
| 80 | cds-pgaptmp_ | 1,8905179442 | 6,0388100289 | NA | RtcB family protein |
| 81 | cds-pgaptmp_ | 1,6299403265 | 9,6943702217 | NA | hypothetical protein |
| 82 | cds-pgaptmp_ | -1,030709295 | 2,3054582052 | NA | carbohydrate porin |
| 83 | cds-pgaptmp_ | -1,191868842 | 4,0444177344 | NA | hemolysin III family protein |
| 84 | cds-pgaptmp_ | 1,0402858353 | 4,6505576381 | baeS | sensor histidine kinase efflux |
| 85 | cds-pgaptmp_ | 2,2285498870 | 0 | NA | hypothetical protein |
| 86 | cds-pgaptmp_ | 2,9428582323 | 0 | NA | RcnB family protein |
| 87 | cds-pgaptmp_ | 1,1407093587 | 9,6466427553 | NA | DUF4184 family protein |
| 88 | cds-pgaptmp_ | 1,5459571826 | 2,5890988292 | NA | DUF3108 domain-containing |
| 89 | cds-pgaptmp_ | -1,265807613 | 3,4983883090 | NA | YihY family inner membrane |
| 90 | cds-pgaptmp_ | 1,0326675974 | 1,4103928569 | NA | nitroreductase family protein |
| 91 | cds-pgaptmp_ | -2,736078000 | 2,2939278223 | NA | beta strand repeat-containing |
| 92 | cds-pgaptmp_ | 1,2240867762 | 2,9172413894 | NA | acyl-CoA dehydrogenase far |
| 93 | cds-pgaptmp_ | 1,0052365975 | 4,0566946176 | rpoA | DNA-directed RNA polymera |
| 94 | cds-pgaptmp_ | 1,0982093679 | 1,3036656803 | secY | preprotein translocase subur |
| 95 | cds-pgaptmp_ | 1,1273825105 | 5,4630344356 | rplO | 50S ribosomal protein L15 |
| 96 | cds-pgaptmp_ | 1,2762119362 | 2,0436828420 | rpmD | 50S ribosomal protein L30 |
| 97 | cds-pgaptmp_ | 1,0193289050 | 6,6385289977 | rpsE | 30S ribosomal protein S5 |

9 hours treatment

| | | |
|---|---|---|
| 98 | cds-pgaptmp_1,02438991711,0755047157rplR | 50S ribosomal protein L18 |
| 99 | cds-pgaptmp_1,01696731246,8579296628rplF | 50S ribosomal protein L6 |
| 100 | cds-pgaptmp_1,15596391006,8596947670rpsH | 30S ribosomal protein S8 |
| 101 | cds-pgaptmp_1,10232982903,8699545455rpsN | 30S ribosomal protein S14 |
| 102 | cds-pgaptmp_1,06816600621,8541022113rplE | 50S ribosomal protein L5 |
| 103 | cds-pgaptmp_1,03334092564,4758104003rplX | 50S ribosomal protein L24 |
| 104 | cds-pgaptmp_1,02114566104,0291143722rplN | 50S ribosomal protein L14 |
| 105 | cds-pgaptmp_1,14937791079,0698292740rpsQ | 30S ribosomal protein S17 |
| 106 | cds-pgaptmp_1,21047141674,9062749879rpmC | 50S ribosomal protein L29 |
| 107 | cds-pgaptmp_1,26330732984,5948192074rplP | 50S ribosomal protein L16 |
| 108 | cds-pgaptmp_1,28918749451,9337919422rpsC | 30S ribosomal protein S3 |
| 109 | cds-pgaptmp_1,39893786663,1562456318rplV | 50S ribosomal protein L22 |
| 110 | cds-pgaptmp_1,40378107961,3417238681rpsS | 30S ribosomal protein S19 |
| 111 | cds-pgaptmp_1,25287767691,1069489085rplB | 50S ribosomal protein L2 |
| 112 | cds-pgaptmp_1,16670087767,0576843469rplW | 50S ribosomal protein L23 |
| 113 | cds-pgaptmp_1,23064637956,6021571477rplD | 50S ribosomal protein L4 |
| 114 | cds-pgaptmp_1,34114500831,5157318078rplC | 50S ribosomal protein L3 |
| 115 | cds-pgaptmp_1,07394690352,6861601086rpsJ | 30S ribosomal protein S10 |
| 116 | cds-pgaptmp_-1,272773637 8,8877659749NA | helicase HerA-like domain-co |
| 117 | cds-pgaptmp_-1,816709724 2,3188209292NA | globin domain-containing pro |
| 118 | cds-pgaptmp_-1,680747822 3,1764259335NA | Rrf2 family transcriptional reg |
| 119 | cds-pgaptmp_1,34369356091,2896850217NA | STAS domain-containing pro |
| 120 | cds-pgaptmp_1,30440334982,4279548785NA | phospholipid-binding protein |
| 121 | cds-pgaptmp_1,14550808251,9870659462NA | DEAD/DEAH box helicase |
| 122 | cds-pgaptmp_-1,586859920 1,2557351200hemF | oxygen-dependent coproporp |
| 123 | cds-pgaptmp_-1,101436707 1,8350593142aroE | shikimate dehydrogenase |
| 124 | cds-pgaptmp_-1,317604932 1,3108820476NA | hypothetical protein |
| 125 | cds-pgaptmp_1,10218733822,6840045262NA | HPP family protein |
| 126 | cds-pgaptmp_1,58224583521,9992348489NA | hypothetical protein |
| 127 | cds-pgaptmp_-1,034190027 7,4493065626NA | Glu/Leu/Phe/Val dehydrogen |
| 128 | cds-pgaptmp_-1,090102270 2,0045635444NA | hypothetical protein |
| 129 | cds-pgaptmp_1,63042294872,6421401136NA | hypothetical protein |
| 130 | cds-pgaptmp_-1,727292724 2,6774042757NA | ParA family protein |
| 131 | cds-pgaptmp_-1,191275078 2,5274383565NA | AI-2E family transporter |
| 132 | cds-pgaptmp_1,19743942081,1830942061NA | MFS transporter |
| 133 | cds-pgaptmp_1,78958898954,6722917929NA | lipase secretion chaperone |
| 134 | cds-pgaptmp_1,52593382061,2373187636NA | lipase family alpha/beta hydr |
| 135 | cds-pgaptmp_1,38783513969,0804196885NA | Tfp pilus assembly protein Fi |
| 136 | cds-pgaptmp_-2,346412818 3,5228383690bfr | bacterioferritin |
| 137 | cds-pgaptmp_-1,091260453 4,0030830533aroK | shikimate kinase AroK |
| 138 | exon-pgaptmp1,05804942631,8169573080NA | NA |
| 139 | cds-pgaptmp_-1,645461564 8,2739995614NA | tetratricopeptide repeat prote |
| 140 | cds-pgaptmp_-1,151587615 3,2364325863NA | aminotransferase class I/II-fo |
| 141 | cds-pgaptmp_-1,999075995 2,0727389605NA | acetyl-CoA hydrolase/transfe |
| 142 | cds-pgaptmp_-1,733062315 2,7148619952NA | DUF805 domain-containing p |
| 143 | cds-pgaptmp_-1,139378912 1,1910782091NA | hypothetical protein |
| 144 | cds-pgaptmp_2,37332673235,9888495370NA | hypothetical protein |
| 145 | cds-pgaptmp_-1,255927788 3,9805476470ycaC | isochorismate family cysteine |
| 146 | cds-pgaptmp_-1,507676063 1,2143252235NA | AzlC family ABC transporter |

| | | |
|---|---|---|
| 147 | cds-pgaptmp_-1,164791885 6,0693349239NA | AzlD domain-containing prot |
| 148 | cds-pgaptmp_1,04540551315,1475390999NA | GGDEF domain-containing |
| 149 | cds-pgaptmp_-1,734875014 8,0945966468NA | DUF485 domain-containing |
| 150 | cds-pgaptmp_-1,706237587 5,1600510390NA | hypothetical protein |
| 151 | cds-pgaptmp_-1,035270114 8,2813504189NA | cysteine hydrolase family pro |
| 152 | cds-pgaptmp_1,06453247891,1770753615NA | TonB-dependent siderophore |
| 153 | cds-pgaptmp_2,29975613064,0458853202NA | TonB-dependent receptor |
| 154 | cds-pgaptmp_-1,082307706 2,9446611146NA | arsenate reductase |
| 155 | cds-pgaptmp_-1,372351851 6,3270464669NA | DUF2147 domain-containing |
| 156 | cds-pgaptmp_-3,233268786 5,2764487559NA | HPF/RaiA family ribosome-a |
| 157 | cds-pgaptmp_-1,819113078 9,3343104089NA | alpha/beta fold hydrolase |
| 158 | cds-pgaptmp_-1,183928501 3,4075322765pncB | nicotinate phosphoribosyltran |
| 159 | cds-pgaptmp_1,12674381564,2119002590hppD | 4-hydroxyphenylpyruvate dio |
| 160 | cds-pgaptmp_-1,463547922 3,3175020406NA | HIT family protein |
| 161 | cds-pgaptmp_1,54415298533,2828504534NA | hypothetical protein |
| 162 | cds-pgaptmp_-1,923890487 1,4737284494ppc | phosphoenolpyruvate carbox |
| 163 | cds-pgaptmp_1,39761168127,5137749524NA | TetR/AcrR family transcriptio |
| 164 | cds-pgaptmp_-1,567074330 2,2776117485NA | hypothetical protein |
| 165 | cds-pgaptmp_-1,375572499 1,2528722773NA | mechanosensitive ion chann |
| 166 | cds-pgaptmp_-2,887306523 1,6264359441bfr | bacterioferritin |
| 167 | cds-pgaptmp_-1,848062186 1,9995401146lysM | peptidoglycan-binding protei |
| 168 | exon-pgaptmp1,06529194181,3163516465NA | NA |
| 169 | exon-pgaptmp1,19949735862,3558021466NA | NA |
| 170 | cds-pgaptmp_1,30029103733,6273569223lolB | lipoprotein insertase outer m |
| 171 | cds-pgaptmp_1,12676586549,3021715176NA | porin |
| 172 | cds-pgaptmp_1,06371829691,4152759707fusA | elongation factor G |
| 173 | cds-pgaptmp_-1,024958139 2,1948062027rhtC | threonine export protein Rht |
| 174 | cds-pgaptmp_-1,143482761 4,7717970615NA | class II glutamine amidotrans |
| 175 | cds-pgaptmp_-2,516728904 4,4949615641NA | bacteriohemerythrin |
| 176 | cds-pgaptmp_1,20442163436,7225899938NA | VOC family protein |
| 177 | cds-pgaptmp_1,21533676020,0004134536NA | hypothetical protein |
| 178 | cds-pgaptmp_1,43185283256,5564926437NA | MFS transporter |
| 179 | cds-pgaptmp_1,09513741819,5481737736NA | ABC transporter permease |
| 180 | cds-pgaptmp_-1,443255043 5,4453628400NA | YfhL family 4Fe-4S dicluster |
| 181 | cds-pgaptmp_-1,689734995 7,5681272376yegQ | tRNA 5-hydroxyuridine modif |
| 182 | cds-pgaptmp_1,87552377601,2252494162NA | hypothetical protein |
| 183 | cds-pgaptmp_1,02846901178,1588765810NA | M48 family metalloprotease |
| 184 | cds-pgaptmp_-1,229543564 1,7861840074csuAB | Csu fimbrial major subunit C |
| 185 | cds-pgaptmp_1,13682372370,0011266658NA | acyl-CoA dehydrogenase fan |
| 186 | cds-pgaptmp_1,14117647937,7076490635NA | flavin-containing monooxyge |
| 187 | cds-pgaptmp_-1,031660667 2,4060884259NA | glutathione S-transferase |
| 188 | cds-pgaptmp_-1,766768215 3,5048301666NA | glutathione binding-like prote |
| 189 | cds-pgaptmp_-1,976060182 0,0006742687NA | heavy-metal-associated dom |
| 190 | cds-pgaptmp_-1,384761752 0,0061600245NA | heavy metal translocating P- |
| 191 | cds-pgaptmp_-1,173878425 0,0039942692cueR | Cu(I)-responsive transcriptio |
| 192 | cds-pgaptmp_-1,389760848 0,0019437260NA | GFA family protein |
| 193 | cds-pgaptmp_1,04753410244,9403365641NA | transposase |
| 194 | cds-pgaptmp_1,01848472480,0006304471NA | hypothetical protein |
| 195 | cds-pgaptmp_1,49804196187,8854697919NA | MFS transporter |

| | | | |
|---|---|---|---|
| 196 | cds-pgaptmp_1,10051357061,9269196466 | NA | putative peptidoglycan-bindir |
| 197 | cds-pgaptmp_1,66096739630,0010499350 | NA | DUF2213 domain-containing |
| 198 | cds-pgaptmp_1,15382045836,8226450180 | NA | TolC family protein |
| 199 | cds-pgaptmp_-1,156662809 9,1255346875 | NA | universal stress protein |
| 200 | cds-pgaptmp_1,44044878282,9878356073 | NA | D-Ala-D-Ala carboxypeptidas |
| 201 | cds-pgaptmp_1,35308188526,4919620114 | NA | TetR/AcrR family transcriptio |
| 202 | cds-pgaptmp_-1,507977862 4,3370326971 | NA | DUF2147 domain-containing |
| 203 | exon-pgaptmp-1,105561566 0,0028161887 | NA | NA |
| 204 | cds-pgaptmp_1,28747943514,4709171116 | NA | alkaline phosphatase D famil |
| 205 | cds-pgaptmp_-1,312098223 1,4010690913 | NA | hypothetical protein |
| 206 | cds-pgaptmp_-1,050059538 9,6968061262 | NA | MFS transporter |
| 207 | cds-pgaptmp_-1,647891178 1,4184670733 | NA | MBL fold metallo-hydrolase |
| 208 | cds-pgaptmp_1,29841987444,8611439517 | NA | multidrug efflux RND transpc |
| 209 | cds-pgaptmp_1,97061915491,8302413993 | NA | transglycosylase SLT domair |
| 210 | cds-pgaptmp_-1,177244120 5,3585796124 | NA | amino acid permease |
| 211 | cds-pgaptmp_1,06567111493,2575384374 | NA | lipoprotein-releasing ABC tra |
| 212 | cds-pgaptmp_1,03088467991,5879546739 | lolD | lipoprotein-releasing ABC tra |
| 213 | cds-pgaptmp_-1,148690500 7,7786102009 | NA | deoxyguanosinetriphosphate |
| 214 | cds-pgaptmp_-1,169478678 8,2108515551 | NA | enoyl-CoA hydratase-related |
| 215 | cds-pgaptmp_-1,143047025 1,2863070296 | cysW | sulfate ABC transporter perm |
| 216 | cds-pgaptmp_-1,403444500 7,5333901335 | cysT | sulfate ABC transporter perm |
| 217 | cds-pgaptmp_-1,922117955 5,1153278668 | NA | RBBP9/YdeN family alpha/be |
| 218 | cds-pgaptmp_-1,974734870 5,3760765767 | NA | sulfate ABC transporter subs |
| 219 | cds-pgaptmp_-1,881543875 1,0288513525 | NA | hypothetical protein |
| 220 | cds-pgaptmp_-1,118808246 1,2860542668 | NA | HU family DNA-binding prote |
| 221 | cds-pgaptmp_1,03177051931,2108202250 | NA | acyl-CoA dehydrogenase far |
| 222 | cds-pgaptmp_1,48976174873,1636416959 | NA | IucA/IucC family protein |
| 223 | cds-pgaptmp_1,21699063741,8467062410 | NA | TonB-dependent receptor |
| 224 | cds-pgaptmp_-1,382959289 9,1440730455 | NA | sulfite exporter TauE/SafE fa |
| 225 | cds-pgaptmp_1,50841558855,8426741266 | NA | hypothetical protein |
| 226 | cds-pgaptmp_1,49647459942,0549188755 | NA | TonB-dependent siderophore |
| 227 | cds-pgaptmp_1,24520702916,6005419874 | NA | porin |
| 228 | cds-pgaptmp_1,09525478503,8729733367 | lipA | lipoyl synthase |
| 229 | cds-pgaptmp_1,31724153028,0345850912 | NA | dienelactone hydrolase famil |
| 230 | cds-pgaptmp_1,00097177200,0008285300 | NA | SDR family oxidoreductase |
| 231 | cds-pgaptmp_1,29552961943,6047028524 | antA | anthranilate 1,2-dioxygenase |
| 232 | cds-pgaptmp_-1,155834507 2,4215317025 | modA | molybdate ABC transporter s |
| 233 | cds-pgaptmp_1,54048139940,0001703911 | NA | hypothetical protein |
| 234 | cds-pgaptmp_-1,097795085 5,3723003372 | NA | fimbrial protein |
| 235 | cds-pgaptmp_1,43300976590,0001031829 | NA | hypothetical protein |
| 236 | cds-pgaptmp_-1,909220473 0,0004737943 | NA | hypothetical protein |
| 237 | cds-pgaptmp_-1,106945420 1,0083993848 | NA | urease subunit beta |
| 238 | cds-pgaptmp_-1,754730775 1,6035120848 | NA | isocitrate lyase |
| 239 | cds-pgaptmp_-1,046960879 1,3004053841 | NA | DUF2239 family protein |
| 240 | cds-pgaptmp_-1,259527992 2,2107750423 | cysD | sulfate adenylyltransferase s |
| 241 | cds-pgaptmp_-1,212130053 9,2872037616 | NA | hypothetical protein |
| 242 | cds-pgaptmp_-1,390504959 2,6338489649 | NA | YqiA/YcfP family alpha/beta |
| 243 | cds-pgaptmp_-1,136822306 3,8482973787 | rubA | rubredoxin RubA |
| 244 | cds-pgaptmp_-1,674057648 2,0278784043 | NA | carbonic anhydrase |

| | | |
|---|---|---|
| 245 | cds-pgaptmp_1,49093751731,7901937639NA | TonB-dependent siderophore |
| 246 | cds-pgaptmp_1,63650920546,6514581813NA | hypothetical protein |
| 247 | cds-pgaptmp_1,02544340043,0973454183NA | MFS transporter |
| 248 | cds-pgaptmp_1,02472111971,0441281086NA | alpha/beta hydrolase |
| 249 | cds-pgaptmp_-1,467685249 1,3865315012betI | transcriptional regulator BetI |
| 250 | cds-pgaptmp_-1,388220600 6,4477031493betB | betaine-aldehyde dehydroge |
| 251 | cds-pgaptmp_-1,182032965 2,4285226957betA | choline dehydrogenase |
| 252 | cds-pgaptmp_-1,080882575 1,8631672485NA | homocysteine S-methyltrans |
| 253 | cds-pgaptmp_1,66317696111,0284987393NA | hypothetical protein |
| 254 | cds-pgaptmp_1,19833442002,3106769693NA | TonB-dependent siderophore |
| 255 | cds-pgaptmp_-2,276892120 1,9995543872NA | universal stress protein |
| 256 | cds-pgaptmp_1,45028864281,6981902667NA | hypothetical protein |
| 257 | cds-pgaptmp_1,48863608480,0018349370NA | hypothetical protein |
| 258 | cds-pgaptmp_-1,697466268 1,6909542379NA | benzoate/H() symporter Ben |
| 259 | cds-pgaptmp_1,27674610471,0902287933aceI | chlorhexidine efflux PACE tra |
| 260 | cds-pgaptmp_-1,417096808 1,9185308642accC | acetyl-CoA carboxylase bioti |
| 261 | cds-pgaptmp_-1,551558877 1,0908348368accB | acetyl-CoA carboxylase bioti |
| 262 | cds-pgaptmp_-1,089763286 1,0404953594aroQ | type II 3-dehydroquinate deh |
| 263 | cds-pgaptmp_1,00781174577,1583593291NA | ANTAR domain-containing re |
| 264 | exon-pgaptmp1,05773880653,4468997586NA | NA |
| 265 | cds-pgaptmp_1,06865692081,6959207396kynU | kynureninase |
| 266 | cds-pgaptmp_1,27874923314,7134809249NA | amino acid permease |
| 267 | cds-pgaptmp_1,07095490902,2846945817NA | SOS response-associated pe |
| 268 | cds-pgaptmp_-1,102638817 4,3131275150map | type I methionyl aminopeptid |
| 269 | cds-pgaptmp_1,39835498696,5682393196NA | DUF2057 family protein |
| 270 | cds-pgaptmp_-1,330585006 1,1775153110NA | cyd operon YbgE family prot |
| 271 | cds-pgaptmp_-1,500666004 2,0217331480cydX | cytochrome bd-I oxidase sub |
| 272 | cds-pgaptmp_-1,745978750 3,1711628695cydB | cytochrome d ubiquinol oxida |
| 273 | cds-pgaptmp_-2,228599843 7,5194871308NA | cytochrome ubiquinol oxidas |
| 274 | cds-pgaptmp_-2,748784032 2,0589339896cydP | cytochrome oxidase putative |
| 275 | cds-pgaptmp_1,01850339032,5765185990NA | TonB-dependent receptor |
| 276 | cds-pgaptmp_1,38112578241,4610994312NA | 3-oxoacid CoA-transferase s |
| 277 | cds-pgaptmp_1,39447836741,2203154605NA | 3-oxoacid CoA-transferase s |
| 278 | cds-pgaptmp_1,14939416454,9799082156NA | 3-carboxy-cis,cis-muconate |
| 279 | cds-pgaptmp_1,07534988895,1243797795NA | MFS transporter |
| 280 | cds-pgaptmp_-2,225639719 6,9260984047NA | DUF6438 domain-containing |
| 281 | cds-pgaptmp_-1,905322485 3,7979689773NA | MarR family winged helix-tur |
| 282 | cds-pgaptmp_1,20426247376,0681238843NA | 3-oxoacid CoA-transferase s |
| 283 | cds-pgaptmp_-1,182449353 2,3702886373NA | LysR substrate-binding doma |
| 284 | cds-pgaptmp_-1,154610009 1,3598614346NA | MerR family transcriptional re |
| 285 | cds-pgaptmp_1,01373654710,0005685580NA | TetR/AcrR family transcriptio |
| 286 | cds-pgaptmp_-1,096930247 5,0228085539NA | DUF2247 family protein |
| 287 | cds-pgaptmp_-1,400512397 4,4463153863NA | SMI1/KNR4 family protein |
| 288 | cds-pgaptmp_-1,033574291 3,8033708854NA | hypothetical protein |
| 289 | cds-pgaptmp_-1,466419845 5,5840526343NA | DUF2314 domain-containing |
| 290 | cds-pgaptmp_-1,124423371 3,1484170145NA | aromatic amino acid transam |
| 291 | cds-pgaptmp_1,46998515034,1683931175NA | hypothetical protein |
| 292 | cds-pgaptmp_-2,157747954 7,7168066413NA | D-amino acid dehydrogenase |
| 293 | cds-pgaptmp_-1,972085155 3,1131880080alr | alanine racemase |

| | | |
|---|---|---|
| 294 | cds-pgaptmp_-2,548356722 2,8302863286NA | RidA family protein |
| 295 | cds-pgaptmp_1,26933327687,1625050583NA | acyl carrier protein |
| 296 | cds-pgaptmp_-1,239273711 7,3521170734NA | Dyp-type peroxidase |
| 297 | cds-pgaptmp_1,69321878964,4755184483NA | hypothetical protein |
| 298 | cds-pgaptmp_-1,421188638 1,7186391404NA | glutathione peroxidase |
| 299 | cds-pgaptmp_-1,256773887 3,8232770668NA | helix-turn-helix transcriptiona |
| 300 | cds-pgaptmp_-1,258528769 2,2568390715NA | MFS transporter |
| 301 | cds-pgaptmp_-1,054586033 1,6814926353NA | TetR/AcrR family transcriptio |
| 302 | cds-pgaptmp_1,17837420221,1666980355pgaA | poly-beta-1,6 N-acetyl-D-glu |
| 303 | cds-pgaptmp_1,45974504177,5532918746pgaB | poly-beta-1,6-N-acetyl-D-glu |
| 304 | cds-pgaptmp_1,52621633263,9972930815pgaC | poly-beta-1,6-N-acetyl-D-glu |
| 305 | cds-pgaptmp_1,62373723056,3168037659pgaD | poly-beta-1,6-N-acetyl-D-glu |
| 306 | cds-pgaptmp_1,84313298701,1715103908NA | LLM class flavin-dependent |
| 307 | cds-pgaptmp_2,27028257127,7547289627NA | hypothetical protein |
| 308 | cds-pgaptmp_2,04054390771,5961404350NA | glycine zipper domain-contai |
| 309 | cds-pgaptmp_-1,051924773 1,3876249665NA | 3-hydroxyacyl-CoA dehydrog |
| 310 | cds-pgaptmp_1,01999315018,2594776166NA | hypothetical protein |
| 311 | cds-pgaptmp_-1,782197540 1,8586116034NA | hypothetical protein |
| 312 | cds-pgaptmp_-3,641239689 1,2839302645NA | OmpW family outer membra |
| 313 | cds-pgaptmp_1,86872921392,4614463933NA | YMGG-like glycine zipper-co |
| 314 | cds-pgaptmp_2,04873700783,1206843462NA | DNA transfer protein p32 |
| 315 | cds-pgaptmp_1,08248181271,3694744133rpoC | DNA-directed RNA polymera |
| 316 | cds-pgaptmp_1,40990523271,9572419329phoU | phosphate signaling complex |
| 317 | cds-pgaptmp_-1,206547379 9,8501110286cpdA | 3',5'-cyclic-AMP phosphodies |
| 318 | cds-pgaptmp_1,25216363041,8536399539NA | ferrous iron transporter B |
| 319 | cds-pgaptmp_-1,154652470 3,0584418417NA | response regulator |
| 320 | cds-pgaptmp_1,63582914332,8105581255NA | GNAT family N-acetyltransfe |
| 321 | cds-pgaptmp_1,09786233982,8177535934NA | lytic transglycosylase domai |
| 322 | cds-pgaptmp_1,85450576592,1225842886NA | flavodoxin family protein |
| 323 | cds-pgaptmp_2,06094230468,9744252810NA | DUF2938 domain-containing |
| 324 | cds-pgaptmp_1,44370893042,7370348405NA | helix-turn-helix domain-conta |
| 325 | cds-pgaptmp_1,80558449302,1422692607NA | esterase-like activity of phyta |
| 326 | cds-pgaptmp_-1,137585905 3,0102327474NA | OprD family outer membrane |
| 327 | cds-pgaptmp_1,04674834187,5738642182NA | ATP-binding protein |
| 328 | cds-pgaptmp_-1,240558610 4,5289108104NA | heteromeric transposase enc |
| 329 | cds-pgaptmp_-1,108140999 3,7845739464NA | Mu transposase C-terminal d |
| 330 | cds-pgaptmp_-1,158678671 5,5405220054NA | hypothetical protein |
| 331 | cds-pgaptmp_-1,032308178 2,7407765116NA | leucyl aminopeptidase |
| 332 | cds-pgaptmp_2,01779387947,9991110928adeB | multidrug efflux RND transpo |
| 333 | cds-pgaptmp_2,35418999651,3228618317adeA | multidrug efflux RND transpo |
| 334 | cds-pgaptmp_1,77526992511,2659833162NA | zinc-binding dehydrogenase |
| 335 | cds-pgaptmp_1,70200262631,0374343029add | adenosine deaminase |
| 336 | cds-pgaptmp_-1,116918700 7,2969300508raiA | ribosome-associated translat |
| 337 | cds-pgaptmp_1,40010273076,0612424607NA | AzlD domain-containing prot |
| 338 | cds-pgaptmp_1,98481845054,5606754198NA | OmpW family outer membra |
| 339 | cds-pgaptmp_1,45608017606,1983866289NA | MFS transporter |
| 340 | cds-pgaptmp_-1,039157567 5,0832883175NA | Nif3-like dinuclear metal cen |
| 341 | cds-pgaptmp_1,21978418730,0001823349NA | urea amidolyase associated |
| 342 | cds-pgaptmp_1,29430598517,4202560043NA | TetR/AcrR family transcriptio |

| | | |
|---|---|---|
| 343 | cds-pgaptmp_1,09478418121,8507577437NA | hypothetical protein |
| 344 | cds-pgaptmp_-1,251867773 1,8534298965NA | hypothetical protein |
| 345 | cds-pgaptmp_1,00342474048,0286895913NA | LysE/ArgO family amino acid |
| 346 | cds-pgaptmp_1,75257270030,0102134894NA | glutathione S-transferase |
| 347 | cds-pgaptmp_1,22725144961,0558240644NA | TonB-dependent siderophore |
| 348 | cds-pgaptmp_2,32212837481,9122432511NA | hypothetical protein |
| 349 | cds-pgaptmp_2,35396612599,9088389072NA | hypothetical protein |
| 350 | cds-pgaptmp_1,09933502601,1454730622NA | matrixin family metalloprotea |
| 351 | cds-pgaptmp_1,21696478136,7109176922NA | M57 family metalloprotease |
| 352 | cds-pgaptmp_1,00066545989,7485169785NA | putative solute-binding protei |
| 353 | cds-pgaptmp_1,06480497612,7728044811NA | alpha/beta fold hydrolase |
| 354 | cds-pgaptmp_1,53388390728,7845328419NA | SIMPL domain-containing pr |
| 355 | cds-pgaptmp_-1,093255946 4,2133979548NA | thiamine pyrophosphate-binc |
| 356 | cds-pgaptmp_1,51226977666,6357147845NA | substrate-binding domain-co |
| 357 | cds-pgaptmp_1,60793690836,0662920010pstC | phosphate ABC transporter p |
| 358 | cds-pgaptmp_1,60389459092,6419158661pstA | phosphate ABC transporter p |
| 359 | cds-pgaptmp_1,17398928981,1035772059NA | MCR_0457 family protein |
| 360 | cds-pgaptmp_-1,143906569 4,4877798385NA | Lon protease family protein |
| 361 | cds-pgaptmp_1,03383722554,4845352945NA | EAL domain-containing prote |
| 362 | cds-pgaptmp_-1,223716495 2,7780798382NA | AI-2E family transporter |
| 363 | cds-pgaptmp_-1,621715330 8,9839717354NA | DUF3015 family protein |
| 364 | cds-pgaptmp_-1,868116989 1,8519769458basJ | acinetobactin biosynthesis is |
| 365 | cds-pgaptmp_-1,167332541 5,6279170986basI | acinetobactin biosynthesis pl |
| 366 | cds-pgaptmp_-2,311560555 2,1994976051basH | acinetobactin biosynthesis th |
| 367 | cds-pgaptmp_-2,605166329 8,3302255174barB | acinetobactin export ABC tra |
| 368 | cds-pgaptmp_-2,233426852 2,4745340236barA | acinetobactin export ABC tra |
| 369 | cds-pgaptmp_-1,616654314 2,3485948444basG | acinetobactin biosynthesis hi |
| 370 | cds-pgaptmp_1,04322898841,3238168694basE | (2,3-dihydroxybenzoyl)adeny |
| 371 | cds-pgaptmp_-1,409051350 4,1936763376basC | putative histamine N-monoo |
| 372 | cds-pgaptmp_-2,376381953 3,0227462930bauA | TonB-dependent siderophore |
| 373 | cds-pgaptmp_-1,764493901 1,5251528157bauB | siderophore-binding periplas |
| 374 | cds-pgaptmp_1,27920293282,4158772290bauC | ferric acinetobactin ABC tran |
| 375 | cds-pgaptmp_1,89031204523,6891572857bauD | ferric acinetobactin ABC tran |
| 376 | cds-pgaptmp_-1,636477877 3,8887283059basB | acinetobactin non-ribosomal |
| 377 | cds-pgaptmp_-1,070054399 3,1210848724NA | TetR/AcrR family transcriptio |
| 378 | cds-pgaptmp_3,04577029663,1016433780NA | RcnB family protein |
| 379 | cds-pgaptmp_1,77433201937,0048499524NA | RcnB family protein |
| 380 | cds-pgaptmp_1,82621587073,3254041952NA | hypothetical protein |
| 381 | cds-pgaptmp_-1,101801632 3,3294420660NA | FKBP-type peptidyl-prolyl cis |
| 382 | cds-pgaptmp_-1,396975645 2,2638956011NA | acetyl-CoA carboxylase carb |
| 383 | cds-pgaptmp_-1,098385191 6,3065122134NA | organic hydroperoxide resist |
| 384 | cds-pgaptmp_1,41231277575,9313571937NA | GFA family protein |
| 385 | cds-pgaptmp_1,72700544721,1476346999NA | DUF4256 domain-containing |
| 386 | cds-pgaptmp_1,62636309283,4990975774NA | acyl-CoA dehydrogenase far |
| 387 | cds-pgaptmp_1,20818926635,3315474433NA | acyclic terpene utilization Atu |
| 388 | cds-pgaptmp_1,01787597381,2583304510NA | SDR family oxidoreductase |
| 389 | cds-pgaptmp_1,05524766135,0540410359NA | acyl-CoA carboxylase subun |
| 390 | cds-pgaptmp_1,13463764191,3258206872NA | acyl-CoA dehydrogenase far |
| 391 | cds-pgaptmp_-1,261067103 1,7292380888NA | hypothetical protein |

| | | | | | |
|---|---|---|---|---|---|
| 392 | cds-pgaptmp_ | -1,077735021 | 4,2761575931 | NA | ClpXP protease specificity-er |
| 393 | cds-pgaptmp_ | -1,193486443 | 2,7481381836 | NA | LemA family protein |
| 394 | cds-pgaptmp_ | 1,0156527159 | 4,6385825877 | tgt | tRNA guanosine(34) transgly |
| 395 | cds-pgaptmp_ | -1,354206334 | 1,3884099339 | NA | rhodanese-like domain-conta |
| 396 | cds-pgaptmp_ | 2,2508976657 | 4,2459233856 | NA | IS91-like element ISVsa3 far |
| 397 | cds-pgaptmp_ | 1,1362690718 | 2,9657987045 | sul2 | sulfonamide-resistant dihydro |
| 398 | cds-pgaptmp_ | 1,0972712899 | 4,6410049615 | glmM | phosphoglucosamine mutase |
| 399 | cds-pgaptmp_ | 1,8032402581 | 1,4654041038 | NA | IS91-like element ISVsa3 far |

9 hours treatment

9 hours treatment

9 hours treatment

9 hours treatment

9 hours treatment

9 hours treatment

9 hours treatment

9 hours treatment

9 hours treatment
